# SSRI antidepressant citalopram reverses the Warburg effect to inhibit hepatocellular carcinoma by directly targeting GLUT1

**DOI:** 10.1101/2024.07.17.603851

**Authors:** Fangyuan Dong, Kang He, Shan Zhang, Kaiyuan Song, Luju Jiang, LiPeng Hu, Qing Li, Xue-Li Zhang, Naiqi Zhang, Bo-Tai Li, Li-Li Zhu, Jun Li, Mingxuan Feng, Yunchen Gao, Jie Chen, Xiaona Hu, Jiaofeng Wang, Chongyi Jiang, Helen He Zhu, Lin-Tai Da, Jianguang Ji, Zhijun Bao, Shu-Heng Jiang

**Affiliations:** Department of Gastroenterology, Huadong Hospital, Shanghai Medical College, Fudan University, Shanghai 200040, China; State Key Laboratory of Systems Medicine for Cancer, Shanghai Cancer Institute, Ren Ji Hospital, School of Medicine, Shanghai Jiao Tong University, Shanghai 200240, China; Shanghai Key Laboratory of Clinical Geriatric Medicine, Shanghai 200040, China; National Clinical Research Center for Aging and Medicine, Shanghai 200040, China; Department of Geriatrics, Huadong Hospital, Shanghai Medical College, Fudan University Shanghai 200040, China; Department of Liver Surgery, Ren Ji Hospital, School of Medicine, Shanghai Jiao Tong University, Shanghai 200127, China; Key Laboratory of Systems Biomedicine (Ministry of Education), Shanghai Center for Systems Biomedicine, Shanghai Jiao Tong University, Shanghai 200240, China; Center for Primary Health Care Research, Lund University/Region Skåne, Sweden; Shanghai Immune Therapy Institute, Ren Ji Hospital, School of Medicine, Shanghai Jiao Tong University, Shanghai 200127, P.R. China; Shanghai United International School Qingpu Campus, Shanghai, 201799, China; Department of General Surgery, Hepato-biliary-pancreatic Center, Huadong Hospital, Fudan University, Shanghai 200040, China; State Key Laboratory of Systems Medicine for Cancer, Renji-Med-X Stem Cell Research Center, Shanghai Cancer Institute & Department of Urology, Ren Ji Hospital, Shanghai Jiao Tong University School of Medicine, Shanghai, 200127, China

**Author notes:** Correspondence (S.H.J.); (Z.B.) (J.J.). These authors contributed equally to this work (Fangyuan Dong, Kang He, Shan Zhang, Kaiyuan Song, Luju Jiang).

**Keywords:** Drug repurposing, Drug discovery, SLC6A4, SERT, Aerobic glycolysis

## Abstract

Although there is growing appreciation for effective repurposing of selective serotonin reuptake inhibitors (SSRIs) for cancer therapy, particularly hepatocellular carcinoma (HCC), efforts are hampered by limited knowledge of their molecular targets and mechanism of action. Global inverse gene-expression profiling method, drug affinity responsive target stability assay, and in silico molecular docking analysis was performed to identify the targets of SSRIs. Murine subcutaneous, orthotopic models, and patient-derived xenograft were employed to explore the therapeutic effects and underlying mechanisms of SSRIs in HCC. The clinical relevance of SSRI use was verified with real world data. SSRIs exhibit significant anti-HCC effects independent of their known target serotonin reuptake transporter. The glucose transporter 1 (GLUT1) is identified as a new target of SSRIs. Citalopram binds to and antagonizes GLUT1, resulting in reduced glycolytic flux and ATP generation. Mutant GLUT1 in the binding site E380 of citalopram compromises the inhibitory effects of citalopram on the Warburg effect and tumor growth. In preclinical models, citalopram dampens the growth kinetics of GLUT1^high^ liver tumors and displays a synergistic effect with anti-PD-1 therapy. Retrospective analysis of health records found that SSRIs use is associated with a lower risk of metastasis among HCC patients. Our study reveals an unprecedented role of SSRIs in cancer metabolism, and establishes a rationale for repurposing SSRIs as potential anticancer drugs for HCC.

## Introduction

Serotonin or 5-hydroxytryptamine (5-HT) is a known monoamine neurotransmitter critical for neuronal processes such as mood, behavior, sleep, cognition, and anxiety ^1, 2^. The serotonin reuptake transporter (SERT) removes serotonin from extracellular space, thereby terminating 5-HT signaling ^3^. Selective serotonin reuptake inhibitors (SSRIs), such as citalopram, escitalopram, fluoxetine, fluvoxamine, paroxetine, and sertraline, block SERT function with high affinity and are the most prescribed antidepressants for treating depression and anxiety disorders ^4^. The drug repurposing strategy has emerged as a useful approach to identifying new uses for already approved drugs ^5^. Given that major depressive disorder is relatively common in cancer patients, repurposing SSRIs as potential anticancer therapeutics represents a time-saving and economic alternative, benefitting from available drug formulations and clinical evidence of safety and toxicity profile.

Accumulating epidemiological studies reveal that SSRIs use has a lower risk of death from various cancer types, such as liver cancer, bladder cancer, kidney cancer, breast cancer, endometrial cancer, and ovarian cancer ^6–11^. Recently, our group also showed that SSRIs monotherapy was inversely associated with colorectal cancer risk, and the combined use of aspirin and SSRIs showed a synergistic effect ^12^. Consistent with clinical observations, data from *in vitro* cell experiments and preclinical models suggest that SSRIs exert substantial anti-tumor effects, including but not limited to cell senescence, cell apoptosis, invasive potential, and drug resistance ^13–16^. However, there is little consensus on the molecular mechanisms of the antineoplastic roles of SSRIs, as both SERT-dependent and SERT- independent actions have been proposed ^17–21^. For instance, targeting SERT with sertraline in colon cancer induces a compensatory effect on serotonin biosynthesis and oncogenic kynurenine production, whereas fluoxetine potently inhibits SMPD1 activity to suppress glioblastoma development ^17, 20^. Fluoxetine can also trigger long-term control of colorectal and pancreatic tumors via serotonylation-related mechanism ^21^. Therefore, further exploration of cancer-specific mechanisms is needed to repurpose SSRIs for cancer therapy.

Hepatocellular carcinoma (HCC), as the major subtype of liver cancer, is the third leading cause of cancer-related mortality worldwide ^22^. Despite substantial progress achieved in HCC treatment, including immune checkpoint inhibitor (ICI)-based therapies, most patients are likely to not respond and eventually succumb to HCC ^23^. Therefore, more effective drugs are still required to enable personalized and cost-effective treatment. A meta-analysis of eight observational studies with one million participants revealed that SSRIs use is associated with a 34% lower risk of HCC ^24^. These facts prompt us to test the possibility and rationality of repurposing SSRIs for HCC treatment.

In this study, we reveal a wide spectrum of anti-tumor effects of SSRIs in HCC. Interestingly, loss-of-function studies show that the anticancer activities of SSRIs are independent of SERT expression. Using the global inverse gene-expression profiling method, drug affinity responsive target stability (DARTS) assay, and *in silico* molecular docking analysis, the glucose transporter 1 (GLUT1) is identified as a new target of SSRIs in HCC. Through binding GLUT1 E380, citalopram blunts the glycolytic metabolism in HCC cells.

## MATERIALS AND METHODS

### Study participants

All patients who were diagnosed with HCC between January 2006 and December 2018 were identified from the Swedish Cancer Registry by using the 10th International Classification of Disease (ICD-10) code (C22.0). The TNM staging system, which includes the size of tumor (T), nodal status (N), and the presence of metastatic disease (M), was used to define the stage at diagnosis of HCC as stage I (Tis/T1 N0 M0), stage II (T2 N0 M0), stage III (T3/T4 N0 M0), and stage IV (Any T N1 M0, any T or N M1). Prescriptions of SSRI were identified by linking to the Swedish Prescribed Drug Register using Anatomical Therapeutic Chemical (ATC) code N06AB. The outcome was cancer metastasis (ICD 10: C77-C79). The information on the date of death and cause of death was collected from the Cause of Death Register. The follow-up started at the date of diagnosis with HCC and ended at the date of metastasis, date of death, or at the end of the follow-up period (December 2018), whichever came first. Patients with follow-up time of fewer than three months were excluded from the study population. Metastasis-free survival was computed using the Kaplan–Meier method. Log-rank test was conducted to examine the difference in survival probability between SSRI users and non-users. Time-dependent Cox regression was used to calculate hazard ratios (HRs) and 95% confidence intervals (CIs) of metastasis associated with the use of SSRI. SSRI use was modeled as a time-dependent variable. Death from any causes was computed as a competing event in order to control the competing risk of death.

### Statistical analysis

Statistical methods were not employed to predetermine sample size. The size of *in vitro* and *in vivo* studies, based on similar studies in the field, was validated by pilot studies. Sample sizes and biological replicates are included in the indicated figure legends. All results for continuous variables are presented as means ± SD from at least three technical replicates. When comparing two independent groups, a two-tailed Student’s t-test was used. Statistical analyses for more than two groups were performed with a one-way analysis of variance (ANOVA) with Tukey’s method or two-way ANOVA with Dunnett’s multiple comparisons as indicated. GraphPad Prism 9 (GraphPad Software Inc., San Diego, CA) was used for statistical analyses. Statistical correlation in human HCC was determined by nonparametric Spearman correlation analysis. Kaplan-Meier survival curves were determined by the log-rank test. A customary threshold of p-value less than 0.05 was considered statistically significant. S Statistical significance in all Figures is denoted as follows: *p < 0.05; **p < 0.01; ***p < 0.001.

## Results

### The anti-tumor effects of SSRIs in HCC

To investigate the potential anti-tumor effects of SSRIs, we treated two HCC cell lines, Huh7 and HCC-LM3, which exhibit high endogenous SERT, with six commonly prescribed SSRIs (citalopram, escitalopram, fluoxetine, fluvoxamine, paroxetine, and sertraline) (Figures S1A and S1B). SSRIs treatment yielded pronounced cytotoxic activities against HCC cells but not the nonmalignant LO-2 cells (Figure S1C). The impacts of SSRIs on HCC cells were examined phenotypically by colony formation assay and Caspase-3/7 assay. As a result, SSRIs led to an approximately 50% decline in the colony-forming ability of HCC cells (Figure 1A) and caused a 1.5 to 2-fold increase in Caspase-3/7 activity under serum-starvation condition (Figure 1B). To evaluate the *in vivo* effects of SSRIs, HCC cells were subcutaneously implanted into the flanks of immune-deficient nude mice, and SSRIs were given by i.p. administration daily for 30 days after bearing palpable tumors (∼50 mm^3^). All 6 SSRIs effectively retarded tumor growth (Figure 1C) and fundamentally prolonged mouse lifespan (Figure 1D), suggesting a class-specific effect. Citalopram was selected for further study because of its remarkable impact. Moreover, we generated orthotopic tumor model with mouse Hepa1-6 in immune-competent C57BL/6 mice. In parallel with its *in vitro* anti-tumor activities (Figures S1D-G), citalopram remarkably delayed tumor growth and even induced complete remissions in a tumor-bearing mouse (Figure 1E). The *in vivo* anti-tumor effects of SSRIs were confirmed at the molecular levels as evidenced by a decrease in the proliferation index Ki-67 and an increase in the apoptosis marker cleaved caspase-3 (CCS3) (Figure S2).

**Fig. 1.**
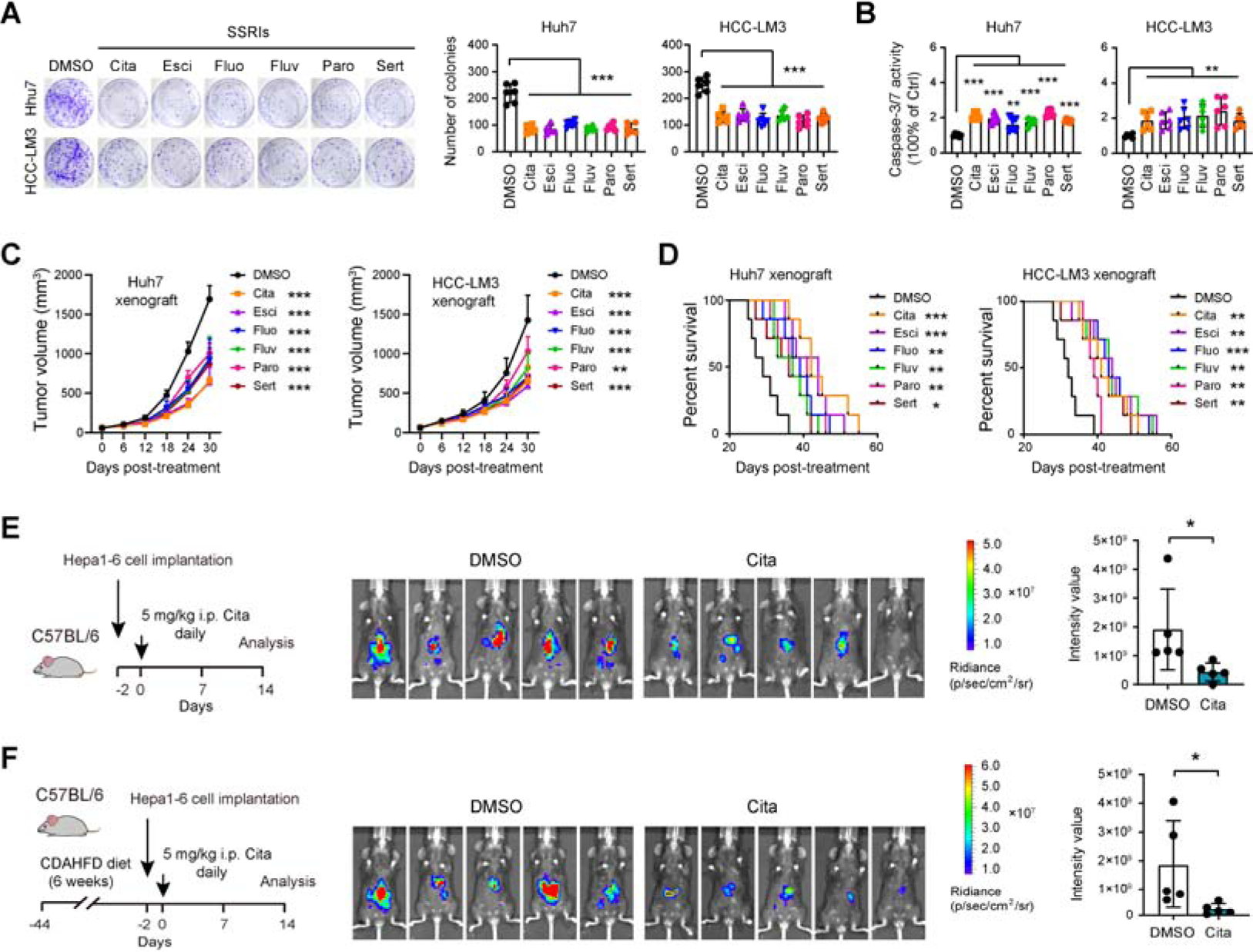
SSRIs exhibit wide anti-tumor effects in HCC. (A) Plate colony formation assay showed the effect of SSRIs (5 μM) on the long-term cell proliferation of Huh7 and HCC-LM3 cells (n = 6 per group). (B) Caspase-3/7 activity in Huh7 and HCC-LM3 cells upon treatment with 5 μM SSRIs for 48 h (n = 6 per group). (C) Huh7 and HCC-LM3 cells were subcutaneously injected into the nude mice, and mice in drug groups were subjected to treatment with 5 mg/kg SSRIs when bore visible tumors (∼50 mm^3^); 30 days later, tumor burden in the control group and six SSRI groups was detected (n = 5 per group). (D) Survival curve of mice engrafted with Huh7 and HCC-LM3 cells in the control group and six different SSRI groups (n = 7 per group). Mice were sacrificed when tumors were larger than 15 mm. (**E** and **F**) In orthotopic xenograft model, which generated with Hepa1-6 cells in immunocompetent C57BL/6 mice or MASH mice (n = 5 per group), the tumor burden after citalopram treatment was monitored by an *in vivo* imaging system. In all panels, *p < 0.05, **p < 0.01, ***p < 0.001. Values as mean ± SD and compared by one-way ANOVA multiple comparisons with Tukey’s method (**A**, **B**), two-way ANOVA with Dunnett’s multiple comparisons (**C**), the log-rank test (**D**), and Student’s t test (**E**, **F**).

Metabolic dysfunction-associated fatty liver disease (MAFLD) is a major driver of HCC ^25,26^. To reflect the MAFLD etiology of HCC, we fed C57BL/6 mice with a choline-deficient, amino acid-defined high-fat diet (CDAHFD) to promote metabolic dysfunction-associated steatohepatitis (MASH)/fibrosis before orthotopic Hepa1-6 cell implantation (Figures S3A and S3B). Likewise, citalopram substantially reduced tumor growth in mice with MASH (Figure 1E). Moreover, histological analysis revealed lesser proliferating cells and more apoptotic cells upon citalopram treatment (Figure S3C).

Given that serotonin is transported by SERT to the liver for degradation and abundant SERT can be detected in the liver tissues (Figure S3D), we tested the possible side effects of SSRIs. For clinical relevancy, a 5_mg/kg/day dose of citalopram for a mouse is equivalent to 25_mg/day in humans, which is within the dose range of citalopram (10-60_mg/day) for treating depression ^27^. The dose escalation study showed that C57BL/6 mice were well- tolerated to a 2-20_mg/kg/day dose of citalopram treatment, which did not have any effect on the mouse body weight or liver injury-associated markers, alanine transaminase (ALT) and aspartate aminotransferase (AST) (Figure S3E). Particularly, a high dose of citalopram (20_mg/kg/day) caused mice to become irritable and fight frequently in the first week of treatment, but mice later returned to the normal state, suggesting the transient side effects of a high dose of SSRIs on the nervous system. Additionally, the tumor-bearing mice with MASH demonstrated tolerance and survival when subjected to a 2-20_mg/kg/day dose of citalopram treatment for 2 weeks. Administration of citalopram resulted in decreased liver weight, body weight, liver-to-body weight ratio, as well as reduced levels of ALT and AST, indicating that the anti-cancer properties of citalopram may contribute to the alleviation of liver injury (Figure S3F).

### Citalopram-induced anti-tumor effects are largely independent of SERT

5-HT uptake by SERT engages tissue transglutaminase-mediated serotonylation, a post- translational modification implicated in many physiological and pathophysiological processes ^28^. As SERT is the known target of SSRIs, we wondered whether the effect of SSRIs can be phenocopied by SERT knockdown in HCC. Two specific shRNAs against *SLC6A4* led to marked reduction of SERT protein level in Huh7 and HCC-LM3 cells (Figure 2A). Although 5- HT had no effects on the proliferation of HCC cells cultured with a complete medium, it protected HCC cells from apoptosis caused by serum starvation (Figures 2B and 2C). In the presence of 5-HT, SERT silencing resulted in only a minimal reduction in HCC cell proliferation (Figure 2B) and had negligible impact on Caspase-3/7 activity (Figure 2C). Considering that residual SERT in the shRNA strategy may still be effective for function, we further knocked out SERT using CRISPR/Cas9 method (Figure 2D). Compared to vector cells, SERT^-/-^ cells had a ∼10% reduction in cell proliferation (Figure 2E) and no discernable increase in cell apoptosis (Figure 2F). Importantly, citalopram sufficiently suppressed the colony-forming ability (Figures 2B and 2E) and increased Caspase-3/7 activity in SERT knockdown or SERT^-/-^ cells (Figures 2C and 2F). In alignment with this, citalopram exerted notable anti-proliferation and pro-apoptosis effects in a cell line with lower SERT expression (Figures S1B, S4A, and S4B). In a mouse colon carcinoma cell line (MC38) with very high level of SERT, the anti-tumor effects of citalopram can be noticed even in the presence of SERT knockdown (Figures S4C-E). *In vivo* investigation with subcutaneous xenograft model further recapitulated the above *in vitro* therapy responses of citalopram as confirmed by reduced tumor burden, increased mouse lifespan, less cell proliferation, and elevated cell apoptosis (Figures 2G, 2H, S5A, and S5B). Specifically, SERT expression remained unaltered upon citalopram treatment (Figure S5C). Together, these data suggest that SSRI might target other molecules instead of SERT to exhibit its anti-tumor effects in HCC.

**Fig. 2.**
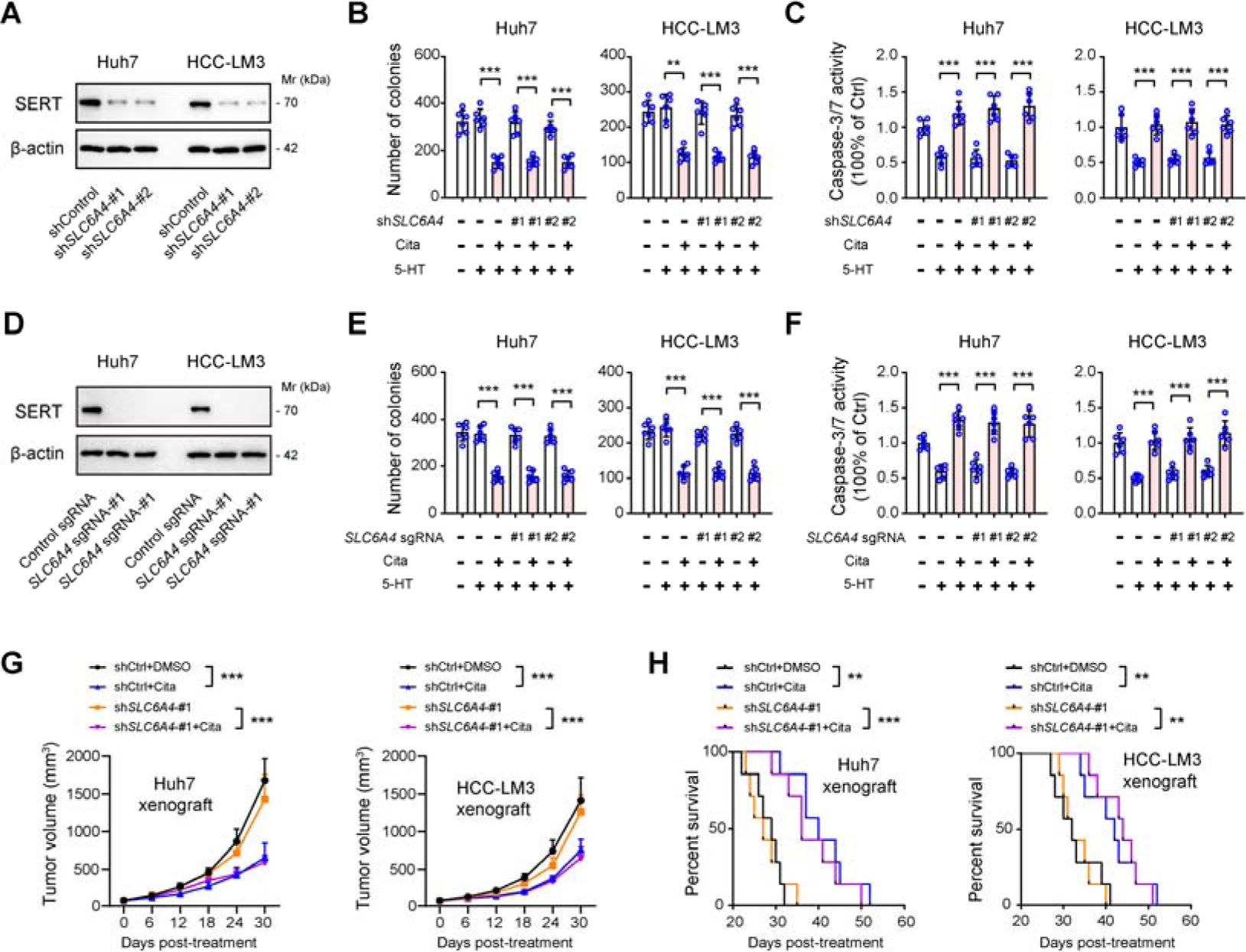
Citalopram-induced anti-tumor effects are largely independent of its known target SERT. (A) Two specific shRNAs were used for the genetic silencing of SERT in Huh7 and HCC-LM3 cells, and the knockdown efficiency was verified by Western blotting analysis. (B) Plate colony formation assay revealed the effect of SERT knockdown alone or combined with citalopram treatment on the long-term cell proliferation of Huh7 and HCC-LM3 cells in the presence of 50 μM 5-HT (n = 6 per group). (C) Caspase-3/7 activity in Huh7 and HCC-LM3 cells upon SERT knockdown or combined treatment with 5 μM citalopram in the presence of 50 μM 5-HT (n = 6 per group). (D) Western blotting showed SERT protein levels in sgControl, *SLC6A4* sgRNA-#1, and *SLC6A4* sgRNA-#2 Huh7 and HCC-LM3 subclones. (E) Plate colony formation assay revealed the effect of SERT knockout alone or combined with citalopram treatment on the long-term cell proliferation of Huh7 and HCC-LM3 cells in the presence of 50 μM 5-HT (n = 6 per group). (F) Caspase-3/7 activity in Huh7 and HCC-LM3 cells upon SERT knockout or combined treatment with 5 μM citalopram in the presence of 50 μM 5-HT (n = 6 per group). (G) shControl and sh*SLC6A4*-#1 Huh7 and HCC-LM3 cells were subcutaneously injected into the nude mice, and mice in the combined treatment group were treated with 5 mg/kg citalopram when bore visible tumors (∼50 mm^3^); 30 days later, tumor burden was examined (n = 5 per group). (H) Survival curve of mice engrafted with shControl and sh*SLC6A4*-#1 Huh7 and HCC-LM3 cells in the presence or absence of treatment with 5 mg/kg citalopram (n = 7 per group). In all panels, *p < 0.05, **p < 0.01, ***p < 0.001. Values as mean ± SD and compared by Student’s t test (**B**, **C**, **E**, **F**), two-way ANOVA with Dunnett’s multiple comparisons (**G**), and the log-rank test (**H**).

### Identification of molecular target of SSRIs in HCC

To predict the possible targets of SSRIs, we searched the SuperPred database ^29^. Apart from SERT, 21 other targets (ADAM10, APEX2, C5, C5aR1, CTSD, DUSP3, GLRA1, GRIN1, GRIN2B, HRH3, KLF5, LSD1, MAOB, NFE2L2, NFKB1, NR1I2, NTRK3, PRCP, SLC2A1, SLC6A5, and TRIM24) were predicted to be targeted by all SSRIs (Figure 3A and Tables S1- 5). Among these candidates, 15 of them can be detected in HCC tissues and subjected to further analysis (Figure S6A).

**Fig. 3.**
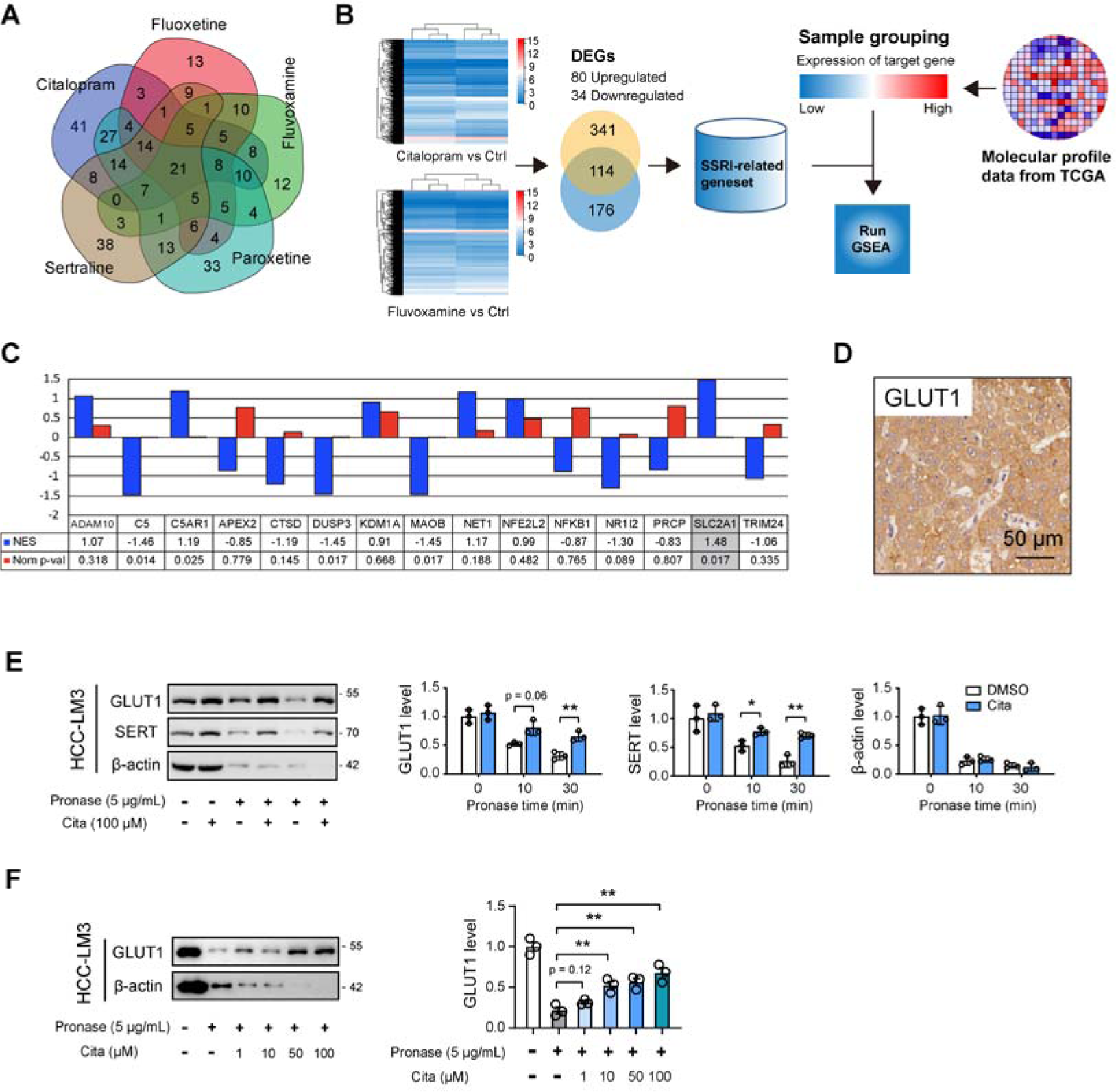
Identification of potential molecular targets of SSRIs in HCC. (A) Venn diagram showed the predicted targets of SSRIs. Data were acquired from the SuperPred database. (B) Strategy for selecting candidate targets of SSRIs in HCC. (C) Gene set enrichment analysis (GSEA) with SSRI-related gene set and RNA-seq data of HCC sample (TCGA cohort) showed enriched genes related to SSRI treatment. NES, normalized enrichment score. (D) Representative immunohistochemical images showed the expression pattern and cellular distribution of GLUT1 in human HCC tissues. Scale bar, 50 μm. (E) The DARTS assay and immunoblot analysis showed GLUT1 and SERT protein stability against 5 μg/mL pronase in the presence and absence of 100 μM citalopram treatment. Two time points (10 min and 30 min) for pronase treatment were used. (F) The DARTS assay and immunoblot analysis showed GLUT1 protein stability against 5 μg/mL pronase in the presence of different concentration of citalopram treatment (0, 1, 10, 50, and 100 μM). In all panels, *p < 0.05, **p < 0.01. Values as mean ± SD and compared by the Student’s t test (**E**) or one-way ANOVA multiple comparisons with Tukey’s method (**F**).

Drug-induced gene signature networks that invert cancer profiles together with computational tools have been widely exploited for drug repurposing ^5, 30^. Using this strategy, we investigated whether SSRI-induced gene expression profiles are inversely matched with the candidate target-associated signaling network. To this end, we first performed RNA sequencing analysis in HCC-LM3 cells upon citalopram or fluvoxamine treatment and identified 80 co-upregulated and 34 co-downregulated genes (Figure 3B and Table S6); these differentially expressed genes (DEGs) were selected for generation of an SSRI-related gene signature. Secondly, we mined the RNAseq data of the liver hepatocellular carcinoma (LIHC) samples (n = 371) in The Cancer Genome Atlas (TCGA) cohort and divided them into high and low groups based on the median expression value of each candidate target. Finally, grouped samples were subjected to Gene Set Enrichment Analysis (GSEA) with the SSRI- related gene signature (Figure 3B). As a result, SLC2A1, also known as glucose transporter 1 (GLUT1), was revealed to be the top-scoring hit negatively correlated with SSRI-related gene profiles in HCC (Figures 3C and S6B). In HCC samples, GLUT1 was largely expressed by cancer cells (Figures 3D).

DARTS is a well-documented approach to identifying potential protein targets for small molecules that confers proteolytic protection of target proteins by interacting with small molecules ^31^. Combined DARTS assay with immunoblotting analysis, we tested GLUT1 stability against pronase in the presence or absence of citalopram treatment. Citalopram did not affect the basal protein and mRNA level of GLUT1 in HCC cells (Figures 3E and S6C). Citalopram can increase GLUT1 stability against pronase, but it had no effect on the proteolytic susceptibility of ACTB (Figure 3E). For validation, SERT was also examined (Figure 3E). Moreover, the stability of GLUT1 against pronase was increased in a dose- dependent manner by citalopram treatment, while no change in β-actin stability was observed, demonstrating that citalopram specifically binds to GLUT1 (Figures 3F). Besides citalopram, other 5 SSRIs also played protective roles of GLUT1 against pronase (Figure S6D).

### Molecular docking reveals GLUT1 as direct target of citalopram

To further address the issue of SSRI/GLUT1 interaction, *in silico* docking analysis was conducted. Citalopram was docked into the substrate-binding site based on the crystal structure of human GLUT1 (PDB ID: 4PYP) in the inward-facing conformation ^32^. By clustering 500 predicted binding poses of citalopram, we found that the top 2 clusters exhibited relatively low binding free energy (Figure 4A). The overall conformation showed that both the predicted poses with the lowest binding free energy from cluster 1 (pose-1) and cluster 2 (pose-2), to some extent, occupied the substrate-binding site of GLUT1 (Figure 4B). In detail, the fluorophenyl group of citalopram in pose-1 established hydrophobic interactions with V328, P385, and L325; the cyanophtalane group stuck into the substrate-binding cavity of GLUT1, while the two groups in pose-2 bind in the opposite orientation (Figure 4C, left). Intriguingly, the carboxylate group of residue E380 interacted with the amino group of citalopram in both poses (Figure 4C, middle and right), implying the importance of the electrostatic interactions between E380 and the amino group of citalopram. Considering the fact that E380 was involved in both two poses, we further examined the functional role of E380 in the binding of citalopram by generating a GLUT1 E380A mutant using the site- directed mutagenesis assay (Figure 4D). For this purpose, the WT and E380A mutant of GLUT1 expression plasmids were transfected into sh*SLC2A1*-expressing (GLUT1^KD^) HCC- LM3 cells. The DARTS assay with immunoblotting showed increased stability of GLUT1 with the transfection of WT HA-GLUT1 against pronase after citalopram treatment, whereas the proteolytic susceptibility of GLUT1 with the transfection of HA-GLUT1 E380A showed no changes upon citalopram treatment (Figure 4E). Because SSRIs have different structures, we tested whether the binding model is restricted to citalopram. It is noteworthy that other SSRIs contain an amino group that was predicted to establish electrostatic interactions with GLUT1 E380 (Figure 4F) and this finding was further validated by DARTS assay (Figure 4G).

**Fig. 4.**
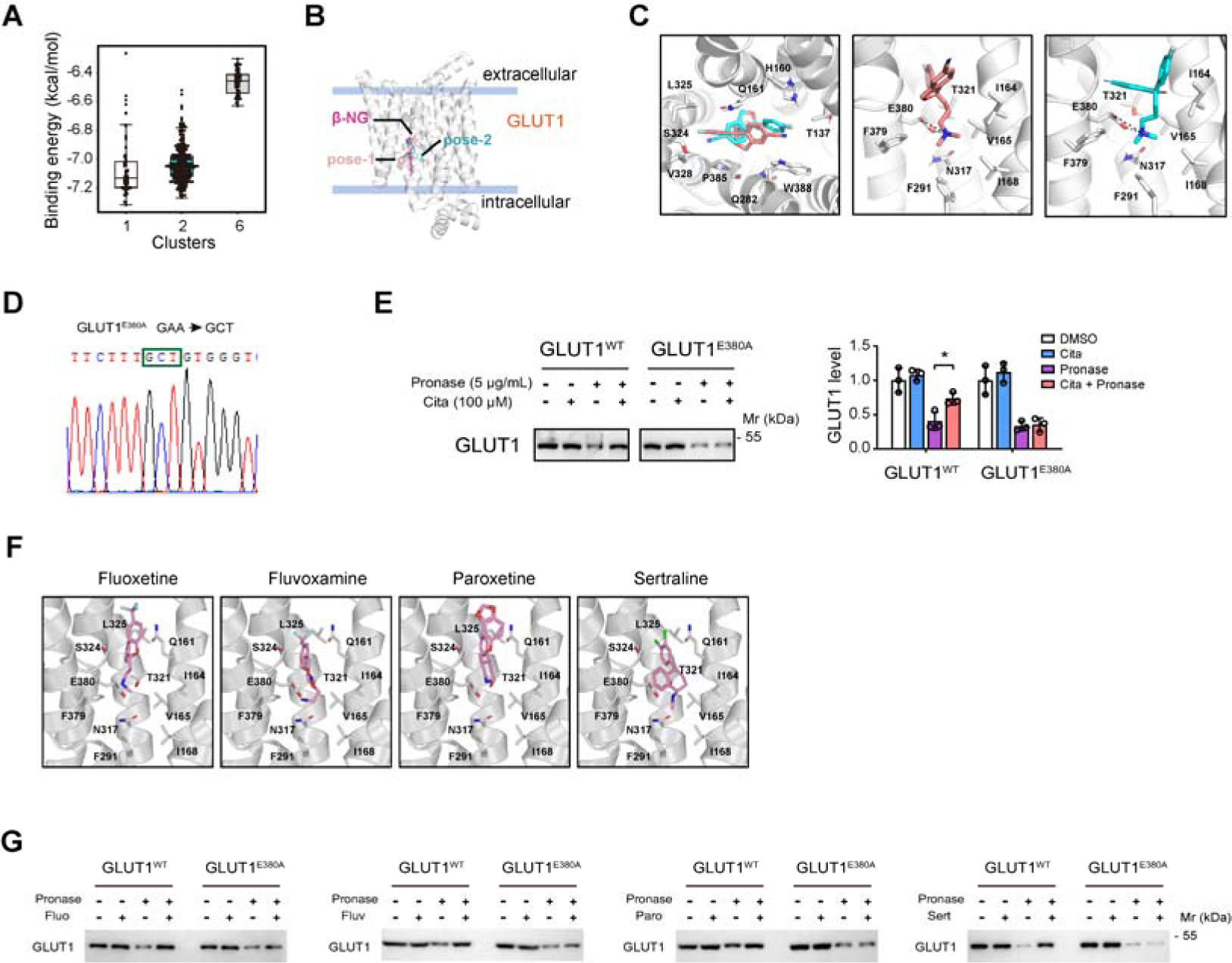
Molecular docking reveals GLUT1 as a direct target of citalopram. (A) For GLUT1, predicted binding energy distribution of the clusters with poses more than 50. (B) The overall conformation of citalopram binding to GLUT1. The best poses from cluster 1 (pose-1) and cluster 2 (pose-2) were shown in salmon and cyan sticks, respectively. The structure of human GLUT1 (PDB ID: 4PYP) was shown in a white cartoon with one substrate β-NG (magenta sticks) bound. (C) The details of citalopram binding to GLUT1; left, the overlapping conformation of pose-1 and pose-2; middle and right, the interactions between citalopram and the amino group of GLUT1 for pose-1 (middle) and pose-2 (right). (D) Sequencing analysis showed the successful generation of GLUT1 E380A mutant. (E) GLUT1-dificient HCC-LM3 cells were transfected with WT and E380A mutant of GLUT1 expression plasmids for 48 h. And then, the cells were harvested, and lysed, followed by DARTS assay with immunoblotting analysis of GLUT1 protein levels. (F) The best-scored complex models of GLUT1 with different SSRIs (fluoxetine, fluvoxamine, paroxetine, and sertraline). (G) GLUT1^KD^ HCC-LM3 cells were transfected with WT and E380A mutant of GLUT1 expression plasmids for 48 h. Then, the cells were harvested, lysed, and treated with indicated SSRIs, followed by DARTS assay with immunoblotting analysis of GLUT1 protein levels. In all panels, *p < 0.05, **p < 0.01. Values as mean ± SD and compared by the Student’s t test (**E**).

### Citalopram inhibits the Warburg effect by targeting GLUT1

Since GLUT1 is responsible for transporting glucose into the cytoplasm and enables the Warburg effect in cancer, we sought to probe the effects of SSRIs on glucose metabolism. Expectedly, glucose uptake, lactate release, and ATP generation were significantly diminished by SSRIs treatment (Figures 5A-C and S7A-C). Concurrently, Seahorse analysis demonstrated that SSRIs attenuated the extracellular acidification rate (ECAR) in HCC cells (Figures 5D and S7D). To uncover whether SSRIs-induced impairment of glycolysis was GLUT1-dependent, we genetically silenced GLUT1 in HCC cells and found that GLUT1 knockdown reduced glycolytic flux to a similar extent as citalopram did, reflected by glucose uptake, lactate release, ATP levels, and ECAR (Figures 5E-I). No additive effects of citalopram on the glycolytic flux in GLUT1 knockdown cells were noticed. Enforced expression of GLUT1^WT^ in endogenous GLUT1-deficient HCC-LM3 cells potentiated glycolytic flux and ATP generation, which can be largely compromised by the addition of citalopram (Figures S7E-I). In contrast, citalopram cannot further abrogate the glycolytic metabolism in cells expressing the mutant version of GLUT1 (GLUT1^E380A^) (Figures S7J-N). Citalopram did not affect the mRNA levels of glycolytic genes (*SLC2A1*, *HK2*, *PKM2* and *LDHA*) or the protein levels of GLUT1 (Figures S6D and S7O), suggesting that the distinct effects of SSRIs on glycolysis are due to the direct inhibition of SSRIs on GLUT1, rather than the SSRIs- mediated decrease in the expression of glycolytic genes.

**Fig. 5.**
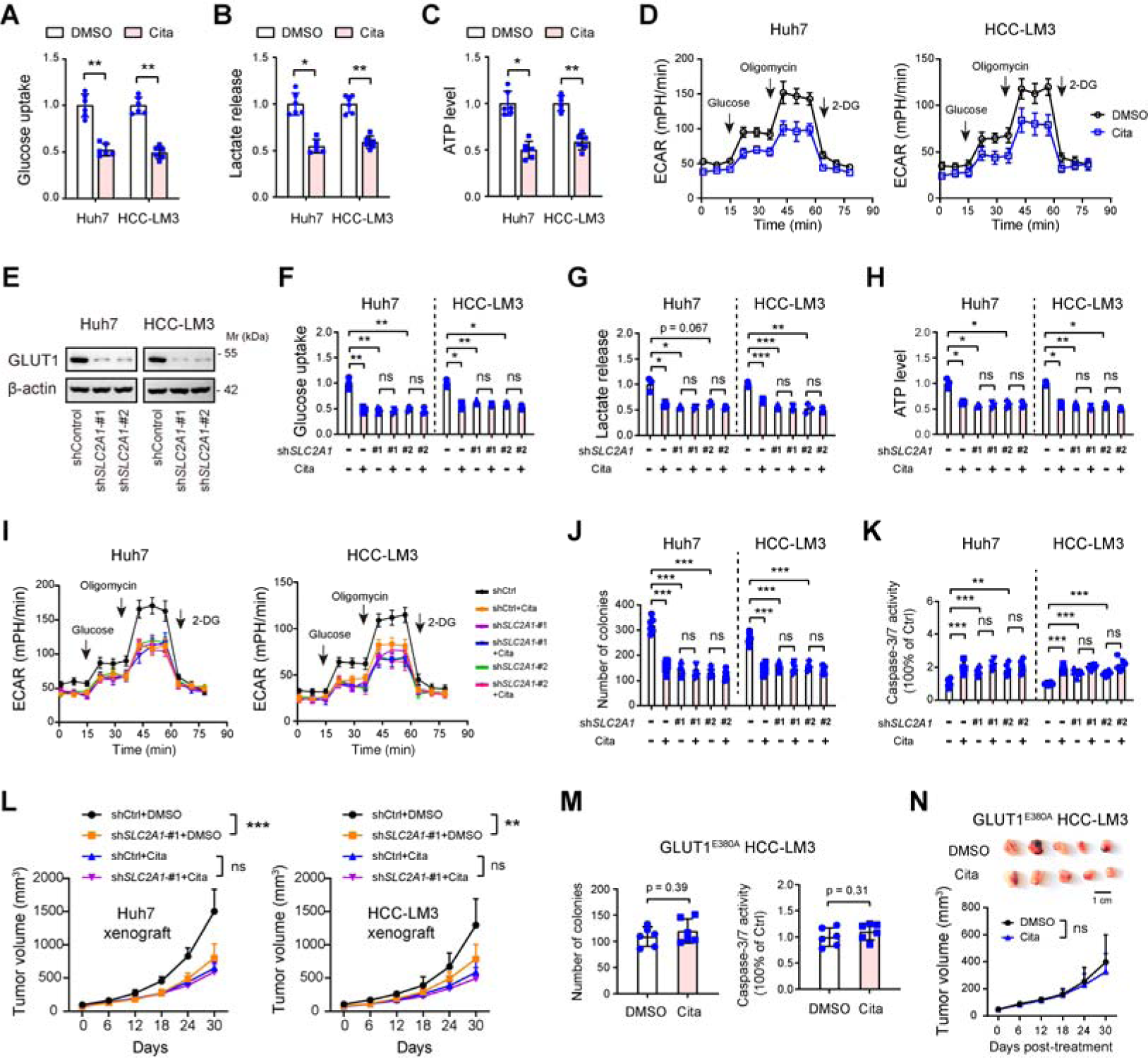
Citalopram inhibits the Warburg effect by targeting GLUT1 on HCC cells. (**A-D**) The effects of SSRIs on the glycolytic ability of Huh7 and HCC-LM3 cells were measured, as indicated by glucose uptake, lactate production, ATP generation (**A**-**C**, n = 6 per group), and extracellular acidification rate (ECAR) (**D**, n = 3 per group). (**E**) Western blotting showed the knockdown efficiency of GLUT1 in Huh7 and HCC-LM3 cells. (**F-I**) The effects of GLUT1 knockdown alone or combined with citalopram treatment on the glycolytic ability of Huh7 and HCC-LM3 cells were measured, as indicated by glucose uptake (**F**), lactate production (**G**), ATP generation (**H**), and ECAR (**I**, n = 3 per group). (J) Plate colony formation assay revealed the effect of GLUT1 knockdown alone or combined with citalopram treatment on the long-term cell proliferation of Huh7 and HCC-LM3 cells (n = 6 per group). (K) Caspase-3/7 activity in Huh7 and HCC-LM3 cells upon GLUT1 knockdown or combined treatment with 5 μM citalopram (n = 6 per group). (L) shControl and sh*SLC2A1*-#1 Huh7 and HCC-LM3 cells were subcutaneously injected into the nude mice, and mice in combined treatment group were treated with 5 mg/kg citalopram when bore visible tumors; 30 days later, tumor burden was examined (n = 5 per group). (M) The effects of citalopram on the cell proliferation and cell apoptosis of GLUT1^E380A^ HCC- LM3 cells (n = 6 per group). (N) Effects of citalopram on the growth of GLUT1^E380A^ HCC-LM3 tumors (n = 5 per group). In all panels, *p < 0.05, **p < 0.01, ***p < 0.001; ns, non-significant. Values as mean ± SD and compared by the Student’s t test (**A**-**C**, **M**, **N**), one-way ANOVA multiple comparisons with Tukey’s method (**F**-**H**, **J**, **K**), and two-way ANOVA with Dunnett’s multiple comparisons (**L**, **N**).

Increased glycolysis supports the rapid proliferation of cancer cells by providing additional energy and substrates of biosynthesis ^33^. In parallel with reduced glycolysis, GLUT1 knockdown inhibited *in vitro* cell proliferation, increased cell apoptosis, and *in vivo* tumor growth, and no significant effects were found upon addition of citalopram treatment (Figures 5J-L). Moreover, the growth inhibition and cell apoptosis induced by citalopram were not found in cells only expressing GLUT1^E380A^ (Figures 5M and 5N), pinpointing that the inhibitory effects of citalopram can be attributed to this binding. SSRIs have been reported to inhibit β-catenin, thereby decreasing tumor growth ^34^. However, in silico docking analysis revealed that SSRIs were less likely interacting with β-catenin (Table S7), arguing that SSRIs act downstream of or in parallel to β-catenin/TCF-driven mechanism rather than directly inhibiting β-catenin.

### Citalopram is prominently effective in highly glycolytic liver tumors and potentiates anti-PD-1 therapy

To aid the clinical relevance of SSRIs for HCC treatment, we first tested the therapeutic potential of SSRIs in the patient-derived xenograft (PDX) model. For this purpose, HCC patients who received preoperative ^18^F-Fluorodeoxyglucose (^18^F-FDG) PET/CT were enrolled (Figure 6A). Two PDXs that greatly differ in GLUT1 expression were selected (Figure 6B). Indeed, HCC samples with intense staining of GLUT1 had a higher maximum standard uptake value (SUVmax), while GLUT1-negative HCC showed a lower SUVmax value (Figure 6C). Correspondingly, GLUT1^high^ xenograft tumors grew faster than GLUT1^low^ xenograft tumors (Figure 6D). Citalopram treatment markedly restrained the growth kinetics of GLUT1^high^ tumors, while had limited growth-inhibitory effect on GLUT1^low^ tumors; as a positive control, the glycolysis blocker 2-deoxy-D-glucose (2-DG) also retarded tumor growth as citalopram did in GLUT1^high^ PDX but not GLUT1^low^ PDX (Figure 6D). The suppressive effects of citalopram or 2-DG in the PDX model were further supported by the staining of Ki67 and CCS3 in tumor tissues (Figure S8A). Consistently, citalopram or 2-DG treatment was beneficial to the survival of mice bearing GLUT1^high^ tumors (Figure 6E).

**Fig. 6.**
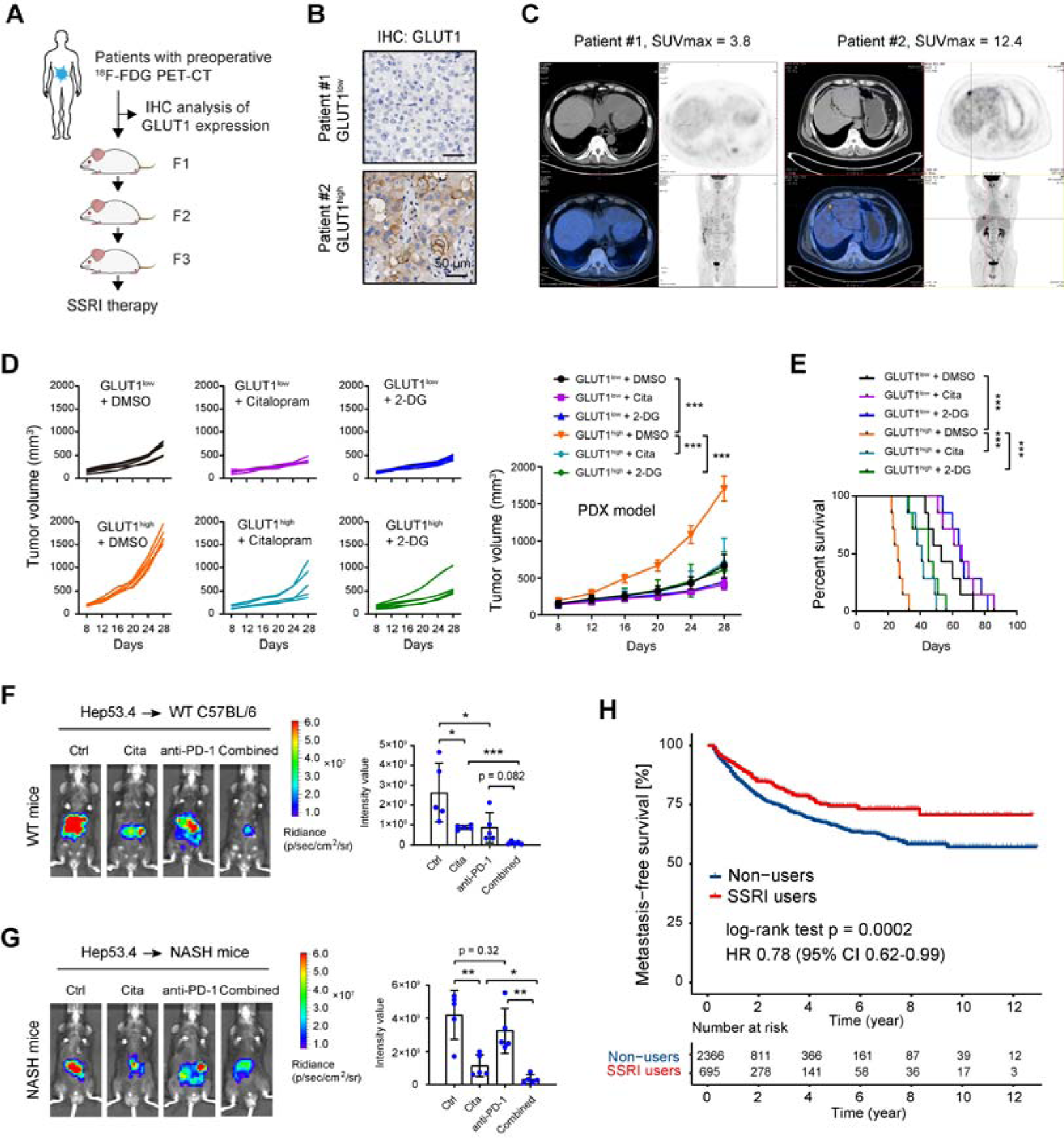
Citalopram is prominently effective in highly glycolytic liver tumors and potentiate anti-PD-1 therapy. (A) Schematic depicting the strategy of patient-derived xenograft (PDX) investigation. (B) GLUT1 expression in two PDX models (patient tissues) was determined by immunohistochemical analysis. Scale bar, 50 μm. (C) PET/CT images showed^18^F-Fluorodeoxyglucose (^18^F-FDG) uptake from two HCC cases used for the PDX study. (D) The growth curve of GLUT1^low^ and GLUT1^high^ tumors in the presence or absence of citalopram treatment and 2-DG treatment. (E) Mouse lifespan in GLUT1^low^ and GLUT1^high^ PDX model upon citalopram or 2-DG treatment. (**F** and **G**) Citalopram and anti-PD-1 therapy in WT C57BL/6 mice (**F**) or MASH C57BL/6 mice (G) bearing Hep53.4 tumors. The tumor burden was monitored by an *in vivo* imaging system. (H) Comparison of metastasis-free survival in HCC patients (non-users versus SSRI users). HR, hazard ratio. Values were compared by the log-rank test. In all panels, *p < 0.05, **p < 0.01, ***p < 0.001. Values as mean ± SD and compared by two- way ANOVA with Dunnett’s multiple comparisons (**D**), the log-rank test (**E**, **H**), and one-way ANOVA multiple comparisons with Tukey’s method (**F**, **G**).

To test whether SSRIs can increase the responsiveness of immune checkpoint blockade therapy, we engrafted mouse Hepa1-6 cells into the immune-competent C57BL/6 mice and evaluated the impact of citalopram and/or anti-PD-1 treatment. As Hepa1-6 tumor is highly immunogenic, anti-PD-1 therapy at 5 μg/g body weight were sufficient to block tumor growth. Therefore, we reduced the dose of anti-PD-1 therapy to 1 μg/g body weight and found the tumor-suppressive effect induced by anti-PD-1 therapy can be enhanced by 5 mg/kg citalopram treatment (Figure S8B). Given that the use of Hepa1-6 is not syngeneic but somehow allogeneic in C57BL/6 animals, we orthotopically injected the syngeneic Hep53.4 cells in the WT C57BL/6 mice and determined the impact of citalopram and anti-PD-1 treatment. To investigate HCC of MAFLD etiology, MASH mice were also employed for generation of the syngeneic Hep53.4 tumor model. Compared to the anti-PD-1 or citalopram treatment alone, a combined treatment robustly reduced tumor burden indicative of an additive or synergistic effect (Figure 6F) in the WT C57BL/6 mice. By contrast, MASH mice were recalcitrant to anti-PD-1 treatment but a combined treatment was still more effective than citalopram treatment alone, suggesting that citalopram also potentiates PD-1 immunotherapy in MASH ones (Figure 6G).

### SSRIs use is associated with reduced disease progression in HCC patients

Finally, we determined whether SSRIs for alleviating HCC are supported by real-world data. A total of 3061 patients with liver cancer were extracted from the Swedish Cancer Register. Among them, 695 patients had been administrated with post-diagnostic SSRIs. As shown in Figure 6H, the Kaplan-Meier survival analysis suggested that patients who utilized SSRIs exhibited a significantly improved metastasis-free survival compared to those who did not use SSRIs, with a P value of log-rank test at 0.0002. Cox regression analysis showed that SSRI use was associated with a lower risk of metastasis (HR = 0.78; 95% CI, 0.62-0.99).

## Discussion

Repurposing clinically approved drugs with known targets for disease treatment has several advantages, including but not limited to more cost-effectiveness and less side effects. Given the well-characterized link between SSRIs use and reduced cancer risk, SSRIs might be repositioned as an adjuvant therapy for cancer treatment. This attractive concept led us to explore the mechanisms of action of SSRIs in cancers.

Using HCC as a model, we reported here that SSRIs exhibit distinct anti-tumor effects in diverse mouse models of HCC. Consistently, escitalopram has been demonstrated to inhibit cell proliferation by inducing autophagy and fluoxetine induces cell apoptosis through extrinsic/intrinsic apoptotic signaling pathways in human HCC cells ^35, 36^. In a diethylnitrosamine/carbon tetrachloride (DEN/CCl_4_)-induced primary liver mouse model, sertraline and fluoxetine synergize with sorafenib to inhibit tumor progression ^37^. In zebrafish, SSRIs block β-catenin-driven liver tumorigenesis ^38^. Notably, the beneficial effects of SSRIs on tumor development have been revealed in a mouse model of chronic stress, which highlights the influences of SSRIs on the neuro-endocrine-immune system ^15^. Different from those mechanisms of action, we highlight the reprogrammed glucose metabolism of cancer cells imposed by SSRIs. This is not contradictory to previous studies as the anti-cancer mechanism of SSRI is expounded from different research aims and strategies. On the contrary, our findings further broaden the underlying cellular mechanisms of SSRIs in cancers.

Characterization of SSRIs targets is central to understanding their anti-tumor effects. Using drug repurposing approach i.e. drug-disease-gene association networking by the coordination of bioinformatics, we revealed the heretofore undescribed target of the SSRI citalopram, GLUT1. Moreover, we ruled out that SERT is not the functional target of SSRIs in HCC cells. Drugs that exert strong efficiency for any off-target in a disease can be explored further in some other diseases. Indeed, many off-targets of SSRIs have been reported, such as voltage dependent anion channel 1 (VDAC1) ^18^, G protein-coupled receptor kinase 2 (GRK2) ^39^, SMPD1 ^17^, nucleotide-binding domain, leucine-rich repeat receptor, and pyrin domain-containing protein 3 (NLRP3) ^40^, β-Amyloid ^41^, and serine hydroxymethyltransferase (SHMT) ^42^. GLUT1 is a rate-limiting transporter for glucose uptake, and its expression correlates with the glycolytic flux in cancers ^43^. GLUT1 is highly expressed in a subset of liver cancer and functionally promotes tumorigenicity ^44^. RNA-interference or pharmacological inhibition of GLUT1 inhibits the Warburg effect and reduces liver tumorigenicity ^45^. Accordingly, we found that citalopram significantly inhibits the Warburg effect of HCC cells, and this effect depends on the direct binding of GLUT1 E380, rather than citalopram-mediated decrease in the expression of glycolytic genes, including GLUT1. While we highlight the anti-HCC effects of SSRIs through targeting GLUT1, it is possible that SSRIs impact tumor progression by modulating the systemic and local microenvironments via serotonergic mechanisms. Furthermore, SSRIs may exert partial anti-tumor effects through non-tumor cell types such as hepatic stellate cells and other types of immune cells, which respond to 5-HT ^46, 47^.

In clinical settings, we reinforced SSRIs as potential therapeutic drugs for HCC treatment with three lines of evidence, 1) citalopram exhibits significant anti-tumor effects in the PDX model, especially in PDX with high glycolytic capacity; 2) citalopram potentiates the immune checkpoint blockade therapy; 3) HCC patients who used SSRIs had better metastasis-free survival. ^18^F-FDG-PET/CT has the advantage to study glucose metabolism mechanisms and SUVmax value can be used to reflect the glucose uptake of a tumor. We confirmed that HCC with high SUVmax value and GLUT1 expression responded better to the treatment of SSRIs. This suggests that SUVmax value and GLUT1 status detected by immunohistochemistry can be used as indicators for clinic use of SSRIs in HCC patients.

The restriction of acquiring the high-resolution crystal structure of SSRI bound to GLUT1 prevents further decoding the models by which SSRIs inhibit the binding of glucose to GLUT1. Several side effects have been reported with long-term use of SSRIs, such as somnolence, memory impairment, fatigue, pruritus, and headaches ^48^. While the primary mechanism of action of SSRIs is similar, each SSRI has its own unique pharmacodynamics, pharmacokinetics, and side effect profile. For drug repurposing, it will be necessary to investigate the anti-tumor effects of SSRIs at optimal doses in cancer patients without depression. This will ensure the most effective and well-tolerated treatment for individuals while also aiming to avoid or minimize any potential side effects.

In conclusion, we report the potential of repurposing SSRIs for HCC treatment using multifaceted approaches and identify a new cancer-relevant target of SSRIs. Mechanistically, we characterize that the SSRI citalopram inhibits the Warburg effect, at least to some extent, through GLUT1-dependent mechanisms. Lastly, we propose that SSRIs should be considered for clinical evaluation in the therapeutic management of a subset of HCC patients.

## Author contributions

Conceptualization: S.H.J.; Methodology: F.D., S.H.J., K.S., K.H., S.Z., L.J.J. and J.J.; Formal Analysis: F.D., S.H.J., S.Z., J.J. and L.J.J.; Resources: K.H., M.X.F., J.J., and S.H.J.; Investigation: all authors; Supervision: S.H.J.; Funding Acquisition: S.H.J., H.H.Z., J.J., X.H., Z.B. and F.D.; Writing-Original Draft: F.D., K.S., S.Z., L.J.J. and S.H.J; Writing-Review & Editing: all authors.

## Acknowledgments

The research was supported by grants from National Natural Science Foundation of China (82173153, 82001469, 8207158), Shanghai Pilot Program for Basic Research- Shanghai Jiao Tong University (21TQ1400225), Innovative research team of high-level local universities in Shanghai (SHSMU-ZDCX20210802), MAS Cancer to Jianguang Ji, “Rising Stars of Medical Talents” Youth Development Program (Youth Medical Talents-Specialist Program), Clinical Medicine Research Center Construction Project of Huadong Hospital (LCZX2202), Shanghai Outstanding Young Medical Personnel Training Program & Excellence Project of Shanghai Municipal Health Commission (20224Z0009), and Key specialized diseases construction of Huadong Hospital (ZDZB2225).

