## Supplementary file for "SSRI antidepressant citalopram reverses the Warburg effect to inhibit hepatocellular carcinoma by directly targeting GLUT1"

**Supplementary Materials**

**Supplementary Methods**

**Animal models**

In the subcutaneous xenograft model, BALB/c nude mice (male, 6-8-week-old) were kept on a 12-hr day/night cycle with free access to food and water. For the pharmacological inhibition study, 2 × 10^6^ human HCC cells (HCC-LM3 and Huh7) in 100 μl Hanks buffered saline (HBS) were injected subcutaneously in the flanks, and mice were randomly divided into indicated groups (DMSO, citalopram, escitalopram, fluoxetine, fluvoxamine, paroxetine, and sertraline) when visible tumors developed. Mice in the drug groups were given intraperitoneal injections of SSRIs (5 mg/kg body weight, Selleck, Shanghai, China) daily for indicated days, while the control group was treated with saline with 0.01% DMSO. For genetic inhibition study, 2 × 10^6^ human HCC cells (shControl, sh*SLC6A4*-#1 or sh*SLC2A1*-#1 HCC-LM3 and Huh7) in 100 μl HBS were injected subcutaneously in the lower back. When bore visible tumors, mice were given intraperitoneal injections of vehicle or citalopram (5 mg/kg body weight, Selleck, Shanghai, China) daily.

In the orthotopic xenograft model, 5 × 10^6^ Hepa1-6 cells transfected with luciferase-expressing lentiviruses were orthotopically transplanted into the liver of WT or MASH C57BL/6 mice. The day after tumor cell implantation, mice were subjected to treatment with citalopram (5 mg/kg body weight) or DMSO. Tumor burden was detected using an IVIS Spectrum system (PerkinElmer, Waltham, MA, USA). In the diet-induced MASH model, mice were fed with choline-deficient, amino acid-defined high-fat diet (45 kcal% fat) containing 0.1% methionine (CDAHFD diet, A06071309, Research Diets) for 6 weeks.

For the anti-PD-1 therapy experiment, 5 × 10^6^ Hepa1-6 or Hep53.4 cells were orthotopically transplanted into the liver of WT C57BL/6 or MASH C57BL/6 mice. Mice were randomly divided and were given intraperitoneal injections of citalopram (5 mg/kg body weight, daily), anti-PD-1 (clone RMP1-14, BioXCell, 1 μg/g or 5 μg/g body weight, every three days), combined citalopram and anti-PD-1, or DMSO control.

For generation of patient-derived xenograft models, two HCC samples were obtained from the Department of Liver Surgery, Ren Ji Hospital, School of Medicine, Shanghai Jiao Tong University. None of the patients had received radiotherapy, chemotherapy, hormone therapy, or other related anti-tumor therapies before surgery. The protocols for the collection of human HCC specimens were approved by the ethics committee of the Ren Ji Hospital, School of Medicine, Shanghai Jiao Tong University. Written informed consent was obtained before enrollment according to the guidelines of the Declaration of Helsinki. Primary HCC tissues (F0) were immediately collected from two patients after surgical resection, washed with antibiotics-containing saline to remove blood components, diced into 2-3 mm pieces, and subcutaneously implanted into the flanks of 6–8-week-old nude mice. When the xenograft tumors (F1) reached 500 mm^3^, the tumor tissues were harvested for serial transplantation to the next generation (F2, F3). In the F3 generation, PDX tumors identified as GLUT1^high^ and GLUT1^low^ were randomly allocated and subjected to intraperitoneal injections of citalopram (5 mg/kg body weight) and/or 2-DG (500 mg/kg body weight) every other day for 4 weeks. Xenograft tumor specimens were fixed with 4% paraformaldehyde, embedded in paraffin, and processed for immunohistochemical staining or were directly frozen into liquid nitrogen for further analysis.

For animal studies, our study exclusively examined male mice. It is unknown whether the findings are relevant for female mice. Tumor volume was recorded by caliper measurements and was calculated by the formula: tumor volume =1/2(length × width^2^). Length reflects the largest diameter, and width indicates the smallest diameter. At the endpoint, mice were sacrificed, the tumor was isolated and tumor weight was measured. All animals received humane care according to the criteria outlined in the “*Guide for the Care and Use of Laboratory Animals*” prepared by the National Academy of Sciences and published by the National Institutes of Health (NIH publication 86-23 revised 1985). The protocols for animal studies were approved by the ethics committee of the Ren Ji Hospital, School of Medicine, Shanghai Jiao Tong University (approval number 202201427 and RA-2021-096). All mice were randomly allocated into each experimental group. The investigator performing the experiment was always blinded to the treatment and a second blinded investigator performed the downstream data analysis.

**Cell culture**

Huh7, HCC-LM3, SNU-398, Hep3B, SMMC-7721, SNU-475, MHCC-97L, SK-Hep1, HepG2, LO2, Hepa1-6 and Hep53.4, MC38, THP-1, and HEK293T were supplemented with a culture medium as suggested by ATCC protocol mixed with 10% fetal bovine serum (FBS; Gibco, USA) and 1% (v/v) streptomycin-penicillin (Sigma-Aldrich, Shanghai, China) at 37°C in a humidified incubator containing 5% CO_2_. All cell lines mentioned above have been tested for short tandem repeat profiling and mycoplasma contamination before cell experiments.

**Cell viability, cell proliferation, and cell apoptosis assay**

For SSRI IC50 analysis, 10,000 HCC cells were seeded in triplicate in 96 well plates. After 24 h, culture media was replaced with fresh media containing increasing concentration (0, 0.1, 1, 5, 10, 50, and 100 μM) of the indicated SSRIs. DMSO was used for vehicle samples. After 48 h of incubation, cell viability was measured using Cell Counting Kit-8 (CCK-8, Dojindo Molecular Technologies, Japan) following the manufacturer’s protocol. Cell viability was indicated by measuring absorbance at 450 nm using a Power Wave XS microplate reader (BIO-TEK, USA). Data were normalized to vehicle samples, transformed, and analyzed with a nonlinear regression curve fit to generate IC50 values with GraphPad Prism 9 (Graphpad software).

For cell proliferation analysis, a total of 1,000 indicated HCC cells were seeded into 6-well plates with three replicates. After incubation with SSRIs for 10-14 days, the plates were washed with phosphate-buffered saline (PBS) twice, fixed with methanol for 10 min, and stained with 0.5% (w/v) crystal violet solution for 10 min. Colonies containing more than 30 cells were counted under the microscope.

Cell apoptosis in liver cancer cells upon SSRIs treatment was measured by the Apo-ONE® Homogeneous Caspase-3/7 Assay (Promega, Madison, WI, USA) as reported previously ^54^. As a normalization, cell number in the same plate was monitored using CellTiter-Blue (Promega, G8081). Relative Caspase-3/7 activity was calculated as the ratio of Apo-ONE/CellTiter-Blue signals.

**shRNA- and sgRNA-mediated gene silencing**

The specific shRNA oligonucleotides against *SLC6A4*/*Slc6a4* and *SLC2A* were synthesized by Genepharma (Shanghai, China). Lentivirus packaging was performed in HEK293T cells with Lipofectamine 2000 (Invitrogen, Carlsbad, CA) and virus titers were determined according to standard protocols. When grown at 70% confluence, HCC cells (Huh7, HCC-LM3, Hep53.4 and Hepa1-6) were infected with the supernatant-containing viruses in the presence of 6 μg/ml polybrene (Sigma-Aldrich, H9268, St. Louis, MO). Cells expressing shRNAs were selected with puromycin (2 μg/ml, Gibco, A1113802). For CRISPR/Cas9-mediated gene knockout, Huh7 and HCC-LM3 cells were first transfected with lentiCas9-Blast vector to express Cas9. To construct the SERT^−/−^ HCC cell lines, a lentiviral delivery system using guide RNA (sgRNA) targeting *SLC6A4* was employed. Lentivirus was generated in HEK293T by transfecting lentiGuide-Puro with packaging plasmids pVSVg and psPAX2. The transfection was performed with Fugene6 (E2693, Promega) in OptiMEM. Two days later, the supernatants were filtered through a 0.45 μm filter, and Cas9-expressing cells were infected with lentivirus in the presence of 6 μg/ml polybrene. Individual colonies were seeded into 96-well plates, grown to confluence and then subjected to knockout efficiency analysis. Sequences for shRNA and sgRNA were available at **Table S8**.

**Quantitative real-time PCR (qRT-PCR)**

RNA was extracted from liver cancer cells or xenograft tissues using the RNAiso Plus reagent (Takara, Japan) and subjected to reverse transcription using the PrimeScript RT-PCR kit (Takara, Japan). Then, qRT-PCR was performed using the 7500 Real-time PCR system (Applied Biosystems, USA). The housekeeping gene *GAPDH* was used to normalize gene expression. Primer sequences used in this study were listed in **Table S9**.

**RNA sequencing analysis**

The HCC-LM3 cells were treated with citalopram or fluvoxamine (5 μM) for 24 h and then underwent RNA sequencing analysis. Total RNA was extracted using the Trizol method, and the quality of both the RNA and cDNA library was assessed before performing paired indexing sequencing on the Illumina NovaSeq 6000. The sequencing process was controlled by data collection software provided by Illumina, and the resulting data was analyzed in real-time. The raw data is available for access from the Sequence Read Archive (SRA) repository under the accession numbers PRJNA1084911.

**Immunoblotting**

Whole cellular lysates were extracted from liver cancer cells using RIPA lysis buffer (P0013B, Beyotime, Shanghai, China) mixed with protease and phosphatase inhibitor cocktails (ab201119, Abcam, Shanghai, China). The protein content was determined using a BCA Protein Assay Kit (Pierce Biotechnology, USA). Proteins were subjected to sodium dodecyl sulfate-polyacrylamide gel electrophoresis (SDS-PAGE) and then transferred to polyvinylidene difluoride (PVDF, Millipore) membranes. After blocking with 5% (m/v) skim milk for 1 h at room temperature, the membranes were incubated with indicated antibodies at 4 °C overnight. The next day, the membranes were washed with PBS three times and incubated with HRP-conjugated secondary antibodies for 45 min, followed by an ECL chemiluminescence assay (SB-WB012, Share-bio, Shanghai, China) with a Bio-Spectrum Gel Imaging System (Bio-Rad). The following antibodies are used in this study: SERT (1:500, ProteinTech, 19559-1-AP), GLUT1 (1:2,000, Cell Signaling Technology, #73015), and β-actin (1:2,000, Abcam, ab8226). The information for secondary antibodies is as follows: goat anti-rabbit IgG (H+L) cross-adsorbed secondary antibody (1:5,000, Thermo Fisher Scientific, G-21234) and goat anti-mouse IgG (H+L) cross-adsorbed secondary antibody (1:5,000, Thermo Fisher Scientific, G-21040).

**Histology, immunohistochemistry, and immunofluorescence analysis**

H&E and Sirius red staining was routinely performed to characterize hepatic steatosis, ballooning degeneration, and fibrosis. Paraffin-embedded sections of mouse and human HCC tissues were subjected to immunohistochemical analysis of SERT, Ki67, cleaved caspase 3, and GLUT1 expression. A tissue microarray containing 202 HCC cases, as reportedly previously ^55^, was used to determine GLUT1. Immunohistochemical analysis was performed as reported previously ^54^. The primary antibodies used in this study were shown as follows: SERT (1:200, ProteinTech, 19559-1-AP), Ki67 (1:400, Cell Signaling Technology, #9449), cleaved caspase 3 (1:400, Cell Signaling Technology, #9661), and GLUT1 (1:500, Cell Signaling Technology, #73015). Scoring of GLUT1 was conducted based on the percentage of positive staining cells or staining intensity.

**Target prediction**

The SuperPred database (https://prediction.charite.de/index.php) was used for target prediction of SSRIs. Briefly, each SSRI was searched by its name via PubChem and then subjected for calculation. The known strong binders and additionally predicted targets were downloaded. Finally, the predicted targets of all SSRIs were merged by Venn analysis (http://bioinformatics.psb.ugent.be/webtools/Venn/).

**Global inverse gene-expression profiling analysis**

This approach is to screen for signatures associated with target genes that are "inverse" to the drug-induced gene expression profile. Total RNA from HCC-LM3 cells upon citalopram or fluvoxamine was extracted with TRIzol reagent (TaKaRa, Japan) followed by purification using an RNA 6000 Nano LabChip kit (Agilent Technologies, USA). Total RNA was dissolved in RNase-free water and the RNA quality was determined by a Bioanalyzer 2100 (NanoDrop, Agilent Technologies, USA). RNA of cell samples was sequenced by BGI NGS platforms (Shenzhen, China). The DEseq2 package was used to identify DEGs (citalopram vs DMSO and fluvoxamine vs DMSO). Co-upregulated and co-downregulated genes were used for the generation of an SSRI-induced gene signature. The RNAseq data of LIHC samples in the TCGA cohort (n = 371, https://portal.gdc.cancer.gov) were subjected for GSEA with SSRI-induced gene signature. Grouping was made based on the median gene expression value of predicted targets.

**Drug affinity responsive target stability (DARTS) assay**

HCC-LM3 and HEK293T cells were lysed with NP-40 lysis buffer (Thermo Fisher Scientific, 89842Y). After centrifugation at 12,000 rpm for 10 min, the supernatant was collected, and protein concentration was quantified by a BCA Protein Assay Kit (Pierce Biotechnology, USA). Samples in equivalent quality were treated with indicated SSRIs and vehicle at 37°C for 4 h. Then, pronase (10 μg/mL, Roche, 10165921001) or distilled water was added and allowed to incubate at room temperature for indicated time points (10 and 30 min). The reaction was stopped by adding protease inhibitors and the products were harvested and analyzed with Western blotting.

***In Silico* docking study**

To predict the binding mode of citalopram/GLUT1, molecular docking was performed using the AutoDock 4.2.6 software package ^56^. The receptor structure for GLUT1 (PDB id: 4PYP) ^32^ was obtained from the RCSB protein data bank (https://www.rcsb.org/). The absolute structure of citalopram (CID: 2771), fluoxetine (CID: 3386), fluvoxamine (CID: 5324346), paroxetine (CID: 43815), and sertraline (CID: 68617) were obtained from the PubChem database (https://pubchem.ncbi.nlm.nih.gov/). For each docking scenario, molecular docking was performed using a standard protocol ^57^ except that the number of genetic algorithm runs was set to 500 to extensively sample the binding modes. The binding energy and cluster information were obtained directly from the AutoDock output.

**Glucose, lactate, and ATP levels**

As reported previously ^54^, the Amplex Red Glucose/Glucose Oxidase Assay Kit (Thermo Fisher Scientific, A22189, USA) and the Lactate Assay Kit (BioVision, K607-100, USA) were used to determine glucose and lactate levels in the cell culture supernatants ^54^. The enhanced ATP assay Kit (Beyotime Biotechnology, S0027, Shanghai, China) was used to detect ATP generation in liver cancer cells according to the manufacturer’s instructions. Total protein level was used to normalize glucose uptake, lactate production, and ATP generation. All the above experiments were run in triplicate and repeated at least two times.

**Extracellular acidification rate (ECAR)**

ECAR in HCC cells was analyzed with the Seahorse Bioscience XF96 Extracellular Flux Analyzer (Seahorse Bioscience, USA) with the Seahorse XF96 Cell Glycolysis Stress Test Kit (Seahorse Bioscience, USA) according to the manufacturer’s instructions. Seahorse XFe96 FluxPak column was hydrated with distilled water at 37°C (CO_2_-free) overnight and equilibrated in Seahorse XF Calibrant solution for 1 h on the day of analysis. To test ECAR, cells were incubated with an unbuffered medium followed by sequential injection of 10 mM glucose, 1 μM oligomycin, and 10 mM 2-DG. ECAR data were collected using Wave Controller 2.6 (Agilent Technologies) and normalized to total protein content and reported as mpH/min. Each datum was determined in triplicate.

**Biochemical analysis**

The activity of serum alanine aminotransferase (ALT) and aspartate aminotransferase (AST) was used to reflect liver function. The ALT and AST activity in serum were measured using the commercial detection kits (MU30059 and MU30060, Bioswamp Life Science Lab, Wuhan, China), following the manufacturer’s guidelines.

**Supplementary Figures**

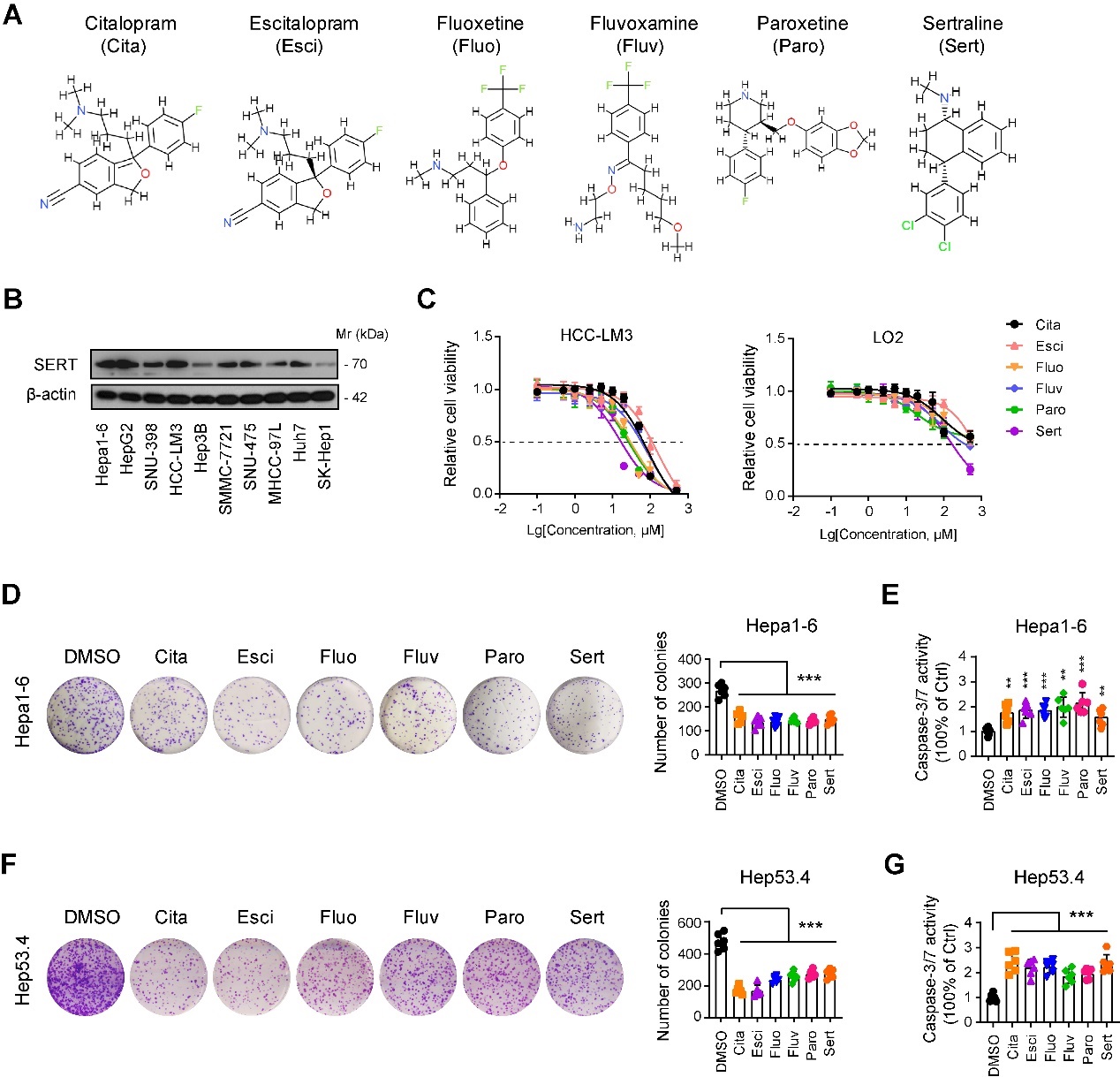

**Figure S1. The *in vitro* anti-tumor effects of SSRIs in liver cancer**. (**A**) The chemical structure for SSRIs. Data were obtained from Super-PRED (https://prediction.charite.de/). (**B**) Western blotting showed SERT protein expression in liver cancer cell lines. (**C**) IC_50_ analysis of SSRIs in HCC-LM3 and LO2 cells. (**D**, **F**) Plate colony formation assay showed the effect of SSRIs on the long-term cell proliferation of Hepa1-6 and Hep53.4 cells (n = 6 per group). (**E**, **G**) Caspase-3/7 activity in Hepa1-6 and Hep53.4 cells upon treatment with 5 μM SSRIs for 48 h (n = 6 per group). In panel **D**-**G**, **p < 0.01, ***p < 0.001. Values as mean ± SD and compared by one-way ANOVA multiple comparisons with Tukey’s method. Data are representative of three independent experiments (**C**-**G**).

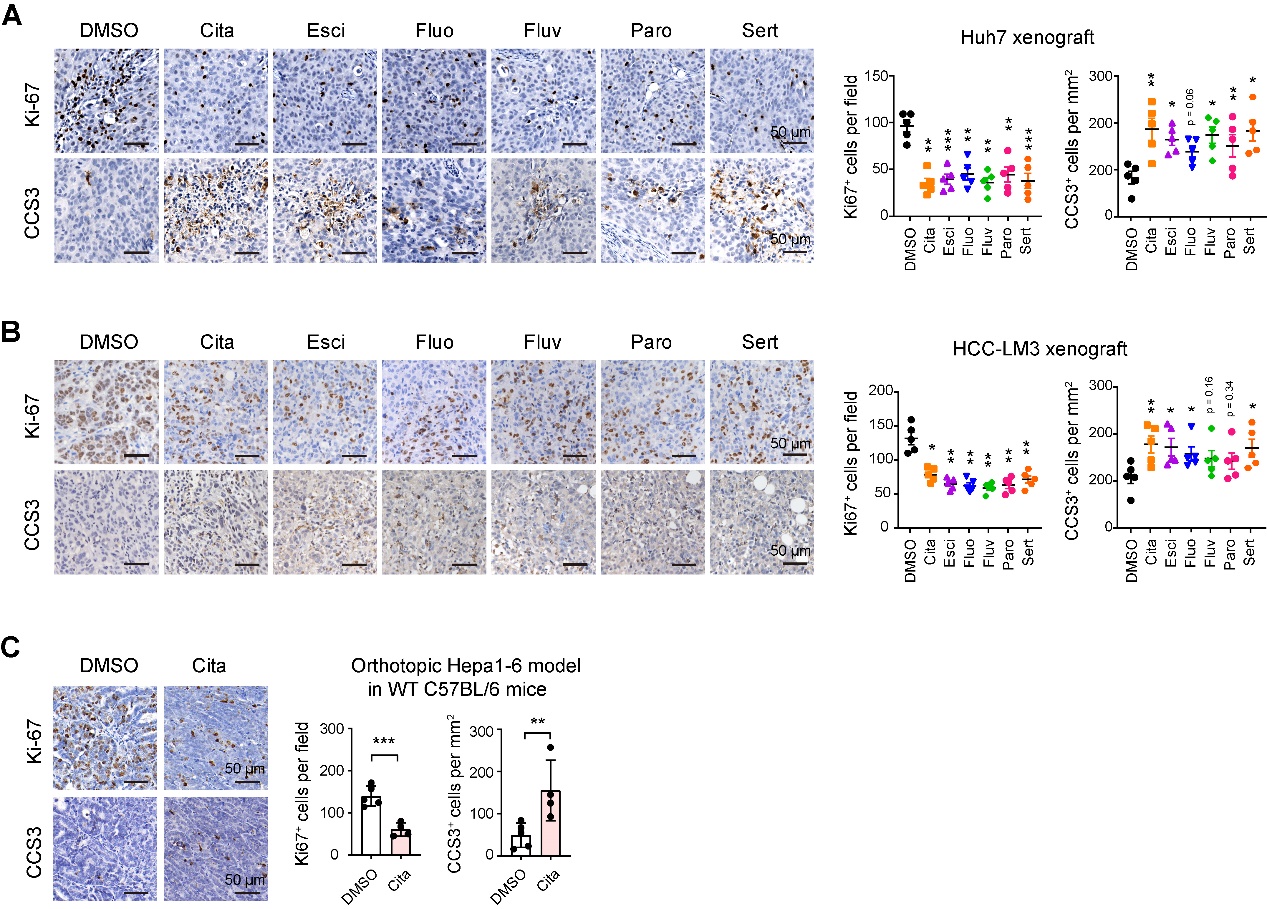

**Figure S2. SSRIs inhibit HCC cell proliferation and promote cell apoptosis in different preclinical models.** (**A** and **B**) Immunohistochemical analysis of cleaved caspase-3 (CCS3) and the proliferation index Ki67 in Huh7- or HCC-LM3-bearing subcutaneous xenograft tumors, treated with six commonly prescribed SSRIs. (**C**) Immunohistochemical analysis of CCS3 and Ki67 in Hepa1-6-bearing orthotopic xenograft tumors from WT C57BL/6 mice, treated with DMSO or 5 mg/kg citalopram. In all panels, *p < 0.05, **p < 0.01, ***p < 0.001. Scale bar, 50 μm. Values as mean ± SD and compared by one-way ANOVA multiple comparisons with Tukey’s method among groups (**A**, **B**), and the Student’s t test (**C**).

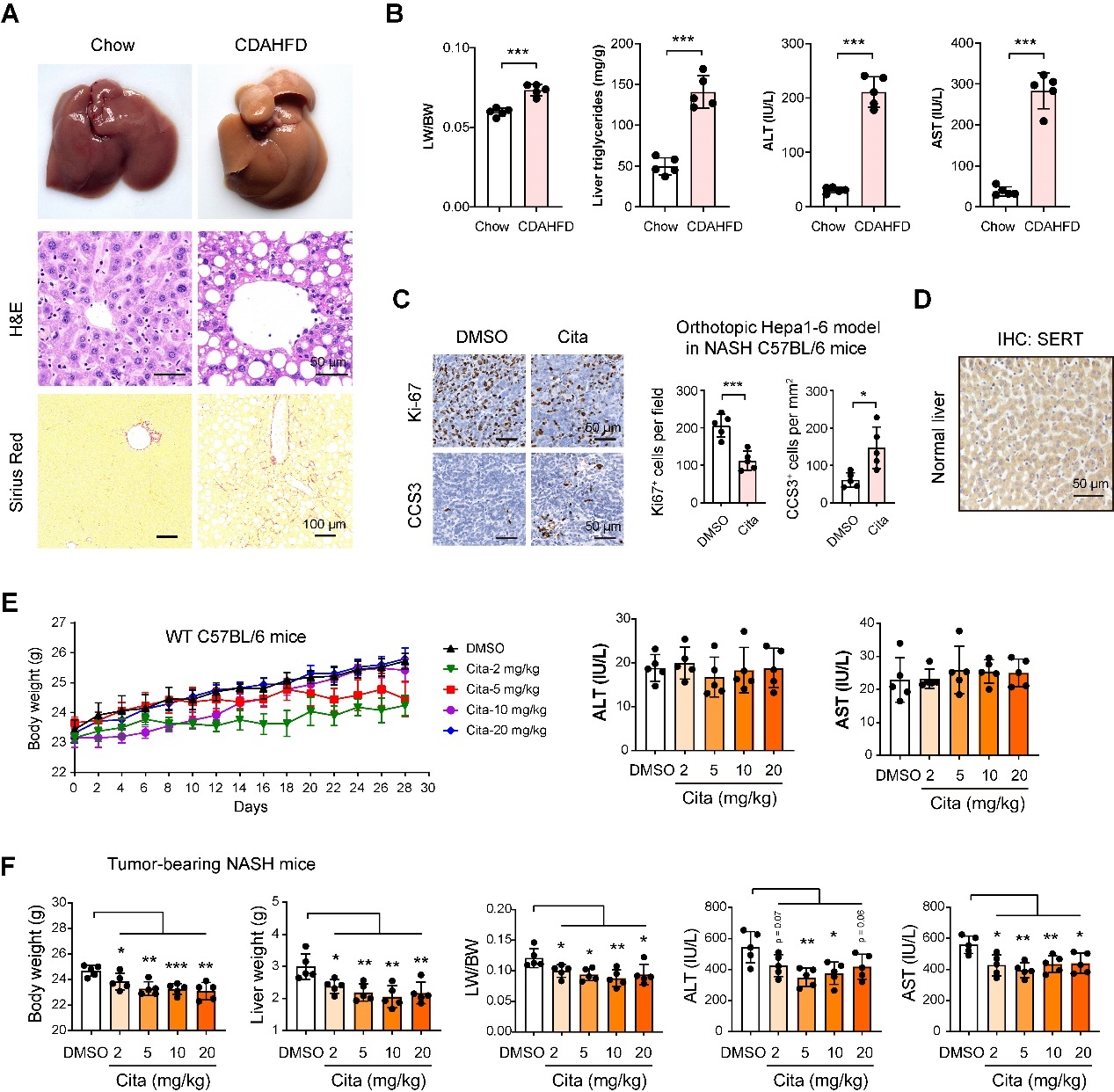

**Figure S3. Measurement of potential side effects of citalopram.** (**A**) H&E and Sirius red staining showed the liver changes of mice fed with chow diet or CDAHFD. (**B**) Comparison of liver weight, body weight, and liver-to-body weight ratio in mice fed with chow diet or CDAHFD (n = 5 per group). (**C**) Immunohistochemical analysis of CCS3 and Ki67 in Hepa1-6-bearing orthotopic xenograft tumors from NASH C57BL/6 mice (CDAHFD model), treated with DMSO or 5 mg/kg citalopram. (**D**) Immunohistochemical images showed SERT protein expression in normal liver tissues. Scale bar, 50 μm. (**E**) Body-weight curves, and serum ALT and AST levels in WT C57BL/6 mice treated with different doses of citalopram (n = 5 per group). (**F**) Body weight, liver weight, liver-to-body weight ratio, and serum ALT and AST levels in NASH C57BL/6 mice treated with different doses of citalopram (n = 5 per group). In all panels, *p < 0.05, **p < 0.01, ***p < 0.001. Values as mean ± SD and compared by the Student’s t test (**B**, **C**) and one-way ANOVA multiple comparisons with Tukey’s method among groups (**F**).

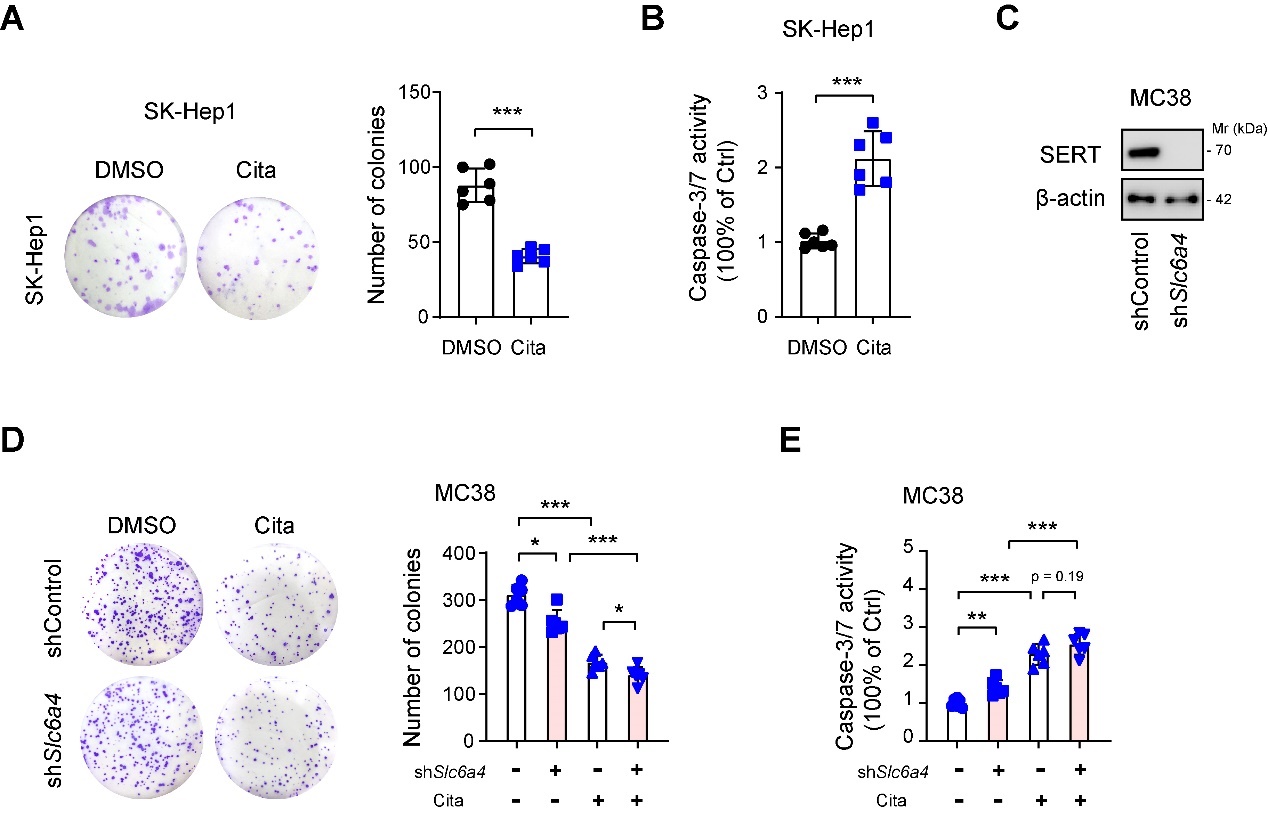

**Figure S4. The SERT-independent anti-tumor effects of citalopram.** (**A**) Plate colony formation assay revealed the effect of citalopram treatment on the long-term cell proliferation of SK-Hep1 cells (n = 6 per group). (**B**) Caspase-3/7 activity in SK-Hep1 cells upon treatment with 5 μM citalopram (n = 6 per group). (**C**) A specific shRNAs were used for the genetic silencing of SERT in MC38 cells and the knockdown efficiency was verified by Western blotting analysis. (**D**) Plate colony formation assay revealed the effect of SERT knockdown alone or combined with citalopram treatment on the long-term cell proliferation of MC38 cells (n = 6 per group). (**E**) Caspase-3/7 activity in MC38 cells upon SERT knockdown or combined treatment with 5 μM citalopram (n = 6 per group). In all panels, *p < 0.05, **p < 0.01, ***p < 0.001. Values as mean ± SD and compared by the Student’s t test (**A**, **B**) and one-way ANOVA multiple comparisons with Tukey’s method among groups (**D**, **E**).

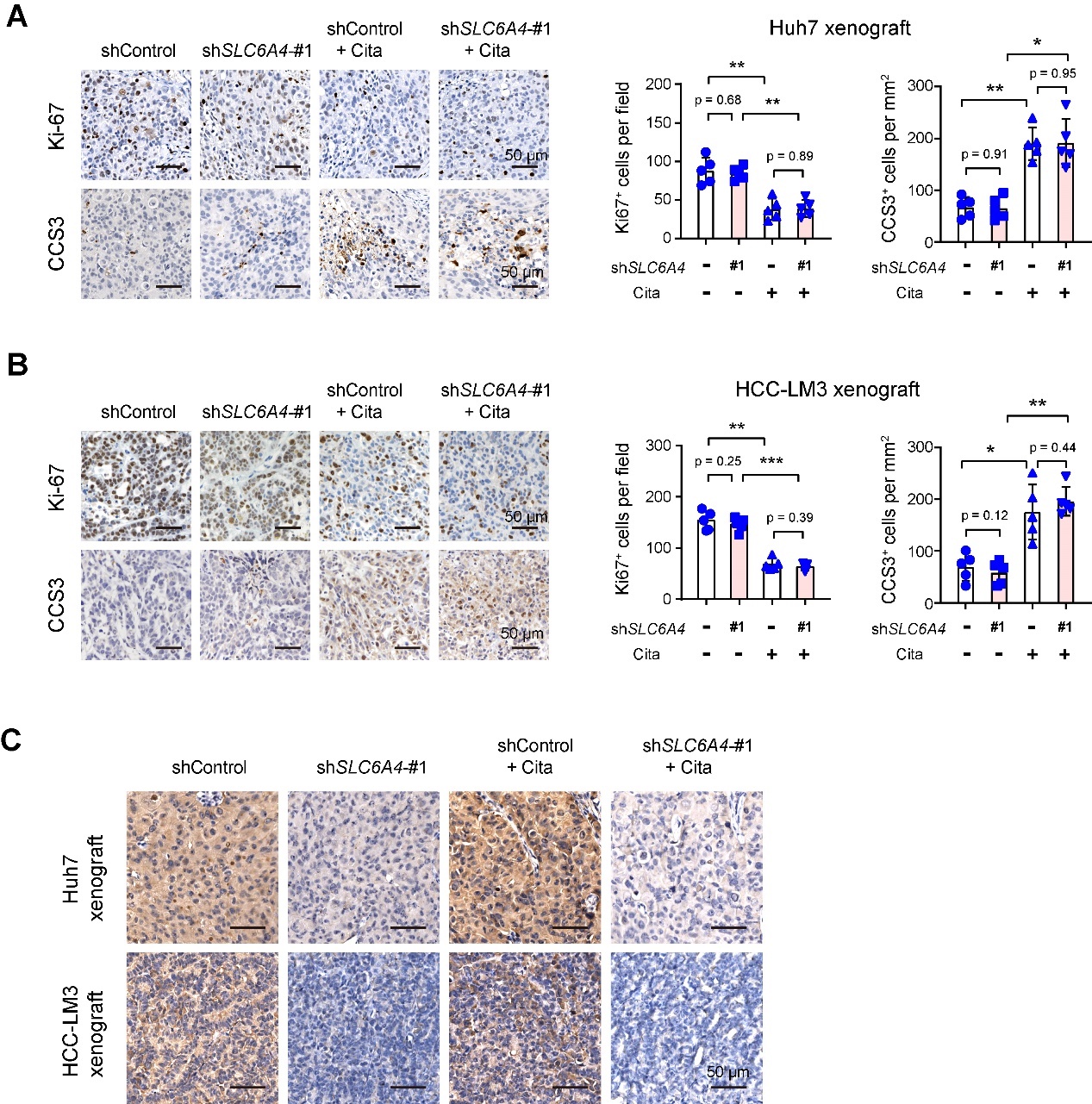

**Figure S5. Citalopram-mediated anti-tumor effects are not dependent on SERT expression in liver cancer**. (**A**, **B**) Immunohistochemical analysis of Ki-67 and CCS3 in Huh7- or HCC-LM3-derived xenograft tissues from the shControl, shControl + citalopram, sh*SLC6A4*-#1, and sh*SLC6A4*-#1 + citalopram groups. (**C**) Immunohistochemical analysis showed SERT protein expression in shControl and sh*SLC6A4*-#1 tumors, treated with DMSO or citalopram. Scale bar, 50 μm. *p < 0.05, **p < 0.01, ***p < 0.001. Values as mean ± SD and compared by one-way ANOVA multiple comparisons with Tukey’s method among groups (**A**, **B**).

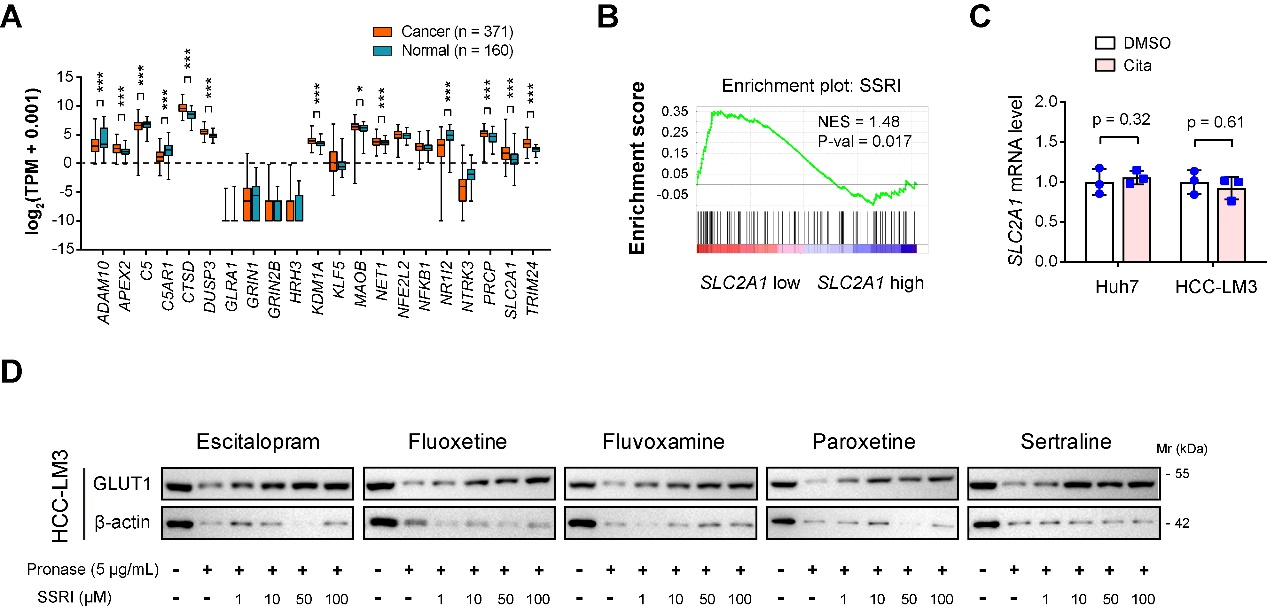

**Figure S6. Identification of potential molecular targets of SSRIs in HCC**. (**A**) The expression levels of 21 predicted targets of SSRIs in normal liver (n = 160) and HCC tissues (n = 371). Data were acquired from the GTEx and TCGA databases. (**B**) Gene Set Enrichment Analysis (GSEA) of HCC RNAseq data (TCGA cohort) with the SSRI-related gene signature. HCC samples were divided into high and low groups based on the median expression value of GLUT1. (**C**) Real-time qPCR analysis of *SLC2A1* mRNA levels upon citalopram treatment (n = 3 per group). (**D**) The DARTS assay and immunoblot analysis showed GLUT1 protein stability against 5 μg/mL pronase in the presence of different concentrations of SSRIs treatment (0, 1, 10, 50, and 100 μM). Values as mean ± SD and compared by the Student’s t test.

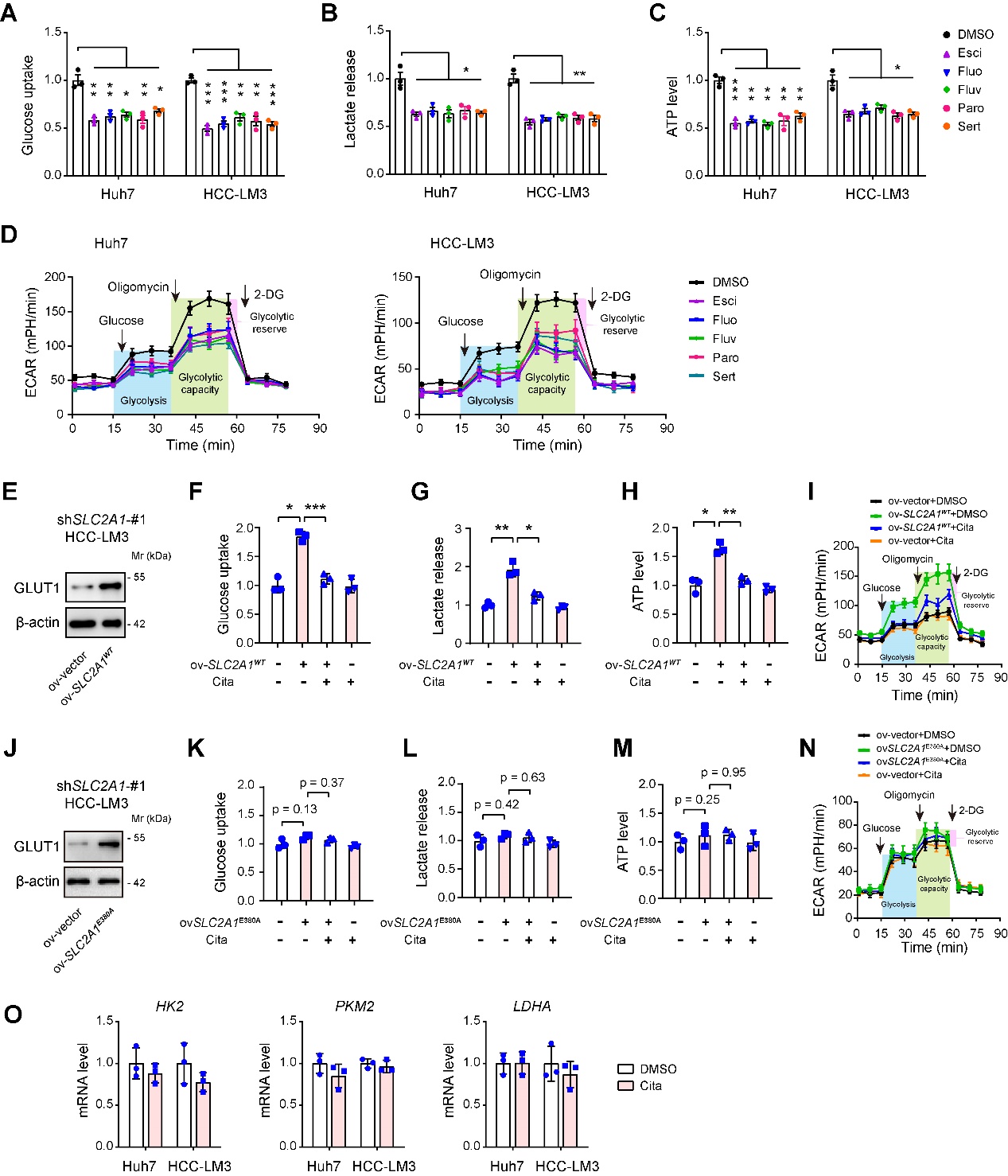

**Figure S7. Citalopram blocks GLUT1-mediated glycolytic flux.** (**A-D**) The effects of different SSRIs on the glycolytic ability of Huh7 and HCC-LM3 cells were measured, as indicated by glucose uptake (**A**), lactate production (**B**), ATP generation (**C**), and extracellular acidification rate (ECAR) (**D**, n = 3 per group). (**E**) Reconstituted expression of WT GLUT1 in GLUT knockdown (GLUT1^KD^) HCC-LM3 cells. (**F-I**) The effects of citalopram and GLUT1 overexpression on the glycolytic ability of GLUT1^KD^ HCC-LM3 cells were measured, as indicated by glucose uptake (**F**), lactate production (**G**), ATP generation (**H**), and extracellular acidification rate (ECAR) (**I**, n = 3 per group). (**J**) Reconstituted expression of mutant GLUT1 E380A in GLUT knockdown (GLUT1^KD^) HCC-LM3 cells. (**K-N**) The effects of citalopram and GLUT1 (E380A) overexpression on the glycolytic ability of GLUT1^KD^ HCC-LM3 cells were measured, as indicated by glucose uptake (**K**), lactate production (**L**), ATP generation (**M**), and extracellular acidification rate (ECAR) (**N**, n = 3 per group). (**O**) Real-time qPCR analysis of *HK2*, *PKM2*, and *LDHA* mRNA levels in Huh7 and HCC-LM3 cells upon citalopram treatment.

In all panels, *p < 0.05, **p < 0.01, ***p < 0.001. Values as mean ± SD and compared by one-way ANOVA multiple comparisons with Tukey’s method among groups (**A**-**C**, **F**-**H**, **K**-**M**) and the Student’s t test (**O**).

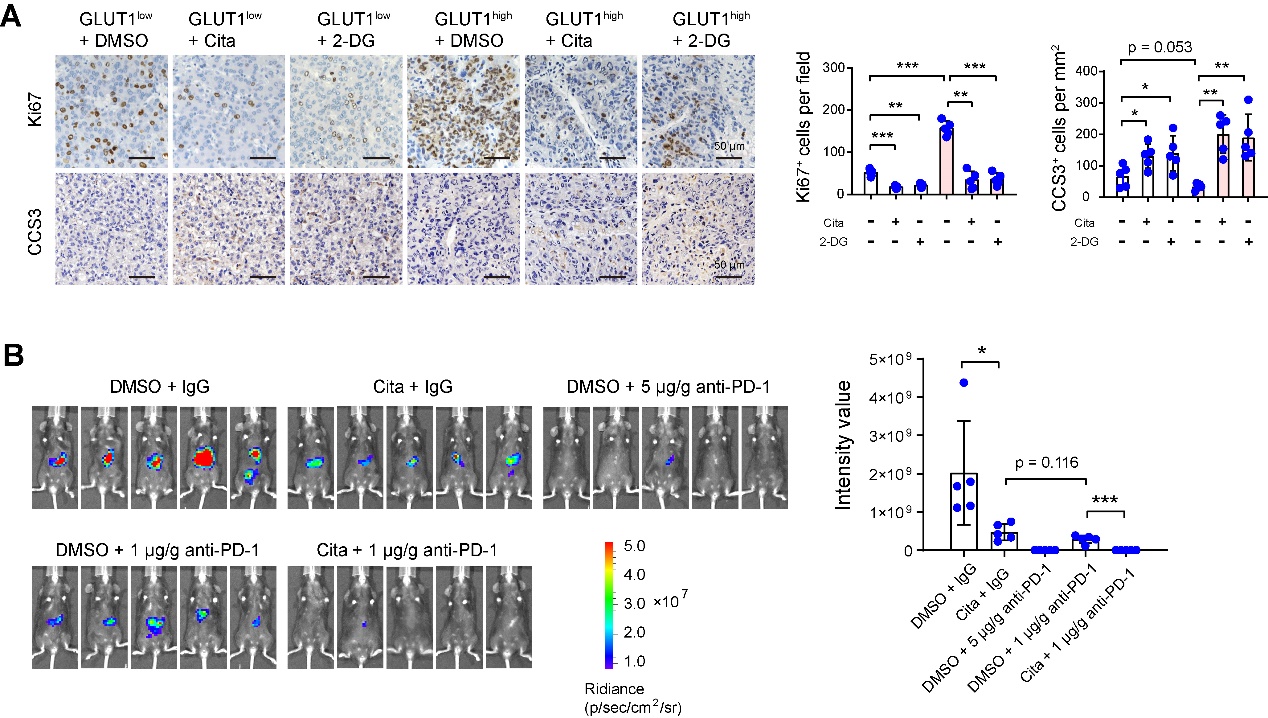

**Figure S8. The anti-tumor effects of SSRIs in the PDX model and orthotopic tumor model.** (**A**) Immunohistochemical analysis of Ki67 and CCS3 in GLUT1-low and GLUT1-high xenograft tumors from nude mice, treated with 5 mg/kg citalopram or 50 mg/kg 2-DG. Scale bar, 50 μm. (**B**) Anti-PD-1 therapy and citalopram treatment in C57BL/6 mice bearing Hepa1-6 tumors. The tumor burden of Hepa1-6 xenograft tumors in the presence of anti-PD-1, citalopram, and combined treatment was monitored by an *in vivo* imaging system (n = 5 per group).*p < 0.05, **p < 0.01, ***p < 0.001. Values as mean ± SD and compared by one-way ANOVA multiple comparisons with Tukey’s method among groups.

**Table S1. Predicted targets of Citalopram and Escitalopram**

| **Target Name** | **ChEMBL-ID** | **UniProt ID** | **PDB Visualization** | **TTD ID** | **Probability** | **Accuracy** |
| --- | --- | --- | --- | --- | --- | --- |
| Nuclear factor NF-kappa-B p105 subunit | CHEMBL3251 | P19838 | 1SVC | Not Available | 98% | 96% |
| DNA-(apurinic or apyrimidinic site) lyase | CHEMBL5619 | P27695 | 6BOW | T13348 | 96.8% | 91% |
| G-protein coupled receptor 55 | CHEMBL1075322 | Q9Y2T6 | Not Available | T87670 | 95% | 78% |
| Kinesin-like protein 1 | CHEMBL4581 | P52732 | 6TIW | T28484 | 95% | 93% |
| Androgen Receptor | CHEMBL1871 | P10275 | 3V49 | T11211 | 94% | 96% |
| Cathepsin D | CHEMBL2581 | P07339 | 4OD9 | T67102 | 93% | 99% |
| Isoprenylcysteine carboxyl methyltransferase | CHEMBL4699 | O60725 | Not Available | Not Available | 91% | 100% |
| Melanin-concentrating hormone receptor 1 | CHEMBL344 | Q99705 | Not Available | T09572 | 91% | 92.5% |
| Beta amyloid A4 protein | CHEMBL2487 | P05067 | 5BUO | T87024 | 89% | 97% |
| Kruppel-like factor 5 | CHEMBL1293249 | Q13887 | Not Available | Not Available | 88% | 86% |
| Glucose transporter | CHEMBL2535 | P11166 | 6THA | Not Available | 87.4% | 99% |
| Serine/threonine-protein kinase MST2 | CHEMBL4708 | Q13188 | 6AO5 | Not Available | 87% | 72% |
| NT-3 growth factor receptor | CHEMBL5608 | Q16288 | 6KZD | Not Available | 86% | 96% |
| Ribosomal protein S6 kinase alpha 1 | CHEMBL2553 | Q15418 | 2Z7Q | Not Available | 86% | 85% |
| Dual specificity phosphatase Cdc25C | CHEMBL2378 | P30307 | 3OP3 | Not Available | 86% | 97% |
| Nitric oxide synthase, inducible | CHEMBL4481 | P35228 | 3E7G | T02703 | 86% | 94.8% |
| Indoleamine 2,3-dioxygenase | CHEMBL4685 | P14902 | 6E43 | T89697 | 85% | 96% |
| Histone deacetylase 8 | CHEMBL3192 | Q9BY41 | 5VI6 | T28887 | 85% | 94% |
| Metabotropic glutamate receptor 5 | CHEMBL3227 | P41594 | 6N4X | Not Available | 85% | 96% |
| Nuclear factor erythroid 2-related factor 2 | CHEMBL1075094 | Q16236 | 2FLU | Not Available | 84% | 96% |
| Mineralocorticoid receptor | CHEMBL1994 | P08235 | 4PF3 | Not Available | 84% | 100% |
| Histone deacetylase 9 | CHEMBL4145 | Q9UKV0 | Not Available | Not Available | 83% | 85% |
| Lysosomal Pro-X carboxypeptidase | CHEMBL2335 | P42785 | 3N2Z | Not Available | 83% | 100% |
| ALK tyrosine kinase receptor | CHEMBL4247 | Q9UM73 | 4Z55 | T56418 | 83% | 97% |
| Glutamate carboxypeptidase II | CHEMBL1892 | Q04609 | 5O5T | Not Available | 82% | 97.5% |
| Tyrosine-protein kinase FER | CHEMBL3982 | P16591 | 6KC4 | Not Available | 82% | 78% |
| Dual specificty protein kinase CLK1 | CHEMBL4224 | P49759 | 6KHD | Not Available | 81% | 85.3% |
| DNA topoisomerase II alpha | CHEMBL1806 | P11388 | 6ZY5 | T17048 | 81% | 89% |
| TRAF2- and NCK-interacting kinase | CHEMBL4527 | Q9UKE5 | 2X7F | Not Available | 81% | 70% |
| Inhibitor of nuclear factor kappa B kinase alpha subunit | CHEMBL3476 | O15111 | 5EBZ | Not Available | 80% | 96% |
| Glycine receptor subunit alpha-1 | CHEMBL5845 | P23415 | 4X5T | T50269 | 80% | 91% |
| Beta-glucocerebrosidase | CHEMBL2179 | P04062 | 6TN1 | T84173 | 78.5% | 85% |
| Histone deacetylase 4 | CHEMBL3524 | P56524 | 2VQM | T63816 | 78% | 93% |
| ADAM10 | CHEMBL5028 | O14672 | 6BE6 | T31902 | 78% | 97.5% |
| Protein tyrosine kinase 2 beta | CHEMBL5469 | Q14289 | 4EKU | T07087 | 78% | 91% |
| LSD1/CoREST complex | CHEMBL3137262 | O60341 | 5L3D | Not Available | 77% | 97% |
| Proteasome Macropain subunit | CHEMBL3492 | P49721 | 5LE5 | Not Available | 77% | 90% |
| Signal transducer and activator of transcription 3 | CHEMBL4026 | P40763 | 6QHD | T29130 | 77% | 83% |
| G-protein coupled bile acid receptor 1 | CHEMBL5409 | Q8TDU6 | 7CFM | T86273 | 77% | 94% |
| C5a anaphylatoxin chemotactic receptor | CHEMBL2373 | P21730 | 6C1R | T15439 | 75.7% | 93% |
| Neurokinin 1 receptor | CHEMBL249 | P25103 | 6HLP | T47094 | 76% | 99% |
| Serotonin 2a (5-HT2a) receptor | CHEMBL224 | P28223 | 6WHA | T32060 | 76% | 91% |
| Beta-glucuronidase | CHEMBL2728 | P08236 | 3HN3 | T96413 | 76% | 78% |
| Serotonin 2c (5-HT2c) receptor | CHEMBL225 | P28335 | 6BQH | T83813 | 76% | 90% |
| Glutathione S-transferase Pi | CHEMBL3902 | P09211 | 5J41 | T21669 | 75% | 94% |
| Adenosine A1 receptor | CHEMBL226 | P30542 | 5N2S | T92072 | 75% | 96% |
| Dual specificity mitogen-activated protein kinase kinase 2 | CHEMBL2964 | P36507 | 1S9I | Not Available | 74.6% | 80% |
| Glutamate NMDA receptor; GRIN1/GRIN2B | CHEMBL1907603 | Q05586 | 5EWM | Not Available | 75% | 96% |
| Kelch-like ECH-associated protein 1 | CHEMBL2069156 | Q14145 | 6WCQ | Not Available | 74% | 82% |
| Acetyl-CoA carboxylase 1 | CHEMBL3351 | Q13085 | 6G2H | Not Available | 74% | 93% |
| Nicotinamide phosphoribosyltransferase | CHEMBL1744525 | P43490 | 4LVF | T54582 | 74% | 96% |
| Mitogen-activated protein kinase kinase kinase 14 | CHEMBL5888 | Q99558 | 6Z1T | Not Available | 74% | 100% |
| 5'-nucleotidase | CHEMBL5957 | P21589 | 6TVE | Not Available | 73% | 98% |
| Pregnane X receptor | CHEMBL3401 | O75469 | 6TFI | T82702 | 73.1% | 95% |
| Peroxisome proliferator-activated receptor alpha | CHEMBL239 | Q07869 | 6KAX | T86591 | 72% | 91% |
| Tyrosine-protein kinase receptor RET | CHEMBL2041 | P07949 | 6Q2O | T60631 | 71% | 92% |
| Glycine transporter 2 | CHEMBL3060 | Q9Y345 | Not Available | Not Available | 71% | 99% |
| Cytochrome P450 2A6 | CHEMBL5282 | P11509 | 2FDV | T06455 | 71% | 72% |
| Choline kinase alpha | CHEMBL3117 | P35790 | 4DA5 | T55709 | 71% | 78% |
| Geranylgeranyl transferase type I | CHEMBL2095164 | P49354 | 2H6F | Not Available | 71% | 92.8% |
| Methionine aminopeptidase 2 | CHEMBL3922 | P50579 | 1B6A | T75596 | 70% | 97% |
| Dual specificity protein kinase CLK2 | CHEMBL4225 | P49760 | 6KHE | Not Available | 70% | 81% |
| Serotonin 7 (5-HT7) receptor | CHEMBL3155 | P34969 | Not Available | T79062 | 70% | 91% |
| Dual specificity protein phosphatase 3 | CHEMBL2635 | P51452 | 3F81 | Not Available | 70% | 94% |
| Protein-arginine N-methyltransferase 1 | CHEMBL5524 | Q99873 | 6NT2 | Not Available | 70% | 97% |
| SUMO-activating enzyme | CHEMBL2095174 | Q9UBE0 | 6XOG | Not Available | 69% | 79% |
| Dual specificity phosphatase Cdc25B | CHEMBL4804 | P30305 | 1QB0 | Not Available | 69% | 79.5% |
| 11-beta-hydroxysteroid dehydrogenase 1 | CHEMBL4235 | P28845 | 4BB5 | T65200 | 69% | 98% |
| AMP-activated protein kinase, alpha-1 subunit | CHEMBL4045 | Q13131 | 6C9H | Not Available | 68% | 73.5% |
| Nitric-oxide synthase, brain | CHEMBL3568 | P29475 | 6CIC | T16117 | 68% | 95% |
| C-X-C chemokine receptor type 4 | CHEMBL2107 | P61073 | 3ODU | T96079 | 68% | 93.1% |
| Lysine-specific demethylase 4A | CHEMBL5896 | O75164 | 6G5X | Not Available | 67% | 99% |
| BMP-2-inducible protein kinase | CHEMBL4522 | Q9NSY1 | 4W9W | Not Available | 67.2% | 78% |
| Serine/threonine-protein kinase GAK | CHEMBL4355 | O14976 | 4Y8D | T00043 | 67% | 89% |
| Sodium channel protein type IV alpha subunit | CHEMBL2072 | P35499 | 6AGF | T02546 | 67% | 92% |
| Dihydrofolate reductase | CHEMBL202 | P00374 | 1KMV | T17345 | 67% | 90% |
| Cathepsin S | CHEMBL2954 | P25774 | 2C0Y | T68290 | 66.3% | 95.6% |
| Proteasome subunit beta type-9 | CHEMBL1944495 | P28065 | 6E5B | Not Available | 66% | 97.5% |
| NADPH oxidase 1 | CHEMBL1287628 | Q9Y5S8 | Not Available | Not Available | 65.9% | 95% |
| Histone deacetylase 7 | CHEMBL2716 | Q8WUI4 | 3C10 | Not Available | 65% | 89% |
| Bile acid receptor FXR | CHEMBL2047 | Q96RI1 | 6HL1 | Not Available | 65% | 96.1% |
| Fatty acid synthase | CHEMBL4158 | P49327 | 3HHD | T16514 | 65% | 82.5% |
| C-C chemokine receptor type 1 | CHEMBL2413 | P32246 | Not Available | T16016 | 65% | 89.5% |
| Aurora kinase B/Inner centromere protein | CHEMBL3430907 | Q96GD4 | 6YIH | T46781 | 65% | 97.5% |
| Integrin alpha-V/beta-3 | CHEMBL1907598 | P05106 | 6UJA | T67103 | 65% | 96% |
| Dipeptidyl peptidase IX | CHEMBL4793 | Q86TI2 | 6EOR | Not Available | 64% | 97% |
| DCN1-like protein 1 | CHEMBL4105838 | Q96GG9 | 6BG3 | Not Available | 64% | 95% |
| Integrin alpha-4/beta-1 | CHEMBL1907599 | P05556 | 3V4V | T97587 | 64% | 93% |
| Galectin-3 | CHEMBL4531 | P17931 | 6FOF | T72038 | 64% | 96.9% |
| Xanthine dehydrogenase | CHEMBL1929 | P47989 | 2E1Q | T40954 | 64% | 96% |
| Acyl-CoA desaturase | CHEMBL5555 | O00767 | 4ZYO | T10897 | 64% | 97.5% |
| Interleukin-1 receptor-associated kinase 1 | CHEMBL3357 | P51617 | 6BFN | Not Available | 64% | 78% |
| Tissue-type plasminogen activator | CHEMBL1873 | P00750 | 1RTF | T45299 | 63% | 93% |
| Histamine H3 receptor | CHEMBL264 | Q9Y5N1 | Not Available | Not Available | 63% | 91% |
| Cathepsin L | CHEMBL3837 | P07711 | 6JD8 | T41141 | 63% | 97% |
| Peptidyl-prolyl cis-trans isomerase NIMA-interacting 1 | CHEMBL2288 | Q13526 | 1PIN | T16308 | 63% | 92% |
| Histone acetyltransferase p300 | CHEMBL3784 | Q09472 | 6GYR | Not Available | 63% | 93% |
| Ghrelin receptor | CHEMBL4616 | Q92847 | 6KO5 | T59604 | 63% | 92% |
| Excitatory amino acid transporter 1 | CHEMBL3085 | P43003 | 5LM4 | Not Available | 62% | 95% |
| Protein kinase N1 | CHEMBL3384 | Q16512 | 4OTH | Not Available | 62% | 81% |
| Tissue factor pathway inhibitor | CHEMBL3713062 | P10646 | 5NMV | T78890 | 62% | 97% |
| PI3-kinase p110-beta subunit | CHEMBL3145 | P42338 | Not Available | T05031 | 61% | 99% |
| Transmembrane protease serine 6 | CHEMBL1795139 | Q8IU80 | Not Available | Not Available | 61.4% | 98% |
| Monoamine oxidase B | CHEMBL2039 | P27338 | 1S3E | Not Available | 61% | 93% |
| Nerve growth factor receptor Trk-A | CHEMBL2815 | P04629 | 2IFG | Not Available | 61% | 87% |
| 11-beta-hydroxysteroid dehydrogenase 2 | CHEMBL3746 | P80365 | Not Available | T43721 | 61% | 95% |
| Dopamine D2 receptor | CHEMBL217 | P14416 | 7JVR | Not Available | 61% | 96% |
| Lipoxin A4 receptor | CHEMBL4227 | P25090 | 6OMM | Not Available | 61% | 100% |
| Serine/threonine-protein kinase ULK3 | CHEMBL5047 | Q6PHR2 | 6FDY | Not Available | 61% | 78.5% |
| Endoplasmic reticulum-associated amyloid beta-peptide-binding protein | CHEMBL4159 | Q99714 | 2O23 | Not Available | 61% | 70% |
| Multidrug resistance-associated protein 1 | CHEMBL3004 | P33527 | 4C3Z | T11288 | 60.6% | 96% |
| Cannabinoid CB1 receptor | CHEMBL218 | P21554 | 6N4B | Not Available | 61% | 97% |
| Integrin alpha-5/beta-1 | CHEMBL2095226 | P05556 | 7NWL | T01851 | 60% | 96% |
| Pyroglutamylated RFamide peptide receptor | CHEMBL5852 | Q96P65 | Not Available | Not Available | 60% | 85% |
| dUTP pyrophosphatase | CHEMBL5203 | P33316 | 3ARA | T39054 | 60% | 99% |
| Arachidonate 5-lipoxygenase | CHEMBL215 | P09917 | 3V98 | Not Available | 59.8% | 93% |
| Adenosine A2b receptor | CHEMBL255 | P29275 | Not Available | T86679 | 60% | 99% |
| Vitamin K-dependent protein C | CHEMBL4444 | P04070 | 6M3B | T24836 | 59% | 94% |
| Hypoxanthine-guanine phosphoribosyltransferase | CHEMBL2360 | P00492 | 5HIA | Not Available | 59% | 87% |
| Proteasome component C5 | CHEMBL4208 | P20618 | 6KWY | Not Available | 59% | 90% |
| Excitatory amino acid transporter 3 | CHEMBL2721 | P43005 | 6X2L | Not Available | 58.7% | 93.5% |
| Neurotensin receptor 2 | CHEMBL2514 | O95665 | Not Available | Not Available | 59% | 100% |
| N-acylsphingosine-amidohydrolase | CHEMBL4349 | Q02083 | 6DXW | Not Available | 58% | 93% |
| Alkaline phosphatase, tissue-nonspecific isozyme | CHEMBL5979 | P05186 | Not Available | T09538 | 58% | 85.4% |
| Lysine-specific demethylase 5C | CHEMBL2163176 | P41229 | 5FWJ | Not Available | 58.1% | 93% |
| PI3-kinase p110-alpha/p85-alpha | CHEMBL2111367 | P27986 | 4JPS | T80276 | 58% | 94% |
| Non-receptor tyrosine-protein kinase TNK1 | CHEMBL5334 | Q13470 | Not Available | Not Available | 58% | 73% |
| WD repeat-containing protein 5 | CHEMBL1075317 | P61964 | 2GNQ | Not Available | 57.9% | 96% |
| Ectonucleotide pyrophosphatase/phosphodiesterase family member 1 | CHEMBL5925 | P22413 | 6WFJ | Not Available | 58% | 92% |
| Dual specificity protein kinase CLK4 | CHEMBL4203 | Q9HAZ1 | 6FYV | Not Available | 58% | 94% |
| Transcription intermediary factor 1-alpha | CHEMBL3108638 | O15164 | 4YBM | Not Available | 58% | 96% |
| Tyrosine-protein kinase Lyn | CHEMBL3905 | P07948 | 5XY1 | Not Available | 58% | 76% |
| Carbonic anhydrase III | CHEMBL2885 | P07451 | 3UYQ | Not Available | 58% | 87% |
| Anandamide amidohydrolase | CHEMBL2243 | O00519 | Not Available | T11754 | 57% | 98% |
| Tyrosine-protein kinase receptor UFO | CHEMBL4895 | P30530 | 5U6B | T82383 | 57% | 91% |
| Transient receptor potential cation channel subfamily M member 8 | CHEMBL1075319 | Q7Z2W7 | Not Available | T41955 | 57% | 79.5% |
| Serine/threonine-protein kinase Chk1 | CHEMBL4630 | O14757 | 3TKI | Not Available | 57% | 97% |
| Nuclear receptor subfamily 4 group A member 1 | CHEMBL1293229 | P22736 | 4RZF | Not Available | 56.6% | 78% |
| Glutaminase kidney isoform, mitochondrial | CHEMBL2146302 | O94925 | 3UO9 | T86734 | 56% | 100% |
| Histone deacetylase 2 | CHEMBL1937 | Q92769 | 7KBG | T51191 | 56% | 95% |
| Glycine transporter 1 | CHEMBL2337 | P48067 | 6ZBV | T69685 | 56% | 95% |
| Retinoid X receptor beta | CHEMBL1870 | P28702 | 7A78 | T60077 | 56% | 95% |
| Voltage-gated potassium channel subunit Kv1.5 | CHEMBL4306 | P22460 | Not Available | T17569 | 56% | 94% |
| Cyclooxygenase-1 | CHEMBL221 | P23219 | 6Y3C | Not Available | 56% | 90% |
| PI3-kinase p110-delta subunit | CHEMBL3130 | O00329 | 6PYR | T67849 | 55% | 96% |
| Tyrosine-protein kinase FRK | CHEMBL4223 | P42685 | Not Available | Not Available | 55% | 71% |
| FML2_HUMAN | CHEMBL5646 | Q6L5J4 | Not Available | Not Available | 55% | 100% |
| Uracil nucleotide/cysteinyl leukotriene receptor | CHEMBL1075162 | Q13304 | Not Available | Not Available | 55% | 80% |
| Free fatty acid receptor 2 | CHEMBL5493 | O15552 | Not Available | T28213 | 55% | 93% |
| Cysteinyl leukotriene receptor 2 | CHEMBL4330 | Q9NS75 | Not Available | T74238 | 55% | 98% |
| Sodium channel protein type II alpha subunit | CHEMBL4187 | Q99250 | 6J8E | Not Available | 55% | 95.5% |
| Thymidylate synthase | CHEMBL1952 | P04818 | 3GH0 | T98397 | 54% | 94% |
| Isocitrate dehydrogenase [NADP] cytoplasmic | CHEMBL2007625 | O75874 | 6BKX | T69563 | 54% | 99% |
| Ribosomal protein S6 kinase alpha 6 | CHEMBL4924 | Q9UK32 | 6G77 | Not Available | 54% | 80% |
| Casein kinase I epsilon | CHEMBL4937 | P49674 | 4HNI | Not Available | 54% | 93% |
| c-Jun N-terminal kinase 2 | CHEMBL4179 | P45984 | 3E7O | Not Available | 54% | 91% |
| Caspase-6 | CHEMBL3308 | P55212 | 2WDP | Not Available | 53.4% | 98% |
| Sodium channel protein type III alpha subunit | CHEMBL5163 | Q9NY46 | Not Available | T76937 | 53% | 96.9% |
| Sodium/hydrogen exchanger 1 | CHEMBL2781 | P19634 | 7DSX | T82028 | 53% | 90% |
| Lysine-specific demethylase 4C | CHEMBL6175 | Q9H3R0 | 5FJK | Not Available | 53% | 97% |
| Sphingosine 1-phosphate receptor Edg-8 | CHEMBL2274 | Q9H228 | Not Available | T50089 | 53% | 100% |
| Histone deacetylase 5 | CHEMBL2563 | Q9UQL6 | 5UWI | Not Available | 53% | 90% |
| Phosphodiesterase 3A | CHEMBL241 | Q14432 | 7LRC | T88975 | 53% | 93% |
| Neurokinin 2 receptor | CHEMBL2327 | P21452 | Not Available | T52790 | 53% | 99% |
| Prostanoid EP1 receptor | CHEMBL1811 | P34995 | Not Available | T15497 | 52% | 96% |
| Macrophage-stimulating protein receptor | CHEMBL2689 | Q04912 | 4QT8 | Not Available | 52% | 90% |
| Excitatory amino acid transporter 2 | CHEMBL4973 | P43004 | Not Available | Not Available | 52% | 99% |
| Voltage-gated N-type calcium channel alpha-1B subunit | CHEMBL4478 | Q00975 | Not Available | T38338 | 52% | 97% |
| Platelet-derived growth factor receptor | CHEMBL2095189 | P09619 | 3MJG | T53524 | 51.5% | 72% |
| Coagulation factor XIII | CHEMBL4530 | P00488 | 4KTY | Not Available | 51% | 96% |
| Nitric-oxide synthase, endothelial | CHEMBL4803 | P29474 | 4D1P | T06046 | 51% | 86% |
| Aminopeptidase N | CHEMBL1907 | P15144 | 4FYT | T67272 | 51% | 93% |
| Fructose-1,6-bisphosphatase | CHEMBL3975 | P09467 | 7C9Q | Not Available | 50% | 93% |
| Tyrosyl-DNA phosphodiesterase 2 | CHEMBL2169736 | O95551 | 5J3P | Not Available | 50% | 98% |
| Dipeptidyl peptidase VIII | CHEMBL4657 | Q6V1X1 | 6EOP | Not Available | 50% | 97% |
| Histone acetyltransferase KAT6A | CHEMBL3774298 | Q92794 | 2OZU | Not Available | 50% | 97.5% |

**Table S2. Predicted targets of Fluoxetine**

| **Target Name** | **ChEMBL-ID** | **UniProt ID** | **PDB Visualization** | **TTD ID** | **Probability** | **Accuracy** |
| --- | --- | --- | --- | --- | --- | --- |
| Nuclear factor NF-kappa-B p105 subunit | CHEMBL3251 | P19838 | 1SVC | Not Available | 95% | 96% |
| Cathepsin D | CHEMBL2581 | P07339 | 4OD9 | T67102 | 93% | 99% |
| Cyclooxygenase-1 | CHEMBL221 | P23219 | 6Y3C | Not Available | 92% | 90% |
| Glycine transporter 2 | CHEMBL3060 | Q9Y345 | Not Available | Not Available | 90% | 99% |
| Glycine transporter 1 | CHEMBL2337 | P48067 | 6ZBV | T69685 | 90% | 95% |
| Monoamine oxidase B | CHEMBL2039 | P27338 | 1S3E | Not Available | 90% | 93% |
| Muscarinic acetylcholine receptor M5 | CHEMBL2035 | P08912 | 6OL9 | T79961 | 89% | 95% |
| Transcription intermediary factor 1-alpha | CHEMBL3108638 | O15164 | 4YBM | Not Available | 88% | 96% |
| DNA-(apurinic or apyrimidinic site) lyase | CHEMBL5619 | P27695 | 6BOW | T13348 | 88% | 91% |
| Kruppel-like factor 5 | CHEMBL1293249 | Q13887 | Not Available | Not Available | 86% | 86% |
| Muscarinic acetylcholine receptor M1 | CHEMBL216 | P11229 | 6OIJ | T28893 | 85% | 94% |
| Serine/threonine-protein kinase ULK1 | CHEMBL6006 | O75385 | 4WNO | Not Available | 85% | 76% |
| Pregnane X receptor | CHEMBL3401 | O75469 | 6TFI | T82702 | 84% | 95% |
| Muscarinic acetylcholine receptor M4 | CHEMBL1821 | P08173 | 5DSG | T20709 | 84% | 94% |
| Signal transducer and activator of transcription 3 | CHEMBL4026 | P40763 | 6QHD | T29130 | 84% | 83% |
| Aminopeptidase N | CHEMBL1907 | P15144 | 4FYT | T67272 | 83.2% | 93% |
| Nuclear factor erythroid 2-related factor 2 | CHEMBL1075094 | Q16236 | 2FLU | Not Available | 83% | 96% |
| Free fatty acid receptor 1 | CHEMBL4422 | O14842 | 5TZR | Not Available | 81% | 93% |
| Lysosomal Pro-X carboxypeptidase | CHEMBL2335 | P42785 | 3N2Z | Not Available | 81% | 100% |
| Glucose transporter | CHEMBL2535 | P11166 | 6THA | Not Available | 81% | 99% |
| Nitric oxide synthase, inducible | CHEMBL4481 | P35228 | 3E7G | T02703 | 80% | 94.8% |
| Kinesin-like protein 1 | CHEMBL4581 | P52732 | 6TIW | T28484 | 80% | 93% |
| Thyroid hormone receptor alpha | CHEMBL1860 | P10827 | 3ILZ | T79591 | 79% | 99% |
| Histamine H3 receptor | CHEMBL264 | Q9Y5N1 | Not Available | Not Available | 79% | 91% |
| Cytochrome P450 2A6 | CHEMBL5282 | P11509 | 2FDV | T06455 | 77% | 72% |
| Nuclear receptor ROR-beta | CHEMBL3091268 | Q92753 | Not Available | Not Available | 77% | 95.5% |
| Adenosine A2b receptor | CHEMBL255 | P29275 | Not Available | T86679 | 77% | 99% |
| Muscarinic acetylcholine receptor M2 | CHEMBL211 | P08172 | 5ZKC | T46185 | 76% | 95% |
| Glutathione S-transferase Pi | CHEMBL3902 | P09211 | 5J41 | T21669 | 76% | 94% |
| C-C chemokine receptor type 2 | CHEMBL4015 | P41597 | 5T1A | T89988 | 75% | 99% |
| Platelet-derived growth factor receptor alpha | CHEMBL2007 | P16234 | 7LBF | T53524 | 75% | 91% |
| Histone-arginine methyltransferase CARM1 | CHEMBL5406 | Q86X55 | 6DVR | T12837 | 74% | 94% |
| Beta-glucosidase | CHEMBL3761 | Q9HCG7 | Not Available | Not Available | 74% | 99% |
| Tyrosyl-DNA phosphodiesterase 1 | CHEMBL1075138 | Q9NUW8 | 6N0D | Not Available | 74% | 71% |
| Dual specificity phosphatase Cdc25C | CHEMBL2378 | P30307 | 3OP3 | Not Available | 74% | 97% |
| Beta-glucuronidase | CHEMBL2728 | P08236 | 3HN3 | T96413 | 72.6% | 78% |
| ADAM10 | CHEMBL5028 | O14672 | 6BE6 | T31902 | 73% | 97.5% |
| Proteasome component C5 | CHEMBL4208 | P20618 | 6KWY | Not Available | 72% | 90% |
| Excitatory amino acid transporter 1 | CHEMBL3085 | P43003 | 5LM4 | Not Available | 71% | 95% |
| G-protein coupled bile acid receptor 1 | CHEMBL5409 | Q8TDU6 | 7CFM | T86273 | 70.2% | 94% |
| Voltage-gated N-type calcium channel alpha-1B subunit | CHEMBL4478 | Q00975 | Not Available | T38338 | 70% | 97% |
| Bcr/Abl fusion protein | CHEMBL2096618 | P00519 | 5N7E | Not Available | 70% | 86% |
| Tyrosine-protein kinase Lyn | CHEMBL3905 | P07948 | 5XY1 | Not Available | 68% | 76% |
| DCN1-like protein 1 | CHEMBL4105838 | Q96GG9 | 6BG3 | Not Available | 67% | 95% |
| GABA-A receptor; alpha-1/beta-2/gamma-2 | CHEMBL2095172 | P14867 | 6X3T | T51487 | 67% | 93% |
| Sodium channel protein type V alpha subunit | CHEMBL1980 | Q14524 | 6LQA | T39716 | 67% | 92.5% |
| Galectin-3 | CHEMBL4531 | P17931 | 6FOF | T72038 | 66% | 96.9% |
| Glutamate NMDA receptor; GRIN1/GRIN2B | CHEMBL1907603 | Q05586 | 5EWM | Not Available | 66% | 96% |
| Dual specificity protein phosphatase 3 | CHEMBL2635 | P51452 | 3F81 | Not Available | 66% | 94% |
| NT-3 growth factor receptor | CHEMBL5608 | Q16288 | 6KZD | Not Available | 66% | 96% |
| Rho-associated protein kinase 1 | CHEMBL3231 | Q13464 | 3V8S | T51282 | 66% | 96% |
| E3 SUMO-protein ligase CBX4 | CHEMBL3232685 | O00257 | 5EPL | Not Available | 65% | 93% |
| Histone deacetylase 5 | CHEMBL2563 | Q9UQL6 | 5UWI | Not Available | 65% | 90% |
| Tyrosine-protein kinase ITK/TSK | CHEMBL2959 | Q08881 | 4HCU | T91761 | 65% | 95% |
| Tyrosine-protein kinase FYN | CHEMBL1841 | P06241 | 2DQ7 | T17980 | 65% | 81% |
| cAMP-dependent protein kinase alpha-catalytic subunit | CHEMBL4101 | P17612 | 4WB8 | Not Available | 65% | 83% |
| Tyrosine-protein kinase YES | CHEMBL2073 | P07947 | 2HDA | Not Available | 64% | 83% |
| Neuronal acetylcholine receptor; alpha3/beta4 | CHEMBL1907594 | P30926 | 6PV7 | T73724 | 64% | 97% |
| Excitatory amino acid transporter 2 | CHEMBL4973 | P43004 | Not Available | Not Available | 64% | 99% |
| TGF-beta receptor type II | CHEMBL4267 | P37173 | 5QIN | T49989 | 64% | 88% |
| Neuronal acetylcholine receptor; alpha4/beta4 | CHEMBL1907591 | P30926 | 6UR8 | T70967 | 63.8% | 100% |
| Amine oxidase, copper containing | CHEMBL3437 | Q16853 | 2C10 | T69619 | 63% | 94% |
| Dual specificity phosphatase Cdc25B | CHEMBL4804 | P30305 | 1QB0 | Not Available | 63% | 79.5% |
| Neurokinin 1 receptor | CHEMBL249 | P25103 | 6HLP | T47094 | 63% | 99% |
| Cyclin-dependent kinase 7 | CHEMBL3055 | P50613 | 7B5O | T58449 | 62% | 82% |
| Calpain 1 | CHEMBL3891 | P07384 | 1ZCM | Not Available | 62% | 93% |
| Serine/threonine-protein kinase PLK1 | CHEMBL3024 | P53350 | 2OWB | T40694 | 62% | 97% |
| Cysteinyl leukotriene receptor 2 | CHEMBL4330 | Q9NS75 | Not Available | T74238 | 61% | 98% |
| Protein Mdm4 | CHEMBL1255126 | O15151 | 6Q9Y | T36741 | 60.8% | 90.2% |
| Sodium channel protein type III alpha subunit | CHEMBL5163 | Q9NY46 | Not Available | T76937 | 61% | 96.9% |
| Peroxisome proliferator-activated receptor delta | CHEMBL3979 | Q03181 | 5U3Q | T36557 | 61% | 94% |
| Telomerase reverse transcriptase | CHEMBL2916 | O14746 | 7BG9 | T86052 | 60% | 90% |
| G-protein coupled receptor 55 | CHEMBL1075322 | Q9Y2T6 | Not Available | T87670 | 60% | 78% |
| LSD1/CoREST complex | CHEMBL3137262 | O60341 | 5L3D | Not Available | 58% | 97% |
| Cystic fibrosis transmembrane conductance regulator | CHEMBL4051 | P13569 | 6MSM | T55654 | 57.8% | 96% |
| Excitatory amino acid transporter 3 | CHEMBL2721 | P43005 | 6X2L | Not Available | 57% | 93.5% |
| Vascular endothelial growth factor receptor 1 | CHEMBL1868 | P17948 | 5T89 | Not Available | 56% | 96% |
| Cannabinoid CB1 receptor | CHEMBL218 | P21554 | 6N4B | Not Available | 56% | 97% |
| 5'-nucleotidase | CHEMBL5957 | P21589 | 6TVE | Not Available | 56% | 98% |
| Tyrosine-protein kinase FRK | CHEMBL4223 | P42685 | Not Available | Not Available | 56% | 71% |
| Beta-glucocerebrosidase | CHEMBL2179 | P04062 | 6TN1 | T84173 | 56% | 85% |
| Trypsin I | CHEMBL209 | P07477 | 2RA3 | T27602 | 56% | 90% |
| Signal transducer and activator of transcription 1-alpha/beta | CHEMBL6101 | P42224 | 1YVL | T64205 | 56% | 73% |
| MAP kinase p38 beta | CHEMBL3961 | Q15759 | 3GP0 | Not Available | 55.7% | 95% |
| C5a anaphylatoxin chemotactic receptor | CHEMBL2373 | P21730 | 6C1R | T15439 | 56% | 93% |
| Glycine receptor subunit alpha-1 | CHEMBL5845 | P23415 | 4X5T | T50269 | 55% | 91% |
| Lipoxin A4 receptor | CHEMBL4227 | P25090 | 6OMM | Not Available | 55.4% | 100% |
| Ephrin type-B receptor 2 | CHEMBL3290 | P29323 | 3ZFM | T73756 | 55% | 78% |
| Proteasome Macropain subunit | CHEMBL3492 | P49721 | 5LE5 | Not Available | 55% | 90% |
| Histone-lysine N-methyltransferase, H3 lysine-79 specific | CHEMBL1795117 | Q8TEK3 | 3UWP | Not Available | 55% | 94% |
| Hypoxanthine-guanine phosphoribosyltransferase | CHEMBL2360 | P00492 | 5HIA | Not Available | 55% | 87% |
| Acetylcholine receptor; alpha1/beta1/delta/gamma | CHEMBL1907588 | P02708 | 5HBT | T04689 | 55% | 98% |
| PI3-kinase p110-delta subunit | CHEMBL3130 | O00329 | 6PYR | T67849 | 55% | 96% |
| Adenosylhomocysteinase | CHEMBL2664 | P23526 | 1LI4 | Not Available | 54% | 87% |
| Serine/threonine protein kinase NLK | CHEMBL5364 | Q9UBE8 | Not Available | Not Available | 54% | 79% |
| Apelin receptor | CHEMBL1628481 | P35414 | 5VBL | T65783 | 54% | 98% |
| Multidrug resistance-associated protein 1 | CHEMBL3004 | P33527 | 4C3Z | T11288 | 54% | 96% |
| Acyl coenzyme A:cholesterol acyltransferase 1 | CHEMBL2782 | P35610 | 6P2J | Not Available | 53% | 92% |
| Cannabinoid CB2 receptor | CHEMBL253 | P34972 | 6KPF | Not Available | 53% | 97% |
| Mitogen-activated protein kinase kinase kinase 5 | CHEMBL5285 | Q99683 | 5ULM | T97589 | 53% | 92% |
| Mitogen-activated protein kinase kinase kinase kinase 2 | CHEMBL5330 | Q12851 | Not Available | Not Available | 52.5% | 79% |
| Sphingosine 1-phosphate receptor Edg-8 | CHEMBL2274 | Q9H228 | Not Available | T50089 | 52% | 100% |
| Choline kinase alpha | CHEMBL3117 | P35790 | 4DA5 | T55709 | 52.1% | 78% |
| Acyl-CoA desaturase | CHEMBL5555 | O00767 | 4ZYO | T10897 | 52% | 97.5% |
| Transient receptor potential cation channel subfamily A member 1 | CHEMBL6007 | O75762 | 6V9V | T84040 | 51% | 92% |
| Monoamine oxidase A | CHEMBL1951 | P21397 | 2Z5Y | Not Available | 51% | 91% |
| Plasminogen activator inhibitor-1 | CHEMBL3475 | P05121 | 3CVM | T15556 | 51% | 83% |
| Protein kinase C delta | CHEMBL2996 | Q05655 | 1YRK | T44861 | 51% | 98% |
| Endoplasmic reticulum aminopeptidase 1 | CHEMBL5939 | Q9NZ08 | 6Q4R | Not Available | 51.1% | 100% |
| Melanin-concentrating hormone receptor 1 | CHEMBL344 | Q99705 | Not Available | T09572 | 51% | 92.5% |
| Adenosine A1 receptor | CHEMBL226 | P30542 | 5N2S | T92072 | 51% | 96% |
| Kallikrein 1 | CHEMBL2319 | P06870 | 1SPJ | T40000 | 50% | 91% |
| Peptidyl-prolyl cis-trans isomerase FKBP5 | CHEMBL2052031 | Q13451 | 5OMP | Not Available | 50% | 88% |
| Non-receptor tyrosine-protein kinase TNK1 | CHEMBL5334 | Q13470 | Not Available | Not Available | 50% | 73% |

**Table S3. Predicted targets of Fluvoxamine**

| **Target Name** | **ChEMBL-ID** | **UniProt ID** | **PDB Visualization** | **TTD ID** | **Probability** | **Accuracy** |
| --- | --- | --- | --- | --- | --- | --- |
| Peroxisome proliferator-activated receptor alpha | CHEMBL239 | Q07869 | 6KAX | T86591 | 98% | 91% |
| Nuclear factor NF-kappa-B p105 subunit | CHEMBL3251 | P19838 | 1SVC | Not Available | 97% | 96% |
| Sphingosine 1-phosphate receptor Edg-6 | CHEMBL3230 | O95977 | Not Available | Not Available | 97% | 94% |
| Endoplasmic reticulum-associated amyloid beta-peptide-binding protein | CHEMBL4159 | Q99714 | 2O23 | Not Available | 96% | 70% |
| Glycine transporter 2 | CHEMBL3060 | Q9Y345 | Not Available | Not Available | 95% | 99% |
| Monoamine oxidase B | CHEMBL2039 | P27338 | 1S3E | Not Available | 95% | 93% |
| Kinesin-like protein 1 | CHEMBL4581 | P52732 | 6TIW | T28484 | 94% | 93% |
| DNA-(apurinic or apyrimidinic site) lyase | CHEMBL5619 | P27695 | 6BOW | T13348 | 93% | 91% |
| Tyrosyl-DNA phosphodiesterase 1 | CHEMBL1075138 | Q9NUW8 | 6N0D | Not Available | 91.7% | 71% |
| Kruppel-like factor 5 | CHEMBL1293249 | Q13887 | Not Available | Not Available | 91% | 86% |
| Proteasome Macropain subunit | CHEMBL3492 | P49721 | 5LE5 | Not Available | 91% | 90% |
| Cannabinoid CB2 receptor | CHEMBL253 | P34972 | 6KPF | Not Available | 90% | 97% |
| Platelet-derived growth factor receptor | CHEMBL2095189 | P09619 | 3MJG | T53524 | 87% | 72% |
| Serotonin 2c (5-HT2c) receptor | CHEMBL225 | P28335 | 6BQH | T83813 | 87% | 90% |
| Bcr/Abl fusion protein | CHEMBL2096618 | P00519 | 5N7E | Not Available | 85% | 86% |
| Sphingosine 1-phosphate receptor Edg-8 | CHEMBL2274 | Q9H228 | Not Available | T50089 | 84% | 100% |
| Carbonic anhydrase III | CHEMBL2885 | P07451 | 3UYQ | Not Available | 83% | 87% |
| Arachidonate 5-lipoxygenase | CHEMBL215 | P09917 | 3V98 | Not Available | 82.9% | 93% |
| Hexokinase type IV | CHEMBL3820 | P35557 | 3F9M | T87166 | 81.9% | 92% |
| C-C chemokine receptor type 2 | CHEMBL4015 | P41597 | 5T1A | T89988 | 82% | 99% |
| Melanin-concentrating hormone receptor 1 | CHEMBL344 | Q99705 | Not Available | T09572 | 81.4% | 92.5% |
| Proteasome component C5 | CHEMBL4208 | P20618 | 6KWY | Not Available | 81% | 90% |
| Nuclear factor erythroid 2-related factor 2 | CHEMBL1075094 | Q16236 | 2FLU | Not Available | 80% | 96% |
| Sodium channel protein type III alpha subunit | CHEMBL5163 | Q9NY46 | Not Available | T76937 | 79% | 96.9% |
| Methionine aminopeptidase 2 | CHEMBL3922 | P50579 | 1B6A | T75596 | 79% | 97% |
| Transcription intermediary factor 1-alpha | CHEMBL3108638 | O15164 | 4YBM | Not Available | 79% | 96% |
| Glycine receptor subunit alpha-1 | CHEMBL5845 | P23415 | 4X5T | T50269 | 79% | 91% |
| Cytochrome P450 2A6 | CHEMBL5282 | P11509 | 2FDV | T06455 | 78% | 72% |
| Pregnane X receptor | CHEMBL3401 | O75469 | 6TFI | T82702 | 78% | 95% |
| E3 SUMO-protein ligase CBX4 | CHEMBL3232685 | O00257 | 5EPL | Not Available | 77% | 93% |
| PI3-kinase p110-beta subunit | CHEMBL3145 | P42338 | Not Available | T05031 | 77% | 99% |
| Aminopeptidase N | CHEMBL1907 | P15144 | 4FYT | T67272 | 76% | 93% |
| Protein Mdm4 | CHEMBL1255126 | O15151 | 6Q9Y | T36741 | 76% | 90.2% |
| NT-3 growth factor receptor | CHEMBL5608 | Q16288 | 6KZD | Not Available | 74% | 96% |
| Macrophage migration inhibitory factor | CHEMBL2085 | P14174 | 6B1K | T39977 | 74% | 81% |
| Platelet-derived growth factor receptor alpha | CHEMBL2007 | P16234 | 7LBF | T53524 | 73% | 91% |
| ADAM10 | CHEMBL5028 | O14672 | 6BE6 | T31902 | 72% | 97.5% |
| Glucose transporter | CHEMBL2535 | P11166 | 6THA | Not Available | 72% | 99% |
| Neurokinin 1 receptor | CHEMBL249 | P25103 | 6HLP | T47094 | 72% | 99% |
| Histone deacetylase 8 | CHEMBL3192 | Q9BY41 | 5VI6 | T28887 | 72% | 94% |
| Chromobox protein homolog 7 | CHEMBL1764946 | O95931 | 4MN3 | Not Available | 71% | 97% |
| Calpain 1 | CHEMBL3891 | P07384 | 1ZCM | Not Available | 71% | 93% |
| Cysteinyl leukotriene receptor 2 | CHEMBL4330 | Q9NS75 | Not Available | T74238 | 70% | 98% |
| Lysosomal Pro-X carboxypeptidase | CHEMBL2335 | P42785 | 3N2Z | Not Available | 70% | 100% |
| LSD1/CoREST complex | CHEMBL3137262 | O60341 | 5L3D | Not Available | 69% | 97% |
| Dual specificity protein phosphatase 3 | CHEMBL2635 | P51452 | 3F81 | Not Available | 69% | 94% |
| Bloom syndrome protein | CHEMBL1293237 | P54132 | 4O3M | Not Available | 68% | 70% |
| Thymidine phosphorylase | CHEMBL3106 | P19971 | 1UOU | T59929 | 67% | 77% |
| Mineralocorticoid receptor | CHEMBL1994 | P08235 | 4PF3 | Not Available | 67% | 100% |
| Rho-associated protein kinase 1 | CHEMBL3231 | Q13464 | 3V8S | T51282 | 66% | 96% |
| Histamine H3 receptor | CHEMBL264 | Q9Y5N1 | Not Available | Not Available | 66% | 91% |
| Tyrosine-protein kinase FRK | CHEMBL4223 | P42685 | Not Available | Not Available | 66% | 71% |
| Serotonin 2a (5-HT2a) receptor | CHEMBL224 | P28223 | 6WHA | T32060 | 65% | 91% |
| Acyl coenzyme A:cholesterol acyltransferase | CHEMBL2265 | P23141 | 5A7G | T76369 | 64% | 86% |
| G-protein coupled receptor 35 | CHEMBL1293267 | Q9HC97 | Not Available | Not Available | 64% | 89% |
| Phosphodiesterase 3A | CHEMBL241 | Q14432 | 7LRC | T88975 | 64% | 93% |
| MAP kinase p38 beta | CHEMBL3961 | Q15759 | 3GP0 | Not Available | 63% | 95% |
| G-protein coupled bile acid receptor 1 | CHEMBL5409 | Q8TDU6 | 7CFM | T86273 | 62% | 94% |
| Prostanoid EP1 receptor | CHEMBL1811 | P34995 | Not Available | T15497 | 62% | 96% |
| Lipoxin A4 receptor | CHEMBL4227 | P25090 | 6OMM | Not Available | 62% | 100% |
| Vascular endothelial growth factor receptor 2 | CHEMBL279 | P35968 | 3V2A | T80975 | 62% | 96% |
| Mitogen-activated protein kinase kinase kinase kinase 2 | CHEMBL5330 | Q12851 | Not Available | Not Available | 62% | 79% |
| Neuronal acetylcholine receptor; alpha4/beta4 | CHEMBL1907591 | P30926 | 6UR8 | T70967 | 61% | 100% |
| Cathepsin D | CHEMBL2581 | P07339 | 4OD9 | T67102 | 61% | 99% |
| Excitatory amino acid transporter 1 | CHEMBL3085 | P43003 | 5LM4 | Not Available | 61% | 95% |
| Tissue factor pathway inhibitor | CHEMBL3713062 | P10646 | 5NMV | T78890 | 60% | 97% |
| C5a anaphylatoxin chemotactic receptor | CHEMBL2373 | P21730 | 6C1R | T15439 | 60% | 93% |
| Kelch-like ECH-associated protein 1 | CHEMBL2069156 | Q14145 | 6WCQ | Not Available | 59.8% | 82% |
| Interleukin-23 receptor | CHEMBL4296013 | Q5VWK5 | 5MZV | T00484 | 60% | 88% |
| Uracil nucleotide/cysteinyl leukotriene receptor | CHEMBL1075162 | Q13304 | Not Available | Not Available | 59% | 80% |
| Excitatory amino acid transporter 3 | CHEMBL2721 | P43005 | 6X2L | Not Available | 59% | 93.5% |
| Acyl-CoA desaturase | CHEMBL5555 | O00767 | 4ZYO | T10897 | 59.3% | 97.5% |
| Amine oxidase, copper containing | CHEMBL3437 | Q16853 | 2C10 | T69619 | 59% | 94% |
| Kallikrein 7 | CHEMBL2443 | P49862 | 2QXI | Not Available | 59% | 94% |
| Cannabinoid CB1 receptor | CHEMBL218 | P21554 | 6N4B | Not Available | 59% | 97% |
| LDL-associated phospholipase A2 | CHEMBL3514 | Q13093 | 3D59 | T69912 | 58% | 97.5% |
| cAMP-dependent protein kinase alpha-catalytic subunit | CHEMBL4101 | P17612 | 4WB8 | Not Available | 57% | 83% |
| Ephrin type-B receptor 2 | CHEMBL3290 | P29323 | 3ZFM | T73756 | 56% | 78% |
| Sphingosine 1-phosphate receptor Edg-3 | CHEMBL3892 | Q99500 | 7C4S | T11241 | 56% | 97% |
| Fructose-1,6-bisphosphatase | CHEMBL3975 | P09467 | 7C9Q | Not Available | 56% | 93% |
| Excitatory amino acid transporter 2 | CHEMBL4973 | P43004 | Not Available | Not Available | 56% | 99% |
| Coagulation factor XIII | CHEMBL4530 | P00488 | 4KTY | Not Available | 56% | 96% |
| Transthyretin | CHEMBL3194 | P02766 | 6SUG | T86462 | 56% | 91% |
| Peroxisome proliferator-activated receptor delta | CHEMBL3979 | Q03181 | 5U3Q | T36557 | 56% | 94% |
| P2X purinoceptor 7 | CHEMBL4805 | Q99572 | Not Available | T63414 | 55.3% | 97.5% |
| Sodium/hydrogen exchanger 1 | CHEMBL2781 | P19634 | 7DSX | T82028 | 55% | 90% |
| Pyroglutamylated RFamide peptide receptor | CHEMBL5852 | Q96P65 | Not Available | Not Available | 55% | 85% |
| Anandamide amidohydrolase | CHEMBL2243 | O00519 | Not Available | T11754 | 54% | 98% |
| PI3-kinase p110-alpha/p85-alpha | CHEMBL2111367 | P27986 | 4JPS | T80276 | 54% | 94% |
| Alkaline phosphatase, tissue-nonspecific isozyme | CHEMBL5979 | P05186 | Not Available | T09538 | 54% | 85.4% |
| Muscarinic acetylcholine receptor M2 | CHEMBL211 | P08172 | 5ZKC | T46185 | 54% | 95% |
| Cyclin-dependent kinase 2/cyclin E1 | CHEMBL1907605 | P24864 | 1W98 | T70176 | 54% | 93% |
| Muscarinic acetylcholine receptor M4 | CHEMBL1821 | P08173 | 5DSG | T20709 | 54% | 94% |
| DNA topoisomerase I | CHEMBL1781 | P11387 | 1K4T | T09826 | 54% | 97% |
| LXR-alpha | CHEMBL2808 | Q13133 | 5HJS | T52297 | 53% | 97% |
| Trypsin I | CHEMBL209 | P07477 | 2RA3 | T27602 | 53% | 90% |
| Neurotensin receptor 2 | CHEMBL2514 | O95665 | Not Available | Not Available | 53% | 100% |
| Muscarinic acetylcholine receptor M5 | CHEMBL2035 | P08912 | 6OL9 | T79961 | 53% | 95% |
| Ectonucleotide pyrophosphatase/phosphodiesterase family member 1 | CHEMBL5925 | P22413 | 6WFJ | Not Available | 53% | 92% |
| Apelin receptor | CHEMBL1628481 | P35414 | 5VBL | T65783 | 53% | 98% |
| G-protein coupled receptor 120 | CHEMBL5339 | Q5NUL3 | Not Available | Not Available | 52% | 96% |
| Purinergic receptor P2Y12 | CHEMBL2001 | Q9H244 | 4PXZ | T46937 | 52% | 96% |
| Glutamate NMDA receptor; GRIN1/GRIN2B | CHEMBL1907603 | Q05586 | 5EWM | Not Available | 51% | 96% |
| Multidrug resistance-associated protein 1 | CHEMBL3004 | P33527 | 4C3Z | T11288 | 50% | 96% |

**Table S4. Predicted targets of Paroxetine**

| **Target Name** | **ChEMBL-ID** | **UniProt ID** | **PDB Visualization** | **TTD ID** | **Probability** | **Accuracy** |
| --- | --- | --- | --- | --- | --- | --- |
| LSD1/CoREST complex | CHEMBL3137262 | O60341 | 5L3D | Not Available | 100% | 97% |
| HERG | CHEMBL240 | Q12809 | 5VA1 | T20251 | 99% | 90% |
| Nitric oxide synthase, inducible | CHEMBL4481 | P35228 | 3E7G | T02703 | 99% | 94.8% |
| Tyrosyl-DNA phosphodiesterase 1 | CHEMBL1075138 | Q9NUW8 | 6N0D | Not Available | 98% | 71% |
| DNA-(apurinic or apyrimidinic site) lyase | CHEMBL5619 | P27695 | 6BOW | T13348 | 98% | 91% |
| Nuclear factor NF-kappa-B p105 subunit | CHEMBL3251 | P19838 | 1SVC | Not Available | 95% | 96% |
| Kinesin-like protein 1 | CHEMBL4581 | P52732 | 6TIW | T28484 | 95% | 93% |
| Histone deacetylase 9 | CHEMBL4145 | Q9UKV0 | Not Available | Not Available | 93.8% | 85% |
| Monoamine oxidase B | CHEMBL2039 | P27338 | 1S3E | Not Available | 93% | 93% |
| NT-3 growth factor receptor | CHEMBL5608 | Q16288 | 6KZD | Not Available | 92% | 96% |
| dUTP pyrophosphatase | CHEMBL5203 | P33316 | 3ARA | T39054 | 92% | 99% |
| Serotonin 2c (5-HT2c) receptor | CHEMBL225 | P28335 | 6BQH | T83813 | 92% | 90% |
| Histone deacetylase 8 | CHEMBL3192 | Q9BY41 | 5VI6 | T28887 | 91% | 94% |
| Anandamide amidohydrolase | CHEMBL2243 | O00519 | Not Available | T11754 | 91% | 98% |
| Heat shock protein HSP 90-beta | CHEMBL4303 | P08238 | 5FWK | Not Available | 90% | 97% |
| Beta-glucuronidase | CHEMBL2728 | P08236 | 3HN3 | T96413 | 90% | 78% |
| C5a anaphylatoxin chemotactic receptor | CHEMBL2373 | P21730 | 6C1R | T15439 | 90% | 93% |
| Histone deacetylase 5 | CHEMBL2563 | Q9UQL6 | 5UWI | Not Available | 90% | 90% |
| Glutamate NMDA receptor; GRIN1/GRIN2B | CHEMBL1907603 | Q05586 | 5EWM | Not Available | 89% | 96% |
| Tyrosine-protein kinase BRK | CHEMBL4601 | Q13882 | 6CZ4 | T73694 | 88% | 76% |
| Glucose transporter | CHEMBL2535 | P11166 | 6THA | Not Available | 87% | 99% |
| Glycine transporter 2 | CHEMBL3060 | Q9Y345 | Not Available | Not Available | 87% | 99% |
| Protein kinase C zeta | CHEMBL3438 | Q05513 | Not Available | T59190 | 86% | 88% |
| Proteasome component C5 | CHEMBL4208 | P20618 | 6KWY | Not Available | 86% | 90% |
| Kruppel-like factor 5 | CHEMBL1293249 | Q13887 | Not Available | Not Available | 86% | 86% |
| G-protein coupled receptor 120 | CHEMBL5339 | Q5NUL3 | Not Available | Not Available | 85% | 96% |
| Cannabinoid CB2 receptor | CHEMBL253 | P34972 | 6KPF | Not Available | 85% | 97% |
| Galectin-3 | CHEMBL4531 | P17931 | 6FOF | T72038 | 85% | 96.9% |
| Mineralocorticoid receptor | CHEMBL1994 | P08235 | 4PF3 | Not Available | 85% | 100% |
| Lysosomal Pro-X carboxypeptidase | CHEMBL2335 | P42785 | 3N2Z | Not Available | 84% | 100% |
| Serotonin 7 (5-HT7) receptor | CHEMBL3155 | P34969 | Not Available | T79062 | 84% | 91% |
| Cathepsin D | CHEMBL2581 | P07339 | 4OD9 | T67102 | 84% | 99% |
| Neuronal acetylcholine receptor; alpha4/beta2 | CHEMBL1907589 | P17787 | 6UR8 | T70967 | 83.9% | 95% |
| Dual specificity phosphatase Cdc25B | CHEMBL4804 | P30305 | 1QB0 | Not Available | 83% | 79.5% |
| Protein tyrosine kinase 2 beta | CHEMBL5469 | Q14289 | 4EKU | T07087 | 83.2% | 91% |
| Dual specificity protein kinase CLK4 | CHEMBL4203 | Q9HAZ1 | 6FYV | Not Available | 81.9% | 94% |
| Histone deacetylase 7 | CHEMBL2716 | Q8WUI4 | 3C10 | Not Available | 82% | 89% |
| Glutathione S-transferase Pi | CHEMBL3902 | P09211 | 5J41 | T21669 | 82% | 94% |
| 5'-nucleotidase | CHEMBL5957 | P21589 | 6TVE | Not Available | 81% | 98% |
| Dihydroorotate dehydrogenase | CHEMBL1966 | Q02127 | 6FMD | T99009 | 81% | 96% |
| Lysine-specific demethylase 4C | CHEMBL6175 | Q9H3R0 | 5FJK | Not Available | 81% | 97% |
| Peroxisome proliferator-activated receptor alpha | CHEMBL239 | Q07869 | 6KAX | T86591 | 81% | 91% |
| DNA topoisomerase II alpha | CHEMBL1806 | P11388 | 6ZY5 | T17048 | 80% | 89% |
| Uracil nucleotide/cysteinyl leukotriene receptor | CHEMBL1075162 | Q13304 | Not Available | Not Available | 80% | 80% |
| Dual specificity protein phosphatase 3 | CHEMBL2635 | P51452 | 3F81 | Not Available | 80% | 94% |
| Sodium channel protein type IV alpha subunit | CHEMBL2072 | P35499 | 6AGF | T02546 | 79% | 92% |
| Thyroid hormone receptor alpha | CHEMBL1860 | P10827 | 3ILZ | T79591 | 78% | 99% |
| Kallikrein 1 | CHEMBL2319 | P06870 | 1SPJ | T40000 | 78% | 91% |
| Protein kinase N1 | CHEMBL3384 | Q16512 | 4OTH | Not Available | 78% | 81% |
| ALK tyrosine kinase receptor | CHEMBL4247 | Q9UM73 | 4Z55 | T56418 | 78% | 97% |
| Cysteine protease ATG4B | CHEMBL1741221 | Q9Y4P1 | 2CY7 | Not Available | 77% | 87.5% |
| Nuclear factor erythroid 2-related factor 2 | CHEMBL1075094 | Q16236 | 2FLU | Not Available | 77% | 96% |
| Proteasome Macropain subunit | CHEMBL3492 | P49721 | 5LE5 | Not Available | 76% | 90% |
| Dual specificity protein kinase CLK2 | CHEMBL4225 | P49760 | 6KHE | Not Available | 75.2% | 81% |
| Phosphodiesterase 3A | CHEMBL241 | Q14432 | 7LRC | T88975 | 75% | 93% |
| Coagulation factor XIII | CHEMBL4530 | P00488 | 4KTY | Not Available | 74% | 96% |
| Histone-arginine methyltransferase CARM1 | CHEMBL5406 | Q86X55 | 6DVR | T12837 | 74% | 94% |
| Inhibitor of nuclear factor kappa B kinase alpha subunit | CHEMBL3476 | O15111 | 5EBZ | Not Available | 74% | 96% |
| Adenosine A1 receptor | CHEMBL226 | P30542 | 5N2S | T92072 | 74% | 96% |
| Histamine H3 receptor | CHEMBL264 | Q9Y5N1 | Not Available | Not Available | 73.3% | 91% |
| Cyclin-dependent kinase 2/cyclin E1 | CHEMBL1907605 | P24864 | 1W98 | T70176 | 73% | 93% |
| Serine/threonine-protein kinase ULK1 | CHEMBL6006 | O75385 | 4WNO | Not Available | 72% | 76% |
| Transcription intermediary factor 1-alpha | CHEMBL3108638 | O15164 | 4YBM | Not Available | 72% | 96% |
| Tyrosine-protein kinase Lyn | CHEMBL3905 | P07948 | 5XY1 | Not Available | 72% | 76% |
| cAMP-dependent protein kinase alpha-catalytic subunit | CHEMBL4101 | P17612 | 4WB8 | Not Available | 71% | 83% |
| Acetyl-CoA carboxylase 1 | CHEMBL3351 | Q13085 | 6G2H | Not Available | 71% | 93% |
| Beta-1 adrenergic receptor | CHEMBL213 | P08588 | 7BVQ | T44068 | 71% | 96% |
| Sodium/hydrogen exchanger 1 | CHEMBL2781 | P19634 | 7DSX | T82028 | 71% | 90% |
| Peptidyl-prolyl cis-trans isomerase NIMA-interacting 1 | CHEMBL2288 | Q13526 | 1PIN | T16308 | 71% | 92% |
| Glutamate receptor ionotropic, AMPA 2 | CHEMBL4016 | P42262 | 2WJW | T42392 | 70% | 87% |
| Signal transducer and activator of transcription 3 | CHEMBL4026 | P40763 | 6QHD | T29130 | 70% | 83% |
| N-acylsphingosine-amidohydrolase | CHEMBL4349 | Q02083 | 6DXW | Not Available | 69.4% | 93% |
| Arachidonate 12-lipoxygenase | CHEMBL3687 | P18054 | 3D3L | Not Available | 69.2% | 76% |
| Glycine receptor subunit alpha-1 | CHEMBL5845 | P23415 | 4X5T | T50269 | 69% | 91% |
| Lysine-specific demethylase 4A | CHEMBL5896 | O75164 | 6G5X | Not Available | 69% | 99% |
| Plasminogen | CHEMBL1801 | P00747 | 4DUR | T89034 | 69% | 92% |
| Cysteinyl leukotriene receptor 2 | CHEMBL4330 | Q9NS75 | Not Available | T74238 | 68% | 98% |
| Platelet-derived growth factor receptor | CHEMBL2095189 | P09619 | 3MJG | T53524 | 68% | 72% |
| C-C chemokine receptor type 1 | CHEMBL2413 | P32246 | Not Available | T16016 | 68% | 89.5% |
| Cyclooxygenase-1 | CHEMBL221 | P23219 | 6Y3C | Not Available | 68% | 90% |
| Cystic fibrosis transmembrane conductance regulator | CHEMBL4051 | P13569 | 6MSM | T55654 | 68.1% | 96% |
| Adenosine A2b receptor | CHEMBL255 | P29275 | Not Available | T86679 | 68% | 99% |
| Histone deacetylase 11 | CHEMBL3310 | Q96DB2 | Not Available | Not Available | 68% | 89% |
| Melanin-concentrating hormone receptor 1 | CHEMBL344 | Q99705 | Not Available | T09572 | 67.6% | 92.5% |
| Dual specificity phosphatase Cdc25C | CHEMBL2378 | P30307 | 3OP3 | Not Available | 67% | 97% |
| Tyrosine-protein kinase receptor UFO | CHEMBL4895 | P30530 | 5U6B | T82383 | 67% | 91% |
| Transient receptor potential cation channel subfamily M member 8 | CHEMBL1075319 | Q7Z2W7 | Not Available | T41955 | 67% | 79.5% |
| Muscarinic acetylcholine receptor M1 | CHEMBL216 | P11229 | 6OIJ | T28893 | 66% | 94% |
| Nicotinamide phosphoribosyltransferase | CHEMBL1744525 | P43490 | 4LVF | T54582 | 65% | 96% |
| Beta-glucocerebrosidase | CHEMBL2179 | P04062 | 6TN1 | T84173 | 65.3% | 85% |
| Renin | CHEMBL286 | P00797 | 4AMT | T61622 | 65% | 97% |
| Phosphodiesterase 4B | CHEMBL275 | Q07343 | 4WZI | T10265 | 65% | 98% |
| Inhibitor of nuclear factor kappa B kinase beta subunit | CHEMBL1991 | O14920 | 4KIK | Not Available | 65% | 97% |
| Histone lysine demethylase PHF8 | CHEMBL1938212 | Q9UPP1 | 3KV4 | Not Available | 64% | 98% |
| Rho-associated protein kinase 1 | CHEMBL3231 | Q13464 | 3V8S | T51282 | 64% | 96% |
| Dipeptidyl peptidase VIII | CHEMBL4657 | Q6V1X1 | 6EOP | Not Available | 64% | 97% |
| Signal transducer and activator of transcription 1-alpha/beta | CHEMBL6101 | P42224 | 1YVL | T64205 | 64% | 73% |
| Pregnane X receptor | CHEMBL3401 | O75469 | 6TFI | T82702 | 64% | 95% |
| Sodium channel protein type IX alpha subunit | CHEMBL4296 | Q15858 | 6J8G | T12119 | 64% | 96% |
| Serine/threonine-protein kinase receptor R3 | CHEMBL5311 | P37023 | 3MY0 | T36959 | 64% | 83% |
| Methionine aminopeptidase 2 | CHEMBL3922 | P50579 | 1B6A | T75596 | 64% | 97% |
| Tyrosine-protein kinase ZAP-70 | CHEMBL2803 | P43403 | 2OZO | Not Available | 63% | 82.5% |
| Tyrosine-protein kinase CSK | CHEMBL2634 | P41240 | 1BYG | T64721 | 63% | 79% |
| PI3-kinase p110-beta subunit | CHEMBL3145 | P42338 | Not Available | T05031 | 63% | 99% |
| P2X purinoceptor 7 | CHEMBL4805 | Q99572 | Not Available | T63414 | 63% | 97.5% |
| Voltage-gated N-type calcium channel alpha-1B subunit | CHEMBL4478 | Q00975 | Not Available | T38338 | 63% | 97% |
| Tissue-type plasminogen activator | CHEMBL1873 | P00750 | 1RTF | T45299 | 62% | 93% |
| Cannabinoid CB1 receptor | CHEMBL218 | P21554 | 6N4B | Not Available | 61.9% | 97% |
| PI3-kinase p110-alpha/p85-alpha | CHEMBL2111367 | P27986 | 4JPS | T80276 | 62% | 94% |
| Serine/threonine-protein kinase MST2 | CHEMBL4708 | Q13188 | 6AO5 | Not Available | 62% | 72% |
| Dipeptidyl peptidase IX | CHEMBL4793 | Q86TI2 | 6EOR | Not Available | 60.8% | 97% |
| ATPase family AAA domain-containing protein 2 | CHEMBL2150837 | Q6PL18 | 5R4V | Not Available | 61% | 96% |
| Muscarinic acetylcholine receptor M5 | CHEMBL2035 | P08912 | 6OL9 | T79961 | 61% | 95% |
| Carboxylesterase 2 | CHEMBL3180 | O00748 | Not Available | Not Available | 60% | 90% |
| Protein-arginine N-methyltransferase 1 | CHEMBL5524 | Q99873 | 6NT2 | Not Available | 60% | 97% |
| Glutaminase kidney isoform, mitochondrial | CHEMBL2146302 | O94925 | 3UO9 | T86734 | 60% | 100% |
| Melanocortin receptor 5 | CHEMBL4608 | P33032 | Not Available | T95302 | 60% | 97% |
| Amine oxidase, copper containing | CHEMBL3437 | Q16853 | 2C10 | T69619 | 60% | 94% |
| Pyroglutamylated RFamide peptide receptor | CHEMBL5852 | Q96P65 | Not Available | Not Available | 60% | 85% |
| Growth factor receptor-bound protein 2 | CHEMBL3663 | P62993 | 1GRI | Not Available | 60% | 90% |
| Casein kinase II alpha/beta | CHEMBL3038477 | P67870 | 6TLS | T51565 | 60% | 99% |
| Ghrelin receptor | CHEMBL4616 | Q92847 | 6KO5 | T59604 | 59.1% | 92% |
| Calpain 1 | CHEMBL3891 | P07384 | 1ZCM | Not Available | 58.9% | 93% |
| Glycine transporter 1 | CHEMBL2337 | P48067 | 6ZBV | T69685 | 59% | 95% |
| Phosphodiesterase 11A | CHEMBL2717 | Q9HCR9 | Not Available | Not Available | 58% | 85% |
| Cytochrome P450 2A6 | CHEMBL5282 | P11509 | 2FDV | T06455 | 58% | 72% |
| Sodium channel protein type V alpha subunit | CHEMBL1980 | Q14524 | 6LQA | T39716 | 58% | 92.5% |
| Sphingosine 1-phosphate receptor Edg-8 | CHEMBL2274 | Q9H228 | Not Available | T50089 | 58% | 100% |
| Tissue factor pathway inhibitor | CHEMBL3713062 | P10646 | 5NMV | T78890 | 57% | 97% |
| Phosphodiesterase 4D | CHEMBL288 | Q08499 | 5WH6 | T02001 | 57% | 97.5% |
| Tyrosine-protein kinase FYN | CHEMBL1841 | P06241 | 2DQ7 | T17980 | 57% | 81% |
| T-cell protein-tyrosine phosphatase | CHEMBL3807 | P17706 | 1L8K | Not Available | 57% | 93% |
| Sodium channel protein type II alpha subunit | CHEMBL4187 | Q99250 | 6J8E | Not Available | 57% | 95.5% |
| Protein arginine N-methyltransferase 6 | CHEMBL1275221 | Q96LA8 | 4QQK | Not Available | 57% | 98% |
| PI3-kinase p110-delta subunit | CHEMBL3130 | O00329 | 6PYR | T67849 | 56% | 96% |
| ADAM10 | CHEMBL5028 | O14672 | 6BE6 | T31902 | 56% | 97.5% |
| G-protein coupled receptor 6 | CHEMBL3714130 | P46095 | Not Available | Not Available | 56% | 97% |
| Farnesyl diphosphate synthase | CHEMBL1782 | P14324 | 5CG5 | T86528 | 56% | 92% |
| Autotaxin | CHEMBL3691 | Q13822 | 4ZG6 | Not Available | 55% | 96% |
| Cyclin-dependent kinase 1/cyclin B | CHEMBL2094127 | P06493 | 6GU2 | T49898 | 55% | 96% |
| Histone deacetylase 4 | CHEMBL3524 | P56524 | 2VQM | T63816 | 55% | 93% |
| Egl nine homolog 1 | CHEMBL5697 | Q9GZT9 | 4BQY | Not Available | 55% | 93.4% |
| Arachidonate 5-lipoxygenase | CHEMBL215 | P09917 | 3V98 | Not Available | 55% | 93% |
| Integrin alpha-4/beta-1 | CHEMBL1907599 | P05556 | 3V4V | T97587 | 55% | 93% |
| Histone deacetylase 2 | CHEMBL1937 | Q92769 | 7KBG | T51191 | 55% | 95% |
| PI3-kinase p110-alpha subunit | CHEMBL4005 | P42336 | 4JPS | T80276 | 55% | 97% |
| Neuronal acetylcholine receptor; alpha3/beta4 | CHEMBL1907594 | P30926 | 6PV7 | T73724 | 55% | 97% |
| Transmembrane protease serine 6 | CHEMBL1795139 | Q8IU80 | Not Available | Not Available | 55% | 98% |
| NADPH oxidase 1 | CHEMBL1287628 | Q9Y5S8 | Not Available | Not Available | 54.5% | 95% |
| C-C chemokine receptor type 2 | CHEMBL4015 | P41597 | 5T1A | T89988 | 54% | 99% |
| 15-hydroxyprostaglandin dehydrogenase [NAD+] | CHEMBL1293255 | P15428 | 2GDZ | Not Available | 54% | 84% |
| Histone deacetylase 1 | CHEMBL325 | Q13547 | 4BKX | T68547 | 54% | 96% |
| Lysine-specific demethylase 5C | CHEMBL2163176 | P41229 | 5FWJ | Not Available | 54% | 93% |
| Toll-like receptor 4 | CHEMBL5255 | O00206 | 4G8A | T81443 | 54% | 92.5% |
| DCN1-like protein 1 | CHEMBL4105838 | Q96GG9 | 6BG3 | Not Available | 54% | 95% |
| Protein kinase N2 | CHEMBL3032 | Q16513 | 4CRS | Not Available | 53% | 78% |
| Glutamate carboxypeptidase II | CHEMBL1892 | Q04609 | 5O5T | Not Available | 53% | 97.5% |
| Aurora kinase B/Inner centromere protein | CHEMBL3430907 | Q96GD4 | 6YIH | T46781 | 53% | 97.5% |
| Phosphodiesterase 5A | CHEMBL1827 | O76074 | 3BJC | T07663 | 53% | 100% |
| Xanthine dehydrogenase | CHEMBL1929 | P47989 | 2E1Q | T40954 | 53% | 96% |
| Trypsin I | CHEMBL209 | P07477 | 2RA3 | T27602 | 53% | 90% |
| Fatty acid synthase | CHEMBL4158 | P49327 | 3HHD | T16514 | 52% | 82.5% |
| Ectonucleotide pyrophosphatase/phosphodiesterase family member 1 | CHEMBL5925 | P22413 | 6WFJ | Not Available | 52% | 92% |
| Vitamin K-dependent protein C | CHEMBL4444 | P04070 | 6M3B | T24836 | 52% | 94% |
| D-amino-acid oxidase | CHEMBL5485 | P14920 | 3ZNN | T33124 | 52% | 97% |
| Dopamine D1 receptor | CHEMBL2056 | P21728 | 7JVP | Not Available | 51.6% | 91% |
| Protein farnesyltransferase | CHEMBL2094108 | P49354 | 2H6F | T86428 | 51% | 98% |
| Plasminogen activator inhibitor-1 | CHEMBL3475 | P05121 | 3CVM | T15556 | 51% | 83% |
| Potassium channel subfamily K member 9 | CHEMBL2321614 | Q9NPC2 | 6GHP | Not Available | 51% | 80% |
| Carnitine O-palmitoyltransferase 1, muscle isoform | CHEMBL2216739 | Q92523 | Not Available | T75888 | 51% | 88% |
| Multidrug resistance-associated protein 1 | CHEMBL3004 | P33527 | 4C3Z | T11288 | 51% | 96% |
| Transthyretin | CHEMBL3194 | P02766 | 6SUG | T86462 | 50% | 91% |
| Receptor-interacting serine/threonine-protein kinase 3 | CHEMBL1795199 | Q9Y572 | 7DA4 | Not Available | 50% | 80% |
| Histone deacetylase 10 | CHEMBL5103 | Q969S8 | Not Available | T94324 | 50% | 90% |
| Macrophage migration inhibitory factor | CHEMBL2085 | P14174 | 6B1K | T39977 | 50% | 81% |

**Table S5. Predicted targets of Sertraline**

| **Target Name** | **ChEMBL-ID** | **UniProt ID** | **PDB Visualization** | **TTD ID** | **Probability** | **Accuracy** |
| --- | --- | --- | --- | --- | --- | --- |
| DNA-(apurinic or apyrimidinic site) lyase | CHEMBL5619 | P27695 | 6BOW | T13348 | 97% | 91% |
| HERG | CHEMBL240 | Q12809 | 5VA1 | T20251 | 94% | 90% |
| Free fatty acid receptor 1 | CHEMBL4422 | O14842 | 5TZR | Not Available | 91% | 93% |
| Cyclooxygenase-1 | CHEMBL221 | P23219 | 6Y3C | Not Available | 90% | 90% |
| LSD1/CoREST complex | CHEMBL3137262 | O60341 | 5L3D | Not Available | 90% | 97% |
| Cathepsin D | CHEMBL2581 | P07339 | 4OD9 | T67102 | 89% | 99% |
| Serotonin 2c (5-HT2c) receptor | CHEMBL225 | P28335 | 6BQH | T83813 | 89% | 90% |
| Dual specificity protein kinase CLK4 | CHEMBL4203 | Q9HAZ1 | 6FYV | Not Available | 88% | 94% |
| Signal transducer and activator of transcription 3 | CHEMBL4026 | P40763 | 6QHD | T29130 | 87% | 83% |
| Lysosomal Pro-X carboxypeptidase | CHEMBL2335 | P42785 | 3N2Z | Not Available | 85% | 100% |
| Nuclear factor NF-kappa-B p105 subunit | CHEMBL3251 | P19838 | 1SVC | Not Available | 85% | 96% |
| Voltage-gated N-type calcium channel alpha-1B subunit | CHEMBL4478 | Q00975 | Not Available | T38338 | 85% | 97% |
| Transcription intermediary factor 1-alpha | CHEMBL3108638 | O15164 | 4YBM | Not Available | 84% | 96% |
| Histone deacetylase 7 | CHEMBL2716 | Q8WUI4 | 3C10 | Not Available | 84% | 89% |
| Tyrosine-protein kinase TYK2 | CHEMBL3553 | P29597 | 4OLI | T78932 | 83% | 97% |
| Serotonin 7 (5-HT7) receptor | CHEMBL3155 | P34969 | Not Available | T79062 | 82% | 91% |
| NT-3 growth factor receptor | CHEMBL5608 | Q16288 | 6KZD | Not Available | 82% | 96% |
| Excitatory amino acid transporter 1 | CHEMBL3085 | P43003 | 5LM4 | Not Available | 81% | 95% |
| Voltage-gated potassium channel subunit Kv1.5 | CHEMBL4306 | P22460 | Not Available | T17569 | 81.1% | 94% |
| Sodium/hydrogen exchanger 1 | CHEMBL2781 | P19634 | 7DSX | T82028 | 80% | 90% |
| Cyclin-dependent kinase 5/CDK5 activator 1 | CHEMBL1907600 | Q00535 | 1UNL | T20973 | 80% | 93% |
| Proteasome component C5 | CHEMBL4208 | P20618 | 6KWY | Not Available | 80% | 90% |
| Glutathione S-transferase Pi | CHEMBL3902 | P09211 | 5J41 | T21669 | 79% | 94% |
| Neurokinin 2 receptor | CHEMBL2327 | P21452 | Not Available | T52790 | 79% | 99% |
| Neuronal acetylcholine receptor; alpha3/beta4 | CHEMBL1907594 | P30926 | 6PV7 | T73724 | 78% | 97% |
| Lysine-specific histone demethylase 1 | CHEMBL6136 | O60341 | 5L3D | Not Available | 78% | 96% |
| Metabotropic glutamate receptor 5 | CHEMBL3227 | P41594 | 6N4X | Not Available | 77% | 96% |
| Cyclin-dependent kinase 1/cyclin B | CHEMBL2094127 | P06493 | 6GU2 | T49898 | 77% | 96% |
| Serine/threonine-protein kinase TAO1 | CHEMBL5261 | Q7L7X3 | Not Available | Not Available | 75% | 89% |
| Tyrosine-protein kinase ZAP-70 | CHEMBL2803 | P43403 | 2OZO | Not Available | 75% | 82.5% |
| Protein kinase N1 | CHEMBL3384 | Q16512 | 4OTH | Not Available | 75% | 81% |
| C5a anaphylatoxin chemotactic receptor | CHEMBL2373 | P21730 | 6C1R | T15439 | 75% | 93% |
| Tyrosyl-DNA phosphodiesterase 1 | CHEMBL1075138 | Q9NUW8 | 6N0D | Not Available | 75% | 71% |
| Pregnane X receptor | CHEMBL3401 | O75469 | 6TFI | T82702 | 74% | 95% |
| Ribosomal protein S6 kinase alpha 1 | CHEMBL2553 | Q15418 | 2Z7Q | Not Available | 73% | 85% |
| ADAM10 | CHEMBL5028 | O14672 | 6BE6 | T31902 | 73% | 97.5% |
| Cathepsin S | CHEMBL2954 | P25774 | 2C0Y | T68290 | 73% | 95.6% |
| Angiotensin-converting enzyme | CHEMBL1808 | P12821 | 5AMB | T82577 | 73% | 93% |
| Glutaminase kidney isoform, mitochondrial | CHEMBL2146302 | O94925 | 3UO9 | T86734 | 73% | 100% |
| Serine/threonine-protein kinase TAO3 | CHEMBL5701 | Q9H2K8 | 6BDN | Not Available | 73% | 97% |
| Histamine H3 receptor | CHEMBL264 | Q9Y5N1 | Not Available | Not Available | 73% | 91% |
| Kruppel-like factor 5 | CHEMBL1293249 | Q13887 | Not Available | Not Available | 72% | 86% |
| G-protein coupled receptor 55 | CHEMBL1075322 | Q9Y2T6 | Not Available | T87670 | 72% | 78% |
| Melanocortin receptor 4 | CHEMBL259 | P32245 | 7AUE | T72458 | 70% | 95% |
| Beta-glucuronidase | CHEMBL2728 | P08236 | 3HN3 | T96413 | 70% | 78% |
| Voltage-gated T-type calcium channel alpha-1H subunit | CHEMBL1859 | O95180 | Not Available | T54644 | 70% | 99% |
| 5'-nucleotidase | CHEMBL5957 | P21589 | 6TVE | Not Available | 70% | 98% |
| Tyrosine-protein kinase ITK/TSK | CHEMBL2959 | Q08881 | 4HCU | T91761 | 69% | 95% |
| Dopamine D1 receptor | CHEMBL2056 | P21728 | 7JVP | Not Available | 69% | 91% |
| Histone deacetylase 8 | CHEMBL3192 | Q9BY41 | 5VI6 | T28887 | 69% | 94% |
| Toll-like receptor 8 | CHEMBL5805 | Q9NR97 | 3WN4 | T48703 | 68% | 96% |
| Dual specificity phosphatase Cdc25B | CHEMBL4804 | P30305 | 1QB0 | Not Available | 67% | 79.5% |
| Dipeptidyl peptidase IX | CHEMBL4793 | Q86TI2 | 6EOR | Not Available | 67% | 97% |
| dUTP pyrophosphatase | CHEMBL5203 | P33316 | 3ARA | T39054 | 66% | 99% |
| Glucose transporter | CHEMBL2535 | P11166 | 6THA | Not Available | 66% | 99% |
| Dual specificity phosphatase Cdc25C | CHEMBL2378 | P30307 | 3OP3 | Not Available | 66% | 97% |
| Ephrin type-A receptor 4 | CHEMBL3988 | P54764 | 4M4P | T70234 | 66.1% | 74% |
| Transmembrane protease serine 6 | CHEMBL1795139 | Q8IU80 | Not Available | Not Available | 66% | 98% |
| Nitric oxide synthase, inducible | CHEMBL4481 | P35228 | 3E7G | T02703 | 66% | 94.8% |
| DCN1-like protein 1 | CHEMBL4105838 | Q96GG9 | 6BG3 | Not Available | 65% | 95% |
| Integrin alpha-5/beta-1 | CHEMBL2095226 | P05556 | 7NWL | T01851 | 65% | 96% |
| Egl nine homolog 1 | CHEMBL5697 | Q9GZT9 | 4BQY | Not Available | 64% | 93.4% |
| Integrin alpha-4/beta-1 | CHEMBL1907599 | P05556 | 3V4V | T97587 | 64% | 93% |
| GABA-A receptor; alpha-1/beta-2/gamma-2 | CHEMBL2095172 | P14867 | 6X3T | T51487 | 64% | 93% |
| Mineralocorticoid receptor | CHEMBL1994 | P08235 | 4PF3 | Not Available | 64% | 100% |
| C-C chemokine receptor type 2 | CHEMBL4015 | P41597 | 5T1A | T89988 | 64% | 99% |
| Ephrin type-A receptor 2 | CHEMBL2068 | P29317 | 3FL7 | T57278 | 64% | 97% |
| Coagulation factor XIII | CHEMBL4530 | P00488 | 4KTY | Not Available | 64% | 96% |
| Thymidine phosphorylase | CHEMBL3106 | P19971 | 1UOU | T59929 | 63% | 77% |
| Tyrosine-protein kinase receptor TYRO3 | CHEMBL5314 | Q06418 | 1RHF | Not Available | 62.6% | 96% |
| Vitamin K-dependent protein C | CHEMBL4444 | P04070 | 6M3B | T24836 | 62% | 94% |
| Ephrin type-B receptor 4 | CHEMBL5147 | P54760 | 6FNK | T49507 | 62% | 97% |
| Muscarinic acetylcholine receptor M1 | CHEMBL216 | P11229 | 6OIJ | T28893 | 62% | 94% |
| Aurora kinase B/Inner centromere protein | CHEMBL3430907 | Q96GD4 | 6YIH | T46781 | 62% | 97.5% |
| Tyrosine-protein kinase SRC | CHEMBL267 | P12931 | 2H8H | T85943 | 62% | 96% |
| Metabotropic glutamate receptor 4 | CHEMBL2736 | Q14833 | 7E9H | Not Available | 61% | 98% |
| Tyrosine-protein kinase YES | CHEMBL2073 | P07947 | 2HDA | Not Available | 61% | 83% |
| Glycine receptor subunit alpha-1 | CHEMBL5845 | P23415 | 4X5T | T50269 | 61% | 91% |
| Fatty acid binding protein adipocyte | CHEMBL2083 | P15090 | 3P6D | Not Available | 61% | 96% |
| Adenosine A1 receptor | CHEMBL226 | P30542 | 5N2S | T92072 | 60% | 96% |
| Glycogen synthase kinase-3 alpha | CHEMBL2850 | P49840 | Not Available | Not Available | 60% | 89% |
| Tyrosine-protein kinase FYN | CHEMBL1841 | P06241 | 2DQ7 | T17980 | 60% | 81% |
| c-Jun N-terminal kinase 3 | CHEMBL2637 | P53779 | 7KSI | Not Available | 60% | 93% |
| Histone-arginine methyltransferase CARM1 | CHEMBL5406 | Q86X55 | 6DVR | T12837 | 60% | 94% |
| Galectin-3 | CHEMBL4531 | P17931 | 6FOF | T72038 | 60% | 96.9% |
| Stimulator of interferon genes protein | CHEMBL4523377 | Q86WV6 | 6NT5 | Not Available | 59% | 95% |
| c-Jun N-terminal kinase 2 | CHEMBL4179 | P45984 | 3E7O | Not Available | 59% | 91% |
| Acetylcholine receptor; alpha1/beta1/delta/gamma | CHEMBL1907588 | P02708 | 5HBT | T04689 | 59% | 98% |
| Tissue factor pathway inhibitor | CHEMBL3713062 | P10646 | 5NMV | T78890 | 59% | 97% |
| Cyclin-dependent kinase 2/cyclin E1 | CHEMBL1907605 | P24864 | 1W98 | T70176 | 59% | 93% |
| Free fatty acid receptor 2 | CHEMBL5493 | O15552 | Not Available | T28213 | 59% | 93% |
| Nuclear receptor ROR-beta | CHEMBL3091268 | Q92753 | Not Available | Not Available | 58% | 95.5% |
| Histone deacetylase 5 | CHEMBL2563 | Q9UQL6 | 5UWI | Not Available | 58.4% | 90% |
| Dihydroorotate dehydrogenase | CHEMBL1966 | Q02127 | 6FMD | T99009 | 58% | 96% |
| Cyclin-dependent kinase 5 | CHEMBL4036 | Q00535 | 4AU8 | T20973 | 58% | 79% |
| Cannabinoid CB2 receptor | CHEMBL253 | P34972 | 6KPF | Not Available | 58% | 97% |
| Formyl peptide receptor 1 | CHEMBL3359 | P21462 | Not Available | T87831 | 58% | 94% |
| Ephrin type-B receptor 2 | CHEMBL3290 | P29323 | 3ZFM | T73756 | 58% | 78% |
| Histone lysine demethylase PHF8 | CHEMBL1938212 | Q9UPP1 | 3KV4 | Not Available | 57% | 98% |
| Nuclear factor erythroid 2-related factor 2 | CHEMBL1075094 | Q16236 | 2FLU | Not Available | 57% | 96% |
| G-protein coupled receptor 35 | CHEMBL1293267 | Q9HC97 | Not Available | Not Available | 57% | 89% |
| Hydroxycarboxylic acid receptor 2 | CHEMBL3785 | Q8TDS4 | Not Available | T88185 | 56% | 94% |
| Sodium/glucose cotransporter 1 | CHEMBL4979 | P13866 | Not Available | Not Available | 56% | 98% |
| Telomerase reverse transcriptase | CHEMBL2916 | O14746 | 7BG9 | T86052 | 56% | 90% |
| Protein kinase C zeta | CHEMBL3438 | Q05513 | Not Available | T59190 | 56% | 88% |
| Glycine transporter 1 | CHEMBL2337 | P48067 | 6ZBV | T69685 | 56% | 95% |
| Serine/threonine-protein kinase ULK1 | CHEMBL6006 | O75385 | 4WNO | Not Available | 56% | 76% |
| Glutamate carboxypeptidase II | CHEMBL1892 | Q04609 | 5O5T | Not Available | 55.5% | 97.5% |
| Thromboxane A2 receptor | CHEMBL2069 | P21731 | Not Available | T76198 | 55% | 93% |
| Glycine transporter 2 | CHEMBL3060 | Q9Y345 | Not Available | Not Available | 55% | 99% |
| Acyl-CoA desaturase | CHEMBL5555 | O00767 | 4ZYO | T10897 | 55.1% | 97.5% |
| Lysine-specific demethylase 4A | CHEMBL5896 | O75164 | 6G5X | Not Available | 55% | 99% |
| Serine/threonine-protein kinase receptor R3 | CHEMBL5311 | P37023 | 3MY0 | T36959 | 55% | 83% |
| Sodium channel protein type IX alpha subunit | CHEMBL4296 | Q15858 | 6J8G | T12119 | 55% | 96% |
| Toll-like receptor 4 | CHEMBL5255 | O00206 | 4G8A | T81443 | 55% | 92.5% |
| Plasminogen activator inhibitor-1 | CHEMBL3475 | P05121 | 3CVM | T15556 | 55% | 83% |
| Lymphocyte differentiation antigen CD38 | CHEMBL4660 | P28907 | 3F6Y | T10877 | 55% | 95% |
| Protein-tyrosine phosphatase 1B | CHEMBL335 | P18031 | 5QGF | Not Available | 55% | 95% |
| Pyroglutamylated RFamide peptide receptor | CHEMBL5852 | Q96P65 | Not Available | Not Available | 54% | 85% |
| Acyl coenzyme A:cholesterol acyltransferase 1 | CHEMBL2782 | P35610 | 6P2J | Not Available | 54% | 92% |
| Bromodomain-containing protein 4 | CHEMBL1163125 | O60885 | 6DNE | T40556 | 54% | 97% |
| Rho-associated protein kinase 1 | CHEMBL3231 | Q13464 | 3V8S | T51282 | 54% | 96% |
| Lipoxin A4 receptor | CHEMBL4227 | P25090 | 6OMM | Not Available | 54% | 100% |
| Cyclin-dependent kinase 2 | CHEMBL301 | P24941 | 5LMK | T70176 | 54% | 91% |
| Carbonic anhydrase XIV | CHEMBL3510 | Q9ULX7 | 5CJF | Not Available | 54% | 95% |
| Carnitine palmitoyltransferase 2 | CHEMBL3238 | P23786 | Not Available | Not Available | 53% | 94% |
| Cathepsin B | CHEMBL4072 | P07858 | 3PBH | T61746 | 53% | 94% |
| P2X purinoceptor 4 | CHEMBL2104 | Q99571 | Not Available | T60330 | 53% | 97.5% |
| Prolyl endopeptidase | CHEMBL3202 | P48147 | 3DDU | T86161 | 53% | 91% |
| TRAF2- and NCK-interacting kinase | CHEMBL4527 | Q9UKE5 | 2X7F | Not Available | 53% | 70% |
| Dual specificity protein phosphatase 3 | CHEMBL2635 | P51452 | 3F81 | Not Available | 53% | 94% |
| Cystic fibrosis transmembrane conductance regulator | CHEMBL4051 | P13569 | 6MSM | T55654 | 53% | 96% |
| Monoamine oxidase B | CHEMBL2039 | P27338 | 1S3E | Not Available | 52% | 93% |
| Dual specificity protein kinase CLK2 | CHEMBL4225 | P49760 | 6KHE | Not Available | 52% | 81% |
| Stem cell growth factor receptor | CHEMBL1936 | P10721 | 2EC8 | T57700 | 52% | 84% |
| Bromodomain-containing protein 2 | CHEMBL1293289 | P25440 | 6DDI | T86399 | 52% | 86% |
| Neurokinin 1 receptor | CHEMBL249 | P25103 | 6HLP | T47094 | 51.6% | 99% |
| Aldo-keto reductase family 1 member C2 | CHEMBL5847 | P52895 | 4XO6 | Not Available | 52% | 92.5% |
| Tyrosine-protein kinase FGR | CHEMBL4454 | P09769 | Not Available | Not Available | 51% | 77% |
| Adenosylhomocysteinase | CHEMBL2664 | P23526 | 1LI4 | Not Available | 51% | 87% |
| Glutamate NMDA receptor; GRIN1/GRIN2B | CHEMBL1907603 | Q05586 | 5EWM | Not Available | 51% | 96% |
| ATPase family AAA domain-containing protein 2 | CHEMBL2150837 | Q6PL18 | 5R4V | Not Available | 51% | 96% |
| Tyrosine-protein kinase BRK | CHEMBL4601 | Q13882 | 6CZ4 | T73694 | 51% | 76% |
| Mitogen-activated protein kinase 7 | CHEMBL5332 | Q13164 | 4IC7 | Not Available | 51% | 93% |
| Cytochrome P450 3A4 | CHEMBL340 | P08684 | 5VCC | T37848 | 51% | 91% |
| Sphingosine 1-phosphate receptor Edg-6 | CHEMBL3230 | O95977 | Not Available | Not Available | 51% | 94% |
| Excitatory amino acid transporter 3 | CHEMBL2721 | P43005 | 6X2L | Not Available | 50% | 93.5% |
| Ephrin type-B receptor 3 | CHEMBL4901 | P54753 | 5L6O | Not Available | 50% | 87.5% |

**Table S6. Co-upregulated and co-downregulated genes upon citalopram and fluvoxamine treatment**

| **Gene Symbol** | **Citalopram vs DMSO** | | **Fluvoxamine vs DMSO** | |
| --- | --- | --- | --- | --- |
|  | **Log_2_FC** | **P-value** | **Log_2_FC** | **P-value** |
| *ACAT2* | 1.297475 | 7.63E-21 | 1.138184 | 1.19E-15 |
| *ACSS2* | 1.838341 | 4.26E-33 | 1.523079 | 3.06E-22 |
| *ACSS3* | 2.66498 | 0.006801 | 2.992956 | 0.008359 |
| *ADM2* | 1.335821 | 5.85E-07 | 1.09367 | 0.000111 |
| *ALDOC* | 1.771214 | 4.99E-19 | 1.33256 | 7.28E-10 |
| *AMIGO3* | 4.946292 | 4.06E-59 | 3.39966 | 0.004498 |
| *B4GALNT2* | 3.302333 | 0.032989 | 3.847686 | 0.003029 |
| *BCL2L2-PABPN1* | 1.872487 | 2.74E-05 | 2.133471 | 6.36E-18 |
| *C5orf24* | 1.213474 | 0.017326 | 1.2457 | 0.000234 |
| *C6orf223* | 1.61066 | 1.65E-11 | 1.21786 | 3.87E-06 |
| *CDK3* | 1.288011 | 0.001547 | 1.278852 | 0.006537 |
| *CYP2J2* | 3.150924 | 0.001162 | 2.62518 | 0.023485 |
| *CYP51A1P2* | 1.219978 | 9.53E-08 | 1.00113 | 2.32E-05 |
| *DHCR7* | 1.387527 | 6.44E-14 | 1.094664 | 1.04E-08 |
| *FABP3* | 1.927367 | 8.00E-12 | 1.319288 | 1.06E-05 |
| *FABP6* | 1.381638 | 0.008822 | 1.157805 | 0.043661 |
| *FADS2* | 1.272561 | 6.90E-14 | 1.052974 | 1.66E-09 |
| *FDPSP1* | 1.213439 | 0.022446 | 1.215093 | 0.01885 |
| *FFAR2* | 1.125356 | 0.026119 | 1.060908 | 0.035908 |
| *FGF9* | 1.275771 | 0.004062 | 1.085128 | 0.013809 |
| *FN3K* | 1.422779 | 3.39E-07 | 1.039572 | 0.000253 |
| *GYG2* | 1.170937 | 0.014177 | 1.278076 | 0.001539 |
| *HMGCS1* | 1.126997 | 1.10E-13 | 1.121823 | 4.64E-13 |
| *HSD17B7* | 1.434542 | 1.00E-12 | 1.202314 | 8.72E-09 |
| *HSP90AB2P* | 1.64764 | 0.001424 | 1.89192 | 0.000378 |
| *HVCN1* | 1.028281 | 0.006052 | 1.18345 | 0.004543 |
| *IGFL2-AS1* | 1.339004 | 4.07E-10 | 1.168224 | 2.87E-08 |
| *INSIG1* | 1.63264 | 2.31E-31 | 1.268535 | 2.48E-11 |
| *JRK* | 1.118447 | 0.007332 | 1.515678 | 4.86E-05 |
| *KIAA0408* | 1.489856 | 1.51E-05 | 1.227988 | 0.000447 |
| *KLHL38* | 2.030621 | 1.14E-06 | 1.454726 | 0.000738 |
| *KLRG1* | 1.16341 | 0.009143 | 1.014483 | 0.028434 |
| *LINC00672* | 1.494105 | 0.00436 | 1.292449 | 0.015881 |
| *LRRC37A4P* | 3.080784 | 0.044586 | 4.028895 | 0.004097 |
| *LSS* | 1.435916 | 3.60E-15 | 1.289307 | 2.49E-12 |
| *MADCAM1* | 1.226846 | 0.007163 | 1.325195 | 0.013572 |
| *MIR378D2HG* | 3.710529 | 0.004627 | 3.63119 | 0.032266 |
| *MIR4707* | 3.004877 | 0.025398 | 2.819577 | 0.035939 |
| *MLXIPL* | 1.307363 | 1.38E-07 | 1.012192 | 7.50E-05 |
| *MSMO1* | 1.448921 | 8.35E-25 | 1.452819 | 6.95E-22 |
| *MVD* | 1.488163 | 7.87E-20 | 1.149368 | 3.50E-12 |
| *NFATC4* | 2.104328 | 0.002703 | 1.822253 | 0.038389 |
| *NLRP1* | 1.970803 | 0.005987 | 1.661843 | 0.026873 |
| *NPR1* | 2.200211 | 4.52E-07 | 1.534823 | 0.002826 |
| *NT5DC4* | 1.785474 | 0.009457 | 1.543452 | 0.043329 |
| *P2RY8* | 3.648833 | 0.000437 | 3.629255 | 5.27E-05 |
| *PCSK9* | 1.667267 | 6.51E-29 | 1.375372 | 2.91E-18 |
| *PDE5A* | 1.951077 | 0.000285 | 1.938597 | 0.005504 |
| *PIK3IP1* | 1.045788 | 0.006815 | 1.126503 | 0.006276 |
| *PLA2G3* | 4.116991 | 1.08E-13 | 3.531567 | 1.18E-08 |
| *PLGLB2* | 3.729847 | 0.004135 | 3.544142 | 0.016191 |
| *PNPLA3* | 1.347604 | 1.75E-11 | 1.137199 | 5.05E-08 |
| *PON1* | 1.649835 | 0.00367 | 1.56643 | 0.006836 |
| *PRDM16* | 1.246707 | 1.10E-05 | 1.107249 | 0.001778 |
| *PRSS8* | 2.204936 | 0.002703 | 1.867066 | 0.049168 |
| *RAB26* | 1.248531 | 8.00E-06 | 1.007028 | 0.000848 |
| *RBM12* | 1.119348 | 0.002675 | 1.09694 | 0.019782 |
| *RPS6KA2* | 1.38581 | 0.001584 | 1.044369 | 0.023348 |
| *SARDH* | 2.014309 | 0.001747 | 1.614096 | 0.029906 |
| *SARM1* | 1.325209 | 0.006186 | 1.272589 | 0.014837 |
| *SC5D* | 1.141638 | 1.21E-05 | 1.605363 | 1.10E-16 |
| *SLC25A18* | 1.928377 | 3.17E-05 | 1.242732 | 0.029185 |
| *SLC2A10* | 1.333912 | 0.040639 | 1.439823 | 0.029086 |
| *SMIM10L2B* | 1.244064 | 0.002624 | 1.390116 | 0.000531 |
| *SQLE* | 1.345438 | 2.37E-14 | 1.271531 | 2.20E-12 |
| *SRP54-AS1* | 1.120044 | 0.009986 | 1.213439 | 0.003134 |
| *ST6GALNAC2* | 3.379204 | 2.39E-08 | 2.241659 | 0.011516 |
| *SYNE4* | 2.694202 | 0.000155 | 2.141846 | 0.027394 |
| *TCF19* | 5.076841 | 1.02E-07 | 5.420236 | 4.07E-08 |
| *TESK2* | 1.592969 | 0.000825 | 2.055373 | 2.11E-05 |
| *TM7SF2* | 1.854629 | 4.53E-15 | 1.551105 | 5.48E-11 |
| *TMOD2* | 1.201566 | 0.03874 | 1.430072 | 0.00807 |
| *TOMM40L* | 1.236191 | 8.68E-07 | 1.377145 | 1.41E-08 |
| *TP53INP1* | 1.270638 | 2.10E-10 | 1.068421 | 1.20E-07 |
| *TP53INP2* | 1.45889 | 9.31E-17 | 1.059715 | 2.35E-08 |
| *TPSP2* | 2.107054 | 2.07E-11 | 1.865981 | 2.51E-07 |
| *UGT1A4* | 4.691135 | 5.99E-95 | 2.001022 | 0.000868 |
| *WFDC21P* | 1.410952 | 0.01375 | 1.600443 | 0.005335 |
| *ZMYND15* | 1.007366 | 0.026842 | 1.156932 | 0.005882 |
| *ZNF497* | 1.784669 | 7.34E-09 | 1.275406 | 9.78E-05 |
| *ABCA1* | -1.02807 | 2.85E-10 | -1.11083 | 2.79E-10 |
| *ABCG1* | -1.66231 | 1.09E-05 | -1.45588 | 3.96E-05 |
| *BMS1P4* | -2.16235 | 0.010499 | -2.31824 | 1.44E-07 |
| *BMS1P9* | -3.09283 | 0.003067 | -2.0687 | 0.035782 |
| *CA4* | -1.42216 | 0.03512 | -2.2633 | 0.010159 |
| *CCDC140* | -1.94842 | 0.003218 | -1.39062 | 0.049407 |
| *CLCN1* | -1.20601 | 0.045477 | -2.06993 | 0.0024 |
| *CNBD1* | -3.45384 | 0.011397 | -3.42899 | 0.012863 |
| *CSF2* | -1.37972 | 0.011997 | -1.35703 | 0.014523 |
| *EEF1A1P45* | -2.36362 | 0.044126 | -2.39258 | 0.048005 |
| *EGOT* | -1.10526 | 0.036805 | -1.53748 | 0.008995 |
| *FMC1-LUC7L2* | -1.27986 | 0.010124 | -1.5325 | 0.002966 |
| *GBP5* | -1.19049 | 0.044421 | -1.81228 | 0.003284 |
| *GNA15* | -1.98463 | 0.030641 | -1.9861 | 0.017984 |
| *HACD2* | -1.17542 | 0.000679 | -1.95164 | 2.63E-42 |
| *ISLR2* | -2.0081 | 0.022262 | -1.98358 | 0.024914 |
| *ITGBL1* | -1.63272 | 0.041465 | -1.6424 | 0.044518 |
| *KRT18P59* | -2.20333 | 0.025584 | -2.18933 | 0.025054 |
| *MARCHF4* | -1.20679 | 0.049555 | -1.30584 | 0.030796 |
| *MTAP* | -1.48393 | 1.35E-19 | -1.15523 | 5.40E-12 |
| *MYL12-AS1* | -1.2861 | 0.036133 | -1.32249 | 0.035131 |
| *PDXDC2P* | -1.2851 | 0.015054 | -1.02316 | 0.008073 |
| *PIGW* | -1.91598 | 0.027169 | -1.92377 | 0.032857 |
| *PPP1R35-AS1* | -6.67882 | 2.87E-07 | -5.72984 | 2.14E-06 |
| *RN7SL809P* | -1.73036 | 0.03624 | -1.51489 | 0.032139 |
| *SCG2* | -1.45394 | 0.001082 | -1.55801 | 0.005468 |
| *SERPINB5* | -1.25646 | 0.002867 | -1.09721 | 0.010133 |
| *SLIT2* | -1.03496 | 0.005179 | -1.13315 | 0.003493 |
| *SNORA58B* | -3.21472 | 0.017015 | -3.18406 | 0.017875 |
| *TMEM164* | -1.80868 | 4.58E-07 | -2.25417 | 1.02E-10 |
| *TOMM6* | -5.35805 | 7.68E-07 | -6.88965 | 3.85E-09 |
| *VEZF1P1* | -1.96556 | 0.023647 | -1.43664 | 0.04975 |
| *WNT2B* | -1.03972 | 0.016969 | -1.00375 | 0.020678 |
| *ZNF791* | -3.04709 | 6.22E-25 | -2.0803 | 1.10E-14 |

**Table S7.** **The SSRI-β-catenin binding energies of the top-1 docked models**

| Compound | Binding energy (kcal/mol) |
| --- | --- |
| Citalopram | -4.56 |
| Fluoxetine | -4.36 |
| Fluvoxamine | -3.49 |
| Paroxetine | -5.13 |
| Sertraline | -5.92 |

**Table S8. Sequences for shRNA and sgRNA in this study**

| **Oligonucleotides** | **Sequences** |
| --- | --- |
| sh*SLC6A4*-#1 | CACCGCGCTATACTACCTCATCTCCCGAAGGAGATGAGGTAGTATAGCGC |
| sh*SLC6A4*-#2 | CACCGCCTCCTCTTCATCACGTATGCGAACATACGTGATGAAGAGGAGGC |
| sh*Slc6a4* | CACCGCCTCCTACTATAACACCATCCGAAGATGGTGTTATAGTAGGAGGC |
| sh*SLC2A1*-#1 | CACCGGAATTCAATGCTGATGATGACGAATCATCATCAGCATTGAATTCC |
| sh*SLC2A1*-#2 | CACCGCTACCCTGGATGTCCTATCTCGAAAGATAGGACATCCAGGGTAGC |
| *SLC6A4* sgRNA-#1 | GAGTCCGGGCAAATATCCAA |
| *SLC6A4* sgRNA-#2 | CATATGTTACCAGAATGGAG |

**Table S9. The sequences for primers used in this study**

| **Gene** | **Forward primer (5’-3’)** | **Reverse primer (5’-3’)** |
| --- | --- | --- |
| *SLC6A4* | ACGGAGTTCTACAGAAGGTTGT | ATAGAGTGCCGTGTGTCATCT |
| *Slc6a4* | TATCCAATGGGTACTCCGCAG | CCGTTCCCCTTGGTGAATCT |
| *SLC2A1* | ATTGGCTCCGGTATCGTCAAC | GCTCAGATAGGACATCCAGGGTA |
| *HK2* | TTGACCAGGAGATTGACATGGG | CAACCGCATCAGGACCTCA |
| *PKM2* | ATAACGCCTACATGGAAAAGTGT | TAAGCCCATCATCCACGTAGA |
| *LDHA* | ATGGCAACTCTAAAGGATCAGC | CCAACCCCAACAACTGTAATCT |
| *GAPDH* | GGAGCGAGATCCCTCCAAAAT | GGCTGTTGTCATACTTCTCATGG |
| *Gapdh* | AGGTCGGTGTGAACGGATTTG | TGTAGACCATGTAGTTGAGGTCA |
